# Heterologous expression and optimization of fermentation conditions for recombinant ikarugamycin production

**DOI:** 10.1101/2024.10.13.616080

**Authors:** Julia K. Evers, Anna Glöckle, Monique Wiegand, Sebastian Schuler, Manuel Einsiedler, Tobias A. M. Gulder

**Author notes:** Correspondence: Tobias A. M. Gulder,.

## Abstract

Ikarugamycin is a member of the natural product family of the polycyclic tetramate macrolactams (PoTeMs). The compound exhibits a diverse range of biological activities, including antimicrobial, antiprotozoal, anti-leukemic, and anti-inflammatory properties. In addition, it interferes with several crucial cellular functions, such as oxidized low-density lipoprotein uptake in macrophages, Nef-induced CD4 cell surface downregulation, and mechanisms of endocytosis. It is therefore used as a tool compound to study diverse biological processes. However, ikarugamycin commercial prices are very high, with up to 1300 € per 1 mg, thus limiting its application. We therefore set out to develop a high-yielding recombinant production platform of ikarugamycin by screening different expression vectors, recombinant host strains, and cultivation conditions. Overall, this has led to overproduction levels of more than 100 mg/L, which, together with a straightforward purification protocol, establishes biotechnological access to affordable ikarugamycin enabling its increased use in biomedical research in the future.

## 1. INTRODUCTION

Ikarugamycin (**1**) is a bacterial polycyclic tetramate macrolactams (PoTeMs). It was the first PoTeM to be discovered, isolated from *Streptomyces* sp. No. 8603 by Jomon *et al* in 1972.^[1]^ The chemical structures of all PoTeMs feature a tetramic acid moiety that is incorporated into a macrolactam ring system and further equipped with a variable set of fused carbocyclic rings. ikarugamycin (**1**)^[2]^ and clifednamides A/B (**2a/b**)^[3]^ have a 5/6/5 membered carbocyclic ring system, whereas other representatives contain a 5/5/6 ring system, such as HSAF (**3**)^[4, 5]^ and the frontalamides (**4**)^[6]^, or a 5/5 ring arrangement, as in cylindramide (**5**)^[7]^ and alteramide A (**6**)^[8]^. By further modifications at the core structure (e.g., by oxidative decoration), an impressive structural diversity is found in the PoTeM family.

In previous work, our group identified the biosynthetic gene cluster (BGC) encoding ikarugamycin (**1**) and showed that only three genes, *ikaABC*, are essential for its biosynthesis,^[9]^ a finding that was also validated by Zhang et al.^[10]^ Analogously to related PoTeM pathways,^[4, 6],[11]^ the iterative PKS/NRPS system IkaA catalyzes the formation of two polyene chains from acetyl-CoA and malonyl-CoA followed by their attachment to the nitrogen functions of L-ornithine with subsequent tetramic acid formation by a thioesterase.^[12, 13]^ Then, the oxidoreductase IkaB forms the two outer rings, putatively involving an NAD(P)H-dependent *C,C*-bond formation event combined with a [4+2] cycloaddition, although the exact order of events and mechanism remains unclear. The alcohol dehydrogenase IkaC catalyzes final formation of the inner ring, likely by an NAD(P)H-dependent reductive Michael addition-like reaction.^[10]^ All other characterized PoTeM BGCs harbor homologous genes, encoding enzymes catalyzing similar reductive reaction sequences for PoTeM core structure assembly.^[14]^ In addition, some BGCs carry genes coding for modifying tailoring enzymes, such as PoTeM hydroxylases^[15, 16]^ (often annotated as sterol desaturases), for example in HSAF (**3**) and frontalamide (**4**) biosynthesis.

Due to their high structural diversity, a broad variety of pharmacological activities are reported for the PoTeM natural product class that range from antifungal to antibacterial and antitumor properties.^[5, 6, 8]^ Ikarugamycin (**1**) exhibits antibacterial activity against gram-positive bacteria, as well as antiprotozoal^[1]^, antifungal^[17]^, anti-leukemic,^[18]^ and anti-inflammatory^[19]^ properties. Moreover, it shows inhibition of the uptake of oxidized low-density lipoprotein (LDL) in macrophages, the inhibition of Nef-induced CD4 cell surface downregulation for replication of HIV-1, and the suspension of clathrin-coated pits (CCP) dependent endocytosis.^[20, 21]^ Ikarugamycin (**1**) therefore is an interesting tool compound to study diverse biological processes and consequently is commercially available from multiple vendors with prices ranging from approx. 400 € to 1300 € per 1 mg. Despite significant research towards establishing alternative access to ikarugamycin (**1**) that has been conducted over the last decades, including several total synthetic^[22, 23, 24, 25, 26, 27]^ and one fully biocatalytic approach,^[28]^ production of **1** is still relying on fermentation. We therefore set out to establish a recombinant production platform utilizing our DiPaC approach^[29, 30, 31],[32]^ that allows large-scale biotechnological access combined with a streamlined isolation procedure.

## 2. RESULTS

### 2.1 Preparation of expression constructs

The *ika* BGC from *Streptomyces* sp. Tü623^[28]^ was selected as a template for DiPaC for downstream recombinant production of ikarugamycin (**1**) in diverse *Streptomyces* expression strains. Four different expression vectors, namely pSET152, pUWL201PW, pWHM4* and pWHM1120 were chosen as expression systems. The two vectors pUWL201PW and pWHM4* comprise the constitutive ermE or ermE-like promoter, whereas the pWHM1120 vector contains the inducible promoter tipA.

The pSET152 plasmid lacks a suitable promoter. Therefore the strong constitutive ermE promoter from the erythromycin BGC^[33]^ was introduced to generate the pSET152_ermE derivative. The *ika* BGC consisting of *ikaABC* was amplified in a single piece by PCR in four individual reactions, each with primers containing homology arms for the respective expression vector and the *ika* cluster to enable subsequent Gibson assembly to obtain the desired expression constructs.^[34]^ As a DNA template for the PCR amplification, a previously assembled fosmid containing the *ika* BGC was used.^[9]^ The entire *ika* cluster, covering *ikaABC* with 12.3 kbp in size, was amplified as a single product. The PCR product was purified using agarose gel electrophoresis and subsequent gel extraction. The purified DNA fragments were assembled by Gibson assembly and the final expression plasmids transformed into *E. coli* DH5*α*. This approach readily delivered all desired, correctly assembled expression vectors without the need for tedious construct screening (Figure 2, see ESI Chapter 7).

**Figure 1.**
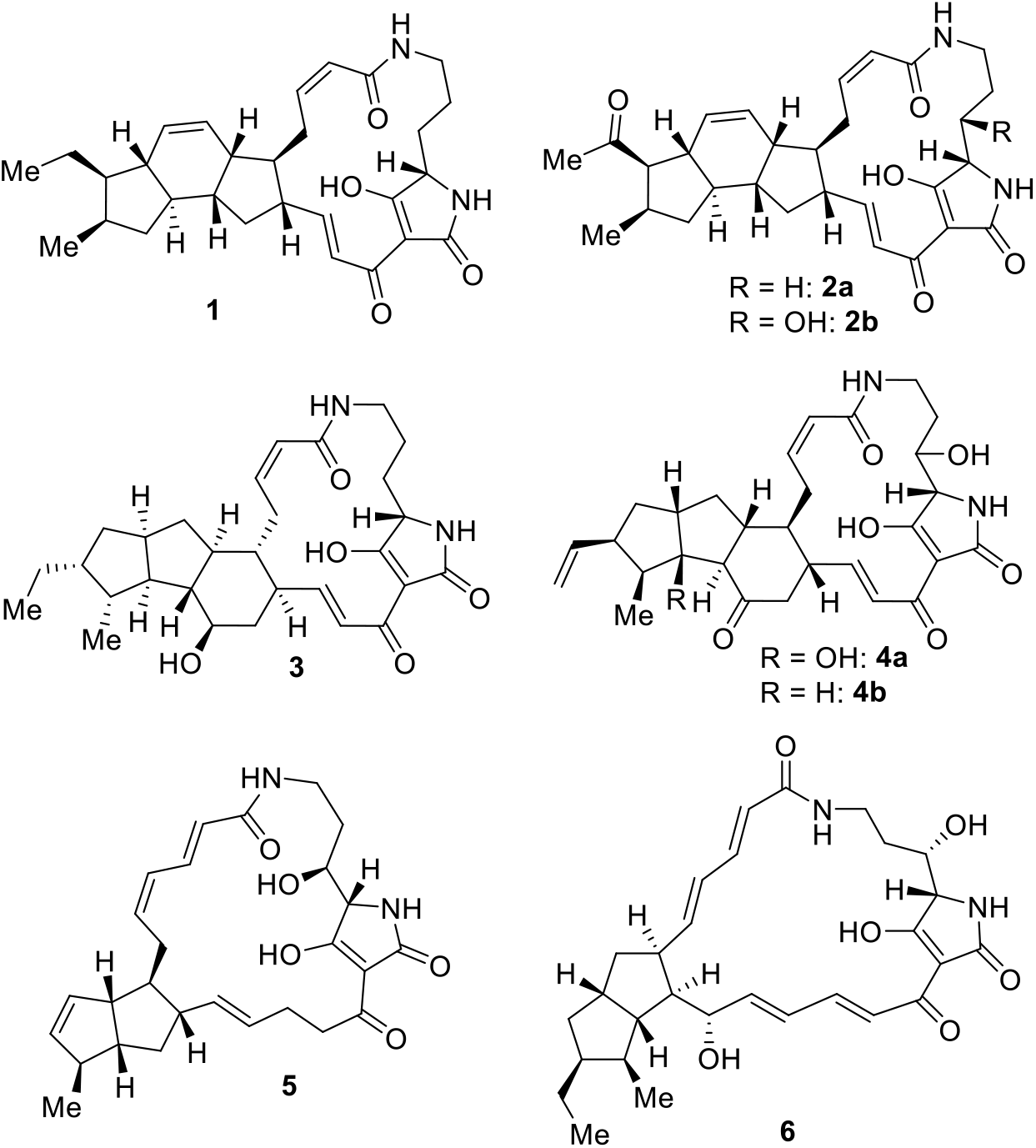
Chemical structures of selected representative showcase PoTeM structural diversity – ikarugamycin (**1**), clifednamides (**2**), HSAF (**3**), frontalamides (**4**), cylindramide (**5**) and alteramide A (**6**).

**Figure 2.**
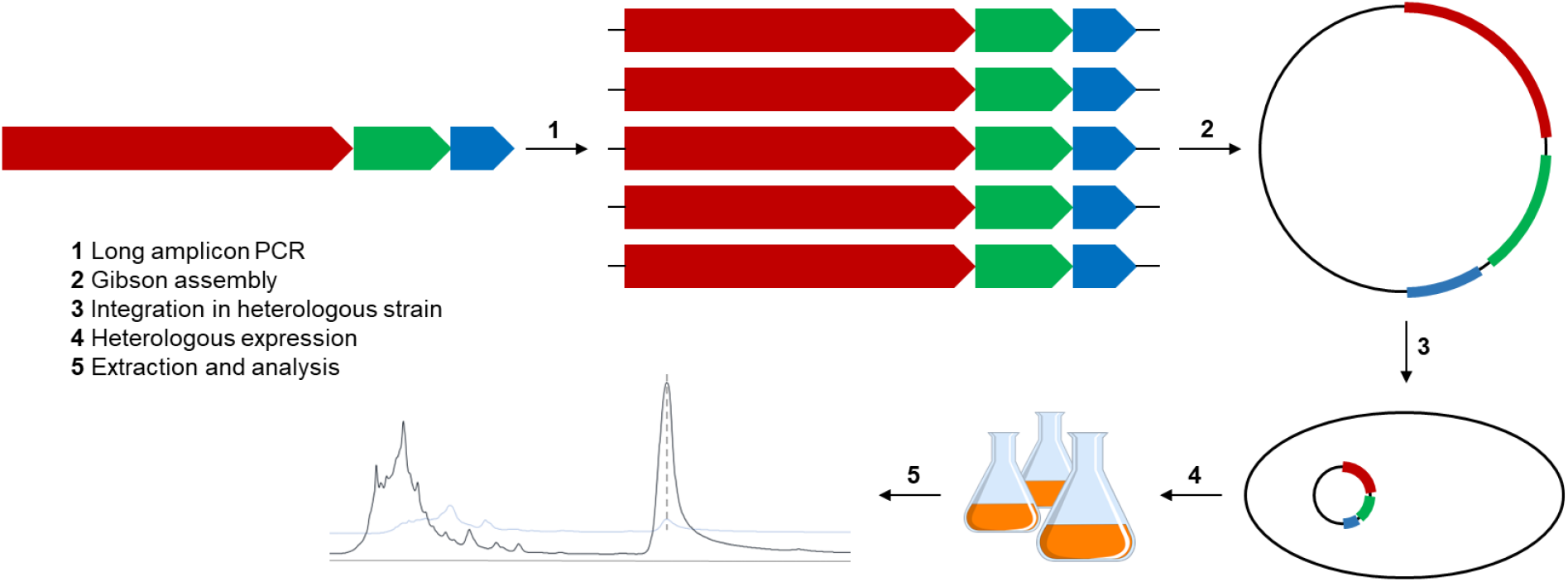
Schematic overview on the cloning of the *ika* BGC into the desired vectors, followed by the integration into recombinant host strains, heterologous expression, extraction, and chemical analysis.

### 2.2 Heterologous expression in *Streptomyces* species

After the successful transfer of the generated *ika* BGC expression vectors into *E. coli* DH5*α*, each construct was initially transferred into two different expression hosts, *Streptomyces albus* DSM40313 and *Streptomyces lividans* TK24 for a first assessment of production titers. Transfer to these *Streptomyces* sp. was either achieved by conjugation (for pSET152_ermE::*ika*) or by protoplast transformation (for pUWL201PW::*ika*, pWHM4*::*ika* and pWHM1120::*ika*). Single colonies of each resulting expression system were taken for the inoculation of pre-cultures in CASO or GYM medium at 28 °C for 48 to 72 hours. The main cultures of 50 mL [Zhang, ISP-4 and GYM medium (+Apra+NA, 50 mL medium in 250 mL Erlenmeyer flask)] were then inoculated with a ratio of 1:10 with the pre-culture and incubated at 28 °C for five or seven days. The cell pellets and supernatants were extracted with ethyl acetate and qualitatively analyzed by LC-MS. Unfortunately, the three constructs pUWL201PW::*ika*, pWHM4*::*ika* and pWHM1120::*ika* did not produce significant yields of the desired compound **1** and were thus omitted in the following optimization experiments.

To find optimal production conditions using expression vector pSET152_ermE::*ika*, six different *Streptomyces* sp. as heterologous hosts and eight different cultivation media were selected for screening. The plasmid pSET152_ermE::*ika* was transferred into the respective strains by conjugation, including *S. albus* DSM40313,^[35]^ *Streptomyces coelicolor* M1154,^[36]^ *S. lividans* TK24,^[37]^ *Streptomyces albus* B2P1, and *Streptomyces albus* Del14.^[38]^ In addition, we included a *Streptomyces* sp. that is natively harboring a PoTeM BGC. We therefore used *Streptomyces albus* DSM 40313, in which we deleted the cognate PoTeM BGC by homologous recombination, leading to expression strain *Streptomyces albus* KO5 (see chapter 4.1 and ESI, Figure S1). Strains were plated on GYM-agar and MS-agar plates. We found that there were no differences in colony growth on different plates, neither in terms of colony appearance nor in growth rate. Pre-cultures from single colonies were incubated in CASO medium (+Apra) at 28 °C for 72 hours and the main cultures were inoculated with 1:10 of pre-culture of each strain in each of the production media to be evaluated (Bennett’s, FMM, ISP4, ISP2, R5A, SGG, YEME, Zhang; +Apra; 50 mL medium in 250 mL Erlenmeyer flask; see Tables S2 for media compositions). For an initial fast pre-screening, all cultures were incubated for three, five and seven days (Figures S10-S15). Cell pellets and supernatants were extracted and analyzed by LC-MS for the evaluation of production titers (see Figure 3 and Figure S7 for representative examples). Pure ikarugamycin (**1**) was used to generate a calibration curve (see Figure S9) with a range of 3.12 ×10^−7^ mmol to 3.12 ×10^−4^ mmol to facilitate quick evaluation of recombinant production titers. Generally, these initial experiments suggested that the production of ikarugamycin (**1**) did likely not yet reach its maximum after seven days for several combinations of a particular recombinant host and medium, for example for *S. albus* KO5 pSEt152_ermE::*ika* in Zhang medium (see Figure 3). The maximum cultivation time for a more in-depth screening was thus extended to nine days (see below). Streptomyces strains *S. coelicolor* M1154 and *S. lividans* TK24 were excluded from further fermentation experiments due to generally very low production titers. The media ISP2, R5A and YEME had the lowest production of **1** in all *Streptomyces* strains and were therefore excluded in further optimization efforts.

**Figure 3.**
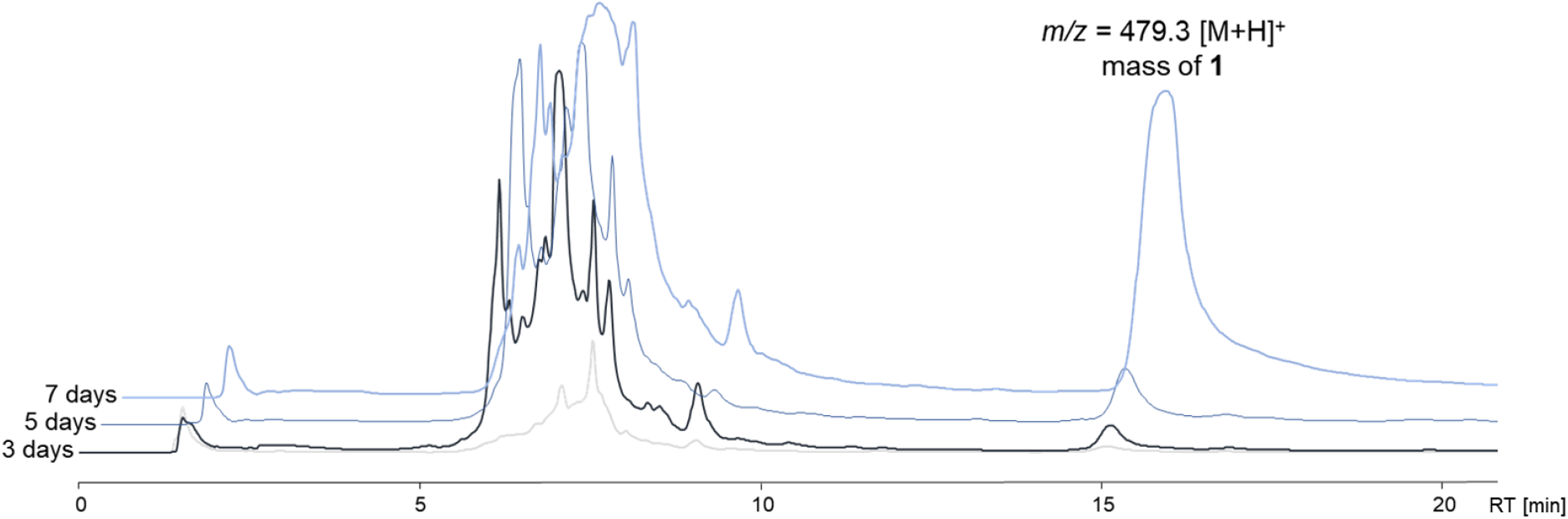
Example of HPLC chromatograms of extracts of the supernatant of the fermentation of *S. albus* KO5 pSET152_ermE::*ika* in Zhang medium after 3, 5 and 7 days.

### 2.3 Optimization of expression conditions

To determine the optimal time point for an efficient production of ikarugamycin (**1**), the remaining four recombinant strains were cultivated for five, seven and nine days in all selected production media with each experiment conducted in triplicate (Figures S16-S19). After extraction and analysis of the samples, it was evident that the three strains, *S. albus* DSM40313 pSET152_ermE::*ika, S. albus* KO5 pSET152_ermE::*ika* and *S. albus* Del14 pSET152_ermE::*ika* were the most promising production strains. However, each of these recombinant systems had maximum production of **1** in different media. The four best-performing media were Bennett’s, ISP4, SGG, and Zhang and production titers were indeed best for 5, 7 or 9 days of fermentation, depending on the production strain (Figure 4 and ESI, Chapter 5).

**Figure 4.**
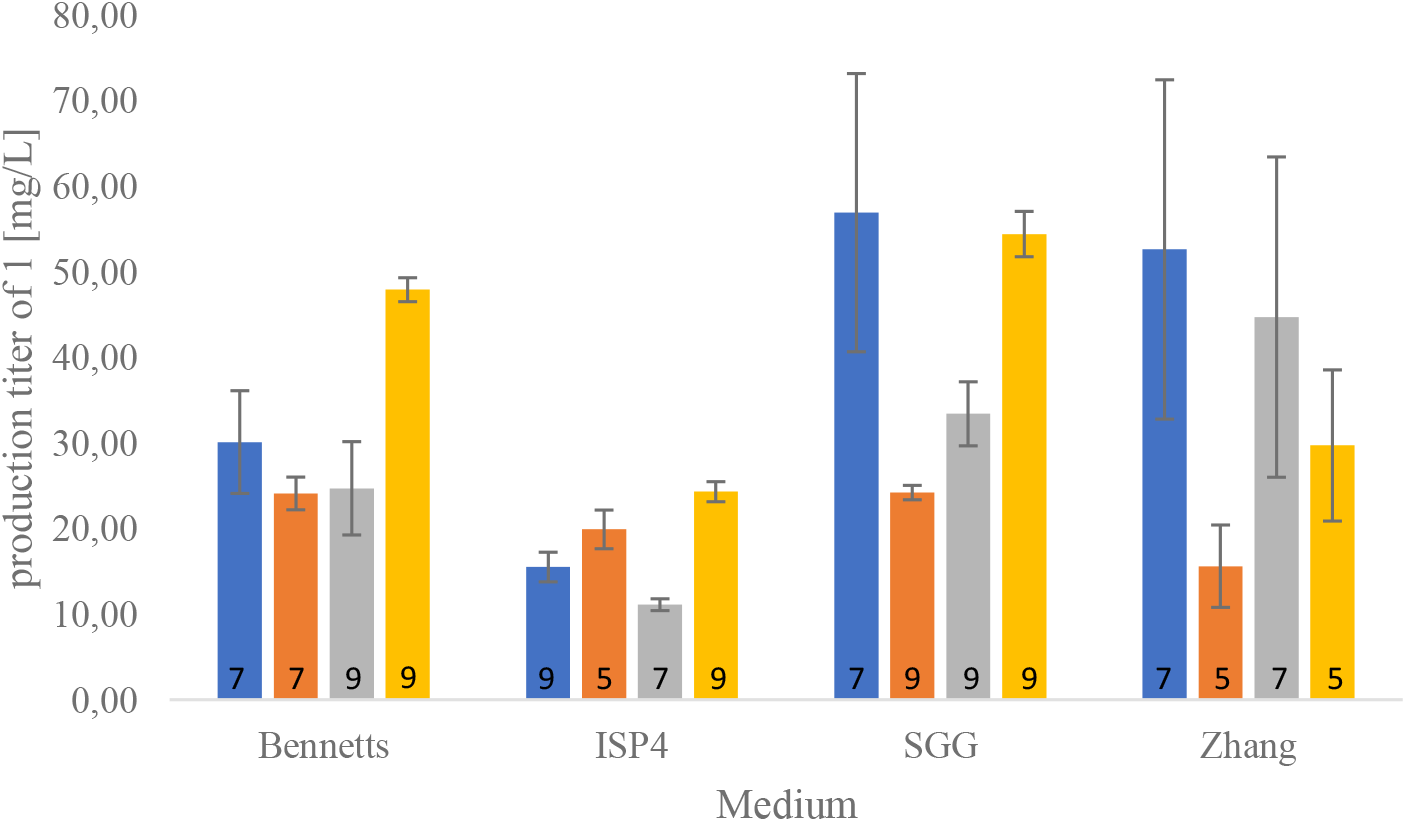
Overview on heterologous production titers of **1** in the four best-performing media (Bennetts, ISP4, SGG, Zhang) after 5, 7 or 9 days depending on host strain (indicated by bottom numbers [days]; see ESI, chapter 5). Strains: *S. albus* DSM40313 (blue), *S. albus* B2P1 (orange), *S. albus* KO5 (grey) and *S. albus* Del14 (yellow).

With these data in hands, we next turned to an upscaling experiment in which we increased the fermentation culture volume from 50 mL to 200 mL while keeping the ratio of fermentation broth volume *versus* cultivation flask size constant (cf. Figure S20). In addition, we tested alterations in the amount of pre-culture used for inoculation, ranging from 1:10 over 1:20 to 1:50 (cf. Figure S21). The amounts of ikarugamycin (**1**) were all calculated for cell lysate and pellet separately by using a calibration curve, but then combined to obtain one value for each cultivation. For these investigations, we used strains *S. albus* DSM40313 pSET152_ermE::*ika* and *S. albus* KO5 pSET152_ermE::*ika* both cultivated in Zhang medium, as this combination of host strain and medium delivered among the best production titers in above pre-screening experiments. After extraction of the cell pellet and the supernatant, chemical analysis revealed an average production titer of 78.59 mg/L ± 3.94 mg/L for *S. albus* DMS40313 pSET152_ermE::*ika* and of 109.61 mg/L ± 21.03 mg/L for *S. albus* KO5 pSET152_ermE::*ika*, both delivering best results with an inoculation ratio of 1:10.

Taken together, the best production conditions developed within this study involve strain *S. albus* KO5 pSET152_ermE::*ika*, its incubation in CASO for pre-culture production at 28 °C for 72 hours, followed by inoculation of the main culture with 10 vol% of the pre-culture into Zhang production medium (+Apra) with fermentation at 28 °C for seven days. After extraction of the cell pellet and the supernatant, this delivers an average overall yield of 109.6 mg/L (maximum value obtained was 135.0 mg/L).

Apart from optimized production titers, the isolation of a natural product in high purity can be very tedious, often requiring multiple chromatographic purification steps. In particular for PoTeMs such as **1**, generally low solubility in all common solvents makes isolation an especially challenging task. Initial attempts using flash chromatography on silica gave insufficient results in both, achieved purity and isolated yield. The method of choice was application of a fast preparative HPLC run for pre-purification, followed by an extraction/precipitation procedure to obtain high-purity **1**. Using this approach, **1** was obtained as a white powder in a very high purity (approx. 95% purity; based on HPLC and NMR analysis, see Figure S22-S28).

## 3. DISCUSSION

In conclusion, the efficient heterologous expression of ikarugamycin (**1**), shown in this work, enables the recombinant production of this important biomedical tool compound. Screening of four different expression vectors (pSET152_ermE, pUWL201PW, pWHM4* and pWHM1120) in a selection of up to six different *Streptomyces* sp. expression strains (*S. lividans* TK24, *S. coelicolor* M1154, *S. albus* DSM40313, *S. albus* KO5, *S. albus* B2P1 and *S. albus* Del14) in a diverse set of eight different production media (Bennett’s, FMM, ISP4, ISP2, R5A, SGG, YEME, Zhang) was performed. The three best producing strains, *S. albus* DSM40313, *S. albus* KO5, and *S. albus* Del14, were used for several titer optimization and scale-up experiments. The best production results were obtained with *S. albus* DMS40313 pSET152_ermE::*ika* and *S. albus* KO5 pSET152_ermE::*ika* with 78.59 mg/L ± 3.94 mg/L and of 109.61 mg/L ± 21.03 mg/L, respectively. Isolation of **1** was readily enabled in high purity, as shown by NMR and LC/MS analysis (cf. Fig. S20–S26). All other heterologous expression approaches reported so far were far less promising. Zhang *et al*. received 103 mg from a 20 L broth (5.15 mg/mL)^[10]^ and Lacret *et al*. obtained 8.9 mg from 1 L broth.^[17]^

With the methods shown in this work, an efficient and time-saving approach for biotechnological production of ikarugamycin (**1**) was established. The recombinant production platform can now be applied to reduce current ikarugamycin (**1**) commercial prices (in the range of 400 € to 1.300 € per 1 mg!) to pave the way for further in-depth testing of its pharmacological relevance and for its increased utilization as tool compound. Given production titers of more than 100 mg/L using our heterologous system and the straightforward isolation protocol enabling isolation of approx. 80% of produced **1** in pure form, a current market value in the range of 6.400–20.800 € can be generated from a single 200 mL expression culture. In addition, the high-level expression platform can also be utilized for expression of novel PoTeM BGCs, for engineering of PoTeM pathways, and for *in vivo* studies on PoTeM biosynthetic tailoring steps and will thus serve as valuable tool to the PoTeM research community.

## 4. EXPERIMENTAL PROCEDURES

### 4.1 Genetic deletion of the cognate PoTeM biosynthetic pathway in *S. albus* DSM 40313 to generate *S. albus* KO5

Deletion of the cognate PoTeM BGC of *S. albus* DSM 40313 was conducted by homologous recombination. For this, a pCC1FOS vector was used, which was linearized by PCR (pCC1FOS_bb_fwd/rev, Figure S1). Based on the genome data of *S. albus* J1074,^[39]^ which likewise produces PoTeM analogs, primers were designed to amplify the regions (3000 bp) upstream (HRup_fwd/rev) and downstream (HRdown_fwd/rev) of the PoTeM BGC. Additionally, a thiostreptone resistance cassette (*thioR*, including its promotor) was amplified from a pUWL201PW plasmid (ThioR_fwd/rev) and added between both homologous regions as selection marker. The four PCR products were purified by gel extraction and ligated by Gibson assembly resulting in pCC1FOS-HRup-*thioR*-HRdown. The reaction mix was used for transformation into *E. coli* DH5α.

The resulting plasmid was conjugated into *S. albus* DSM40313 using *E. coli* ET12567 pUZ8002 (see Chapter 4.3). Cells were incubated at 30 °C for 7 d to allow homologous recombination. To verify the knockout, several colonies were picked, grown in CASO medium, and genomic DNA (gDNA) was extracted. For the extraction of gDNA, cells were harvested (9.000 rpm, 10 min), washed with a NaCl-solution (0.9% (w/v)), and re-suspended in 5 mL lysis buffer (25 mM EDTA, 300 mM sucrose, 25 mM Tris-HCl, pH 7.5). The cells were lysed by three cycles of freezing on liquid nitrogen and thawing in a water bath (50 °C). Then, lysozyme (1 mg/mL) and RNase (10 μg/mL) were added, and the solution was incubated at 37 °C for 1 h. Subsequently, proteinase K (0.5 mg/mL) and SDS (final 1% (w/v)) was added, and the suspension was incubated again (37 °C, 30 min then 55 °C, 30 min). NaCl (final concentration 1 M) and acetyltrimethylammoniumbromide (final concentration 1% (w/v)) were added, and the sample was incubated at 65 °C for 10 min. An equal amount of phenol:chloroform:isoamylalcohol (25:24:1) solution was added to the sample and the mixture was inverted until it was homogenous, followed by incubation on ice (30 min) with repeated mixing. The phases were separated by centrifugation (9.000 rpm, 10 min), the aqueous phase was transferred into a fresh 15 mL centrifuge tube and washed two further times with the phenol:chloroform:isoamylalcohol solution. The aqueous phase was divided into 1.2 mL fractions and isopropanol (720 μL) was added. After gentle inverting (15–30 times), the DNA was pelleted (13.500 rpm, 30 min). The supernatant was removed, and the gDNA was washed with ethanol (1 mL, 70% (v/v)). After pelleting of the gDNA (13.500 rpm, 15 min), the supernatant was removed, and the gDNA was air dried. Extracted gDNA was dissolved in 50–200 μL TE-buffer (10 mM Tris-HCl, 1 mM EDTA) and used for validation of the genetic deletion. This screening PCR was performed to prove the presence of *thioR* (screening 1, see Figure S2). Additionally, a second PCR was conducted to verify the absence of the PoTeM BGC to avoid false positive results of the first screening, caused by impurities of pCC1FOS-HRup-*thioR*-HRdown (screening 2).

### 4.2 Amplification and cloning of the *ika* BGC

The entire *ika* BGC was amplified by long-amplicon PCR from a previously established fosmid^[9]^ using Q5 DNA polymerase and different primer pairs with homology overhangs to the respective expression plasmids (see Table S3). Reactions mixtures contained 1 × Q5 reaction buffer, 1 × high-GC enhancer, 200 µM dNTPs, 250 µM of each primer, 5–50 ng template DNA, and 0.125 µL Q5 polymerase (0.02 U/µL), and the required amount of ddH_2_O to reach 25 µL final volume. For PCR cycling conditions, see Table S9. Analysis of DNA was performed using agarose gel electrophoresis. 1% (w/v) agarose gels were prepared with 1x TAE buffer and supplemented with 1.75 µL SERVA DNA Strain Clear G per 40 mL. DNA samples were mixed with 5 × DNA loading dye (2.5 µL) and loaded to the gel that was run at 120 mV for 30 min. Gels were analyzed using a Bio-Imaging-System Gene Genius using GeneSnap software by Syngene.

PCR products were either purified using a PCR purification kit (Jena Bioscience) or after preparative agarose gel separation by using the Gel Extraction Kit (Peqlab) or the Monarch DNA Gel extraction kit (NEB). The vectors were linearized by restriction digest and dephosphorylated. For the latter, the heat-inactivated preparative restriction digest was mixed with 6 μL Antarctic phosphatase buffer, 2 μL Antarctic phosphatase in a total volume of 60 μL. The reaction was incubated at 37 °C overnight and the Antarctic phosphatase was heat-inactivated at 65 °C for 10 min. Further purification was conducted using the PCR purification kit. The vector inserts and expression vector backbones were fused using HiFi DNA Assembly Mix (NEB).^[34]^ For a single reaction batch 0.02 pmol vector and 0.1 pmol insert in a total volume of 10 µL were combined with the 2 × HiFi DNA Assembly Mix and incubated at 50 °C for 1 h. The Gibson assembly mix was used to transform chemically competent *E. coli* DH5α cells, which were prepared as follows: a LB culture (250 mL) was inoculated with a pre-culture (2.5 mL, freshly prepared) of *E. coli* DH5α and incubated at 37 °C until an OD_600_ of 0.5 to 0.6 was reached. The culture was centrifuged (5 min, 4.000 rpm, 4 °C) and the cell pellet was washed with cold glycerol solution (4 ×, 10%, v/v) on ice. In the last step, cells were re-suspended in 9.3 mL glycerol solution supplemented with 700 µL DMSO, and aliquoted into 100 µL batches. The aliquots were frozen in liquid nitrogen and stored at –80 °C. For transformation, 10 µL of the Gibson assembly mix were added to an aliquot of competent cells and incubated on ice for 30 min, followed by a heat shock at 42 °C for 1.5 min with subsequent cooling on ice for 2 min. SOC medium (900 µL) was added and the resulting culture was incubated for 1 h at 37 °C while shaking (200 rpm). The culture was centrifuged (5 min, 4.000 rpm), the concentrated cell pellet plated on LB agar with the required selection antibiotics and incubated at 37 °C overnight.

To verify the successful integration of a plasmid into *E. coli* (or *Streptomyces*, see below), a colony PCR was conducted using Taq polymerase. Single colonies were re-suspended in 50 µL ddH_2_O and 5 µL of this mixture were used as template for the PCR. To this cell suspension, 2.5 µL 10 × reaction buffer (200 mM Tris, 100 mM (NH_4_)_2_SO_4_, 100 mM KCl, 20 mM MgSO_4_, 1% Triton-X, pH 8.8), 1 µL DMSO, 100 µM dNTPs, 200 µM of each primer, and 0.125 µL of Taq polymerase were added and the resulting mixture was supplemented with ddH_2_O to reach a final volume of 25 µL. The PCR cycling was performed according to Table S4. PCR reactions were analyzed using agarose gel electrophoresis (see above). Positive colonies were used for plasmid isolation. For this, an LB overnight culture (5 mL) of the *E. coli* DH5α clone supplemented with the required selection antibiotic was prepared to use for purification of the plasmid DNA with the peqGOLD Plasmid Isolation Kit (Peqlab) according to the manufacturer’s instructions for ‘High copy number plasmids’, using ddH_2_O for plasmid elution from the spin column, instead of the elution buffer provided with the kit. The correct assembly of the expression vectors was validated by analytical restriction digest and sequencing (see Figures S3 to S6). For this analysis 240 ng of the DNA, 1 µL of the appropriate buffer (NEB) and 0.125 µL of each restriction enzyme were combined in 10 µL total reaction volume. The mixture was at least incubated for 1 h at 37 °C and analyzed using agarose gel electrophoresis (see above).

### 4.3 Conjugation into *Streptomyces* host strains

Horizontal conjugation of DNA from *E. coli* to *Streptomyces* was conducted by conjugation using *E. coli* ET12567 pUZ8002. The expression constructs were transformed into electrocompetent *E. coli* ET12567 pUZ8002 by electroporation. For production of electrocompetent cells, a 250 mL LB culture was inoculated with 2.5 mL of the preculture containing the *E. coli* strain. Cells were cultivated at 37 °C, 200 rpm until an OD_600_ between 0.4 and 0.5 was reached. Cells were washed with 10% (v/v) glycerol for three times with all washing steps being performed on ice and intermediate centrifugation at 4.000 rpm (5 min, 4 °C). Finally, cells were harvested by centrifugation, re-suspended in 500 µL 10% (v/v) glycerol, and aliquoted in 50 µL aliquots. Aliquots were frozen in liquid nitrogen and stored at –80 °C. For electroporation, between 5 to 100 ng DNA were added to electrocompetent cells and transferred to an electroporation cuvette and pulsed with 2.5 kV, approx. 5 ms. 900 µL SOC were added to the cells, which were transferred into a sterile 1.5 mL reaction tube and recovered at 37 °C, 200 rpm, for 1 h. The cells were selected on LB-agar containing Kan, Cam, and the specific selection antibiotic of the respective expression construct. Transfer was validated by colony PCR (see above). PCR-verified cells were grown in 15 mL LB (supplemented with Kan, Cam, specific selection antibiotic) at 37 °C, 200 rpm until an OD_600_ of 0.4 to 0.6 was reached. The cells were washed three times with 15 mL LB without any antibiotic with intermediate centrifugation (4.000 rpm, 5 min, 4 °C). In the last washing step, the cells were re-suspended in 500 µL LB. In parallel, *Streptomyces* spores (derived of the respective expression strain grown until sporylation on MS agar plates) needed to be heat-activated by re-suspending 10 µL spore samples in 500 mL 2xYT medium and treatment at 50 °C for 10 min. Spores were cooled on ice, added to the *E. coli* cells, and the mixture was harvested at 4.000 rpm, 2 min. The resulting pellet was plated on MS agar (supplemented with 10 mM MgCl_2_ and 60 mM CaCl_2_) and incubated for 16–20 h at 30 °C. After incubation, plates were overlaid with 1 mg NA and 1.25 mg Apra. The plates were sealed with parafilm and incubated for 4–7 d at 30 °C. To proceed with exconjugants, single clones were picked and re-suspended in 50 µL ddH_2_O. Of this cell suspension 25 µL were used to inoculate a 5 mL CASO preculture (supplemented with Apra and NA) for further expression experiments, and 25 µL were plated on MS agar (supplemented with Apra and NA) for medium-time storage. Successful integration of each plasmid into *Streptomyces* was tested by colony PCR (see above).

### 4.4 Cultivation of recombinant hosts, culture extraction, and chemical analysis

The above CASO precultures were incubated at 28 °C for three days while shaking (180 rpm). These precultures were employed to start new cultures in CASO bouillon (30–40 mL+Apra) using an inoculation ratio of 1:10. For production, the resulting second precultures were used to inoculate the production medium (50 mL to 1 L), using inoculation ratios of 1:10 to 1:50, and incubation was continued for the desired time (three to nine days) at 28 °C and 200 rpm. The ratio of culture volume to size of the applied Erlenmeyer flask was mostly 1:5, only 1:3 for 1 L cultures.

For further analysis, the cultures were centrifuged (4.500 rpm, 20 min) and supernatant and cell pellet extracted individually. For extraction of the supernatant, the pH was adjusted to 2– 3 using 1 M HCl and subsequently extracted with ethyl acetate (3 × volume equivalent to culture volume). The organic phases were combined, dried over MgSO_4_, filtered, and the solvent removed under reduced pressure. The cell pellets were re-suspended in acetone/methanol (1:1, 20 mL) and cells disrupted by using an ultrasonic bath for 30 minutes. Cell debris were removed by centrifugation (6.000 rpm, 10 minutes) and discarded. The organic solvent was dried over MgSO_4_ and removed *in vacuo*. All organic culture extracts were stored at –80 °C until further analysis.

Directly before analysis by HPLC-MS, the samples were dissolved in a defined volume of methanol (10 µL per 1 mL of cultivation medium) and the resulting solution filtered using a 0.22 µm syringe filter. Analysis was performed on an Azura® HPLC manufactured by Knauer, consisting of the following components: AS 6.1L sampler, P 6.1L pump, DAD 2.1L detector. This device was coupled to an ESI mass spectrometer manufactured by Advion with single-quadrupole mass analyzer. The system was controlled by ClarityChrom software in combination with Advion Mass Express software. Separation was achieved on a reversed-phase C8 column (Eurospher II, 100-5-C8A, 100 × 3 mm, 5 µm) using the following gradient with water (A) and acetonitrile (B) as mobile phases, both buffered with TFA (0.05 %): 0–2 min at 95% A; 2–10 min gradient to 45% A; 10–20 min at 45% A, 20–20.5 min change to 0% A; 20.5–24 min at 0% A; 24–24.5 min change to 95% A; 24.5–28 min at 95% A. Ikarugamycin (**1**, identified by retention time, UV spectrum, and MS data in each run) production titer was calculated based on integrated UV peak area at 280 nm using a calibration curve that was established using purified **1** (Figure S9).

### 4.5 Isolation of purified ikarugamycin (1)

Isolation of purified **1** was achieved by a combination of a fast pre-purification step using a short HPLC run followed by compound precipitation. Preparative HPLC was performed on a Jasco system consisting of an UV-1575 Intelligent UV/VIS-Detector, two PU-2086 Plus Intelligent Prep pumps, MIKA 1000 Dynamic Mixing Chamber, 1000 µL injection port and a LC-NetII/ADC. The system was controlled by the Galaxie software. Optimized preparative HPLC conditions for pre-purification were (water (A) and acetonitrile (B) as mobile phases, both buffered with 0.05 % TFA): 0–2 min at 95% A, 2–7 min gradient to 5% A, 8–13 min at 5% A, 13–13.5 min to 95% A, 13.5–15 min at 95% A. This delivered **1** as a light-brown amorphous solid, indicating the still insufficient purity (also evident from HPLC analysis; Figure S8). To further increase purity, an extraction/precipitation method was elaborated. Therefore, pre-purified **1** was dissolved in a mixture of acetone:methanol:ethyl acetate (1:1:1, 0.2 mL/mg), ddH_2_O (0.2 mL/mg) was added, and a small-scale extraction was executed. The aqueous phase was carefully discarded and the organic solvent was removed *in vacuo*. The procedure was repeated two to three times until **1** precipitated when starting to remove the solvent *in vacuo*, which allowed isolation of pure **1** by centrifugation or filtration.

## Supporting information

ESI File

## Data Availability Statement

All data that support the findings of this paper are reported in the experimental section or the ESI.

## Supporting Information

Additional supporting information can be found online in the Supporting Information section at the end of this article.

## Literature

[1] K. Jomon, Y. Kuroda, M. Ajisaka, H. Sakai, J. Antibiot. 1972, 25, 271–280.

[2] I. Shosuke, H. Yoshimasa, Bull. Chem. Soc. Jpn. 1977, 50, 1813–1820.

[3] S. Cao, J. A. Blodgett, J. Clardy, Org. Lett. 2010, 12, 4652–4654.

[4] F. Yu, K. Zaleta-Rivera, X. Zhu, J. Huffman, J. C. Millet, S. D. Harris, G. Yuen, X.-C. Li, L. Du, Antimicrob. Agents Chemother. 2007, 51, 64–72.

[5] P. Graupner, S. Thornburgh, J. Mathieson, E. Chapin, G. Kemmitt, J. Brown, C. Snipes, J. Antibiot. 1997, 50, 1014–1019.

[6] J. A. Blodgett, D.-C. Oh, S. Cao, C. R. Currie, R. Kolter, J. Clardy, Proc. Natl. Acad. Sci. U. S. A. 2010, 107, 11692–11697.

[7] S. Kanazawa, N. Fusetani, S. Matsunaga, Tetrahedron Lett. 1993, 34, 1065–1068.

[8] H. Shigemori, M. A. Bae, K. Yazawa, T. Sasaki, J. Kobayashi, J. Org. Chem. 1992, 57, 4317–4320.

[9] J. Antosch, F. Schaefers, T. A. Gulder, Angew. Chem. Int. Ed. 2014, 53, 3011–3014.

[10] G. Zhang, W. Zhang, Q. Zhang, T. Shi, L. Ma, Y. Zhu, S. Li, H. Zhang, Y. L. Zhao, R. Shi, Angew. Chem. Int. Ed. 2014, 53, 4840–4844.

[11] Y. Liu, H. Wang, R. Song, J. Chen, T. Li, Y. Li, L. Du, Y. Shen, Org. Lett. 2018, 20, 3504–3508.

[12] X. Mo, T. A. M. Gulder, Nat. Prod. Rep. 2021, 38, 1555–1566.

[13] L. Lou, G. Qian, Y. Xie, J. Hang, H. Chen, K. Zaleta-Rivera, Y. Li, Y. Shen, P. H. Dussault, F. Liu, J. Am. Chem. Soc. 2011, 133, 643–645.

[14] P. Harper Christopher, A. Day, M. Tsingos, E. Ding, E. Zeng, D. Stumpf Spencer, Y. Qi, A. Robinson, J. Greif, A. V. Blodgett Joshua, Appl. Environ. Microbiol. 2024, 90, e00600–00624.

[15] C. Greunke, J. Antosch, T. A. Gulder, Chem. Commun. 2015, 51, 5334–5336.

[16] Y. Li, J. Huffman, Y. Li, L. Du, Y. Shen, MedChemComm 2012, 3, 982–986.

[17] R. Lacret, D. Oves-Costales, C. Gomez, C. Diaz, M. de la Cruz, I. Perez-Victoria, F. Vicente, O. Genilloud, F. Reyes, Mar Drugs 2014, 13, 128–140.

[18] R. Popescu, E. H. Heiss, F. Ferk, A. Peschel, S. Knasmueller, V. M. Dirsch, G. Krupitza, B. Kopp, Mutat. Res. -Fundam. Mol. Mech. Mutagen. 2011, 709, 60–66.

[19] B. Malcomson, H. Wilson, E. Veglia, G. Thillaiyampalam, R. Barsden, S. Donegan, A. El Banna, J. S. Elborn, M. Ennis, C. Kelly, S. D. Zhang, B. C. Schock, Proc Natl Acad Sci U S A 2016, 113, E3725–3734.

[20] K. Hasumi, C. Shinohara, S. Naganuma, A. Endo, Eur. J. Biochem. 1992, 205, 841– 846.

[21] T. Luo, B. L. Fredericksen, K. Hasumi, A. Endo, J. V. Garcia, J Virol 2001, 75, 2488– 2492.

[22] R. K. Boeckman Jr, R. B. Perni, J. Org. Chem. 1986, 51, 5486–5489.

[23] R. K. Boeckman Jr, C. H. Weidner, R. B. Perni, J. J. Napier, J. Am. Chem. Soc. 1989, 111, 8036–8037.

[24] R. K. Boeckman, J. J. Napier, E. W. Thomas, R. I. Sato, J. Org. Chem. 1983, 48, 4152–4154.

[25] L. A. Paquette, D. Macdonald, L. G. Anderson, J. Am. Chem. Soc. 1990, 112, 9292– 9299.

[26] L. A. Paquette, D. Macdonald, L. G. Anderson, J. Wright, J. Am. Chem. Soc. 1989, 111, 8037–8039.

[27] W. R. Roush, C. K. Wada, J. Am. Chem. Soc. 1994, 116, 2151–2152.

[28] C. Greunke, A. Glöckle, J. Antosch, T. A. Gulder, Angew. Chem. Int. Ed. 2017, 56, 4351–4355.

[29] C. Greunke, E. R. Duell, P. M. D’Agostino, A. Glöckle, K. Lamm, T. A. M. Gulder, Metab. Eng. 2018, 47, 334–345.

[30] P. M. D’Agostino, T. A. M. Gulder, ACS Synth. Biol. 2018, 7, 1702–1708.

[31] E. R. Duell, P. M. D’Agostino, N. Shapiro, T. Woyke, T. M. Fuchs, T. A. M. Gulder, Microb. cell fact. 2019, 18, 32.

[32] P. M. D’Agostino, C. J. Seel, X. Ji, T. Gulder, T. A. M. Gulder, Nature Chemical Biology 2022, 18, 652–658.

[33] M. J. Bibb, G. R. Janssen, J. M. Ward, Gene 1985, 38, 215–226.

[34] D. G. Gibson, L. Young, R.-Y. Chuang, J. C. Venter, C. A. Hutchison, H. O. Smith, Nat. Methods 2009, 6, 343–345.

[35] D. P. Labeda, J. R. Doroghazi, K.-S. Ju, W. W. Metcalf, Int. J. Syst. Evol. Microbiol. 2014, 64, 894–900.

[36] G. Wang, T. Hosaka, K. Ochi, Appl. Environ. Microbiol. 2008, 74, 2834–2840.

[37] D. A. Hopwood, T. Kieser, H. M. Wright, M. J. Bibb, J. Gen. Microbiol. 1983, 129, 2257–2269.

[38] M. Myronovskyi, B. Rosenkränzer, S. Nadmid, P. Pujic, P. Normand, A. Luzhetskyy, Metabol. Engin. 2018, 49, 316–324.

[39] C. Olano, I. García, A. González, M. Rodriguez, D. Rozas, J. Rubio, M. Sánchez-Hidalgo, A. F. Braña, C. Méndez, J. A. Salas, Microb. Biotechnol. 2014, 7, 242–256.

