## Supplementary material for "Heterologous expression and optimization of fermentation conditions for recombinant ikarugamycin production": ESI File

#### Supporting Information

### 1 Material and Methods

#### 1.1 Commercial Materials

The used, commercially available materials were purchased from the following manufacturers: Carl Roth (Karlsruhe, Germany), BASF (Ludwigshafen, Germany), Grüssing (Filsum, Germany), Sigma-Aldrich (Taufkirchen, Germany), New England Biolabs (Frankfurt am Main, Germany), Jena Bioscience (Jena, Germany), Carbolution Chemicals (Saarbrücken, Germany), VWR (Darmstadt, Germany) and Thermo Fisher Scientific (Schwerte, Germany). The used solvents for extraction (EtOAc) and HPLC analysis (MeOH as sample solvent, MeCN as eluent) were obtained from Merck and Fisher Scientifics.

#### 1.2 NMR

Nuclear Magnetic Resonance (NMR) spectra were recorded on a Bruker AVANCE III 600 spectrometer at ambient temperature. The chemical shifts are given in δ-values (ppm) relative to TMS ($^{1}$H, $^{13}$C). $^{1}$H and $^{13}$C spectra were referenced internally using the residual solvent resonances (DMSO-$d_6$: $\delta_H$ = 2.50 ppm, $\delta_C$ = 39.52 ppm). The coupling constants J are given in Hertz [Hz] and determined assuming first-order spin-spin coupling. The following abbreviations were used for the allocation of signal multiplicities: s – singlet, bs – broad singlet, d – doublet, bd – broad doublet, t – triplet, bt – broad triplet, q – quartet, m – multiplet, or any combination thereof.

#### 1.3 Primers

**Table S1.** Primers used in this study.

| Name | Sequence (5' → 3') | Application |
|---|---|---|
| ermE*_RBS_OE_for | ATCTAGGAATTCGCGGTCGATCTTGACGGCTGGCGAGAGGTGCGGGGAGGATCTGACCGACGCGGTCCACACGTGGCACCGCGATGCTGT | Overlap extension of *ermE* promoter |
| ermE*_RBS_OE_rev | AGCTTAGAATTCGTCCGTACCTCCGTTGCTCCGCTGGATCCTACCAACCGGCACGATTGTGCCCACAACAGCATCGCGGTGCCACGTG | Overlap extension of *ermE* promoter |
| For_pSET152-ermE-rev | GCTGCAGGTCGACTCTAGAGAGGCCTTCCGTACCTCCGTTGCT | Amplification of *ermE* |
| Rev_pSET152-ermE-rev | ACAGCTATGACATGATTACGAATTCGCGGTCGATCTTGACGGC | Amplification of *ermE* |
| For_pSET152_ermE_rev-IKA | AGGTCGACTCTAGAGAGGCCTCTACAGGGCGACCAGGACCTTG | *ika* amplification pSET152_ermE |
| Rev_pSET152_ermE_rev-IKA | CAACGGAGGTACGGAAGGATGTATTCATGGATTCCATGCACCACCCTGC | *ika* amplification pSET152_ermE |
| For_pULW201PW-Ika | GACGGTATCGATAAGCTTGATATCGAATTCATGGATTCCATGCACCACCCTGC | *ika* amplification pUWL201PW |
| Rev_pUWL201PW-Ika | TGCAGAGCTTCTAGAACTAGTGGATCCTACAGGGCGACCAGGACCTTG | *ika* amplification pUWL201PW |
| For_pWHM4*_IKA | GATCCCCGGGTACCGAGCTCGAATTCATGGATTCCATGCACCACC | *ika* amplification pWHM4* |
| Rev_pWHM4*_IKA_neu | GTAAAACGACGGCCAGTGAATTCTACAGGGCGACCAGGACCTTG | *ika* amplification pWHM4* |
| For_pWHM1120-IKA | TTGCATGCCTGCAGGTCGACTCTAGAATGGATTCCATGCACCACCCTGC | *ika* amplification pWHM41120 |
| Rev_pWHM1120-IKA | AGCTCGGTACCCGGGGATCCTCTACAGGGCGACCAGGACCTTG | *ika* amplification pWHM41120 |
| SEQ-Primer TüAlcD ForII | GGTGTTCACGATGCTCA | Sequencing Primer |
| Screening_ikaA_rev | GCAGGATCTTGTAGTCGA | Sequencing Primer |
| pUWL201PWseq_rev_AL4 | GATGTCGGACCGGAGTT | Sequencing Primer |
| pUWL201PWseq_AL6 | CAATACGCAAACCGCCTCT | Sequencing Primer |
| M13_for | GTAAAACGACGGCCAGT | Sequencing Primer |
| M13_rev | CAGGAAACAGCTATGAC | Sequencing Primer |
| pCC1FOS_bb_fwd | GCGACACACTTGCATCGG | Primer for KO Exp. |
| pCC1FOS_bb_rev | CAGGCGTAGCAACCAGGC | Primer for KO Exp. |
| HRup_fwd | ACCGCGCCATAGGAATCCGTCCGGGACAATACC | Primer for KO Exp. |
| HRup_rev | ACGCCTGGTTGCTACGCCTGGGCAGCCTGATCCCGCCG | Primer for KO Exp. |
| HRdown_fwd | ATCCGATGCAAGTGTGTCGCGCCAGGGCGAGCTGGTTGG | Primer for KO Exp. |
| HRdown_rev | CAACCGATAAGCCCGGCCGGCACCTCAC | Primer for KO Exp. |
| ThioR_fwd | CCGGCCGGGCTTATCGGTTGGCCGCGAGATTCCTG | Primer for KO Exp. |
| ThioR_rev | ACGGATTCCTATGGCGCGGTCGCGGTCG | Primer for KO Exp. |

#### 1.4 Media and buffers

**Table S2.** Composition of cultivation media used in this study.

| Medium | Components |
|---|---|
| GYM agar | 4.00 g D-glucose<br>4.00 g yeast extract<br>10.0 g malt extract<br>2.00 g $CaCO_3$<br>12.0 g agar<br>Add 1.00 L $ddH_2O$ |
| MS agar | 10.0 g agar<br>10.0 g mannitol<br>10.0 g soya flour<br>Add 475 mL $ddH_2O$<br>After sterilization add $MgCl_2$ (10 mM) and $CaCl_2$ (60 mM) |
| SOB/SOC | 20.0 g tryptone<br>5.00 g yeast extract<br>0.58 g NaCl<br>0.19 g KCl<br>Add 980 mL $ddH_2O$<br>After sterilization add $MgCl_2$ (10 mM) and $MgSO_4$ (10 mM)<br>For transforming SOB into SOC 9 mL of D-glucose solution (40%) was added |
| 2 x YT | 16.0 g tryptone<br>10.0 g yeast extract<br>5.00 g NaCl<br>Add 1 L $ddH_2O$.<br>Adjust the pH to 7.0 with NaOH. |
| Bennett's | 20.0 g starch<br>20.0 g Pharmamedia<br>10.0 g corn steep liquor<br>3.00 g $CaCO_3$<br>Add 1.00 L $ddH_2O$ |
| FMM (Fischmehl-Medium) | 20.0 g D-glucose<br>10.0 g fish flour<br>1.00 g $CaCO_3$<br>Add 1.00 L $ddH_2O$<br>Adjust pH to 7.0 |
| ISP-4 | 10.0 g soluble starch<br>1.00 g $MgSO_4$<br>1.00 g NaCl<br>2.00 g $(NH_4)_2SO_4$<br>2.00 g $CaCO_3$<br>Add 1.00 L $ddH_2O$<br>Add trace salts (1.00 mg $FeSO_4$, 1.00 mg $MnCl_2$, 1.00 mg $ZnSO_4$) |

| | |
|---|---|
| ISP-2 | 4.00 g yeast extract powder<br>10.0 g malt extract powder<br>4.00 g D-glucose<br>Add 1.00 L dd$H_2O$<br>Adjust pH to 7.2 with KOH |
| R5A | 103 g sucrose<br>0.25 g $K_2SO_4$<br>10.1 g $MgCl_2 \cdot 6\ H_2O$<br>10.0 g D-glucose<br>0.10 g casamino acids<br>5.00 g yeast extract<br>21.0 g MOPS<br>Add 1.00 L dd$H_2O$<br>Adjust pH to 6.8<br>Add 2.00 mL trace element solution (200 mg/L $FeCl_3 \cdot 6\ H_2O$, 10 mg/L $CuCl_2 \cdot 2\ H_2O$, 10 mg/L $MnCl_2 \cdot 6\ H_2O$, 10 mg/L $Na_2B_4O_7 \cdot 10\ H_2O$, 10 mg/L $(NH_4)_6Mo_7O_{24} \cdot 4\ H_2O$, 40 mg/L $ZnCl_2$) |
| SGG | 10.0 g D-glucose<br>10.0 g glycerol<br>2.50 g cornsteep powder<br>5.00 g peptone<br>10.0 g soluble starch<br>2.00 g yeast extract<br>3.00 g $CaCO_3$<br>1.00 g NaCl<br>Add 1.00 L dd$H_2O$<br>Adjust pH to 7.3 |
| YEME | 103 g sucrose<br>3.00 g yeast extract<br>5.00 g peptone<br>3.00 g malt extract<br>10.0 g D-glucose<br>Add 1.00 L dd$H_2O$<br>Adjust pH to 7.0 with NaOH |
| Zhang medium | 25.0 g D-glucose<br>7.00 g soybean powder<br>2.50 g yeast extract<br>5.00 g $(NH_4)_2SO_4$<br>4.00 g NaCl<br>8.00 g $CaCO_3$<br>0.40 g $KH_2PO_4$<br>Add 1.00 L dd$H_2O$ |

#### 1.5 Antibiotics

**Table S3.** Antibiotics used in this study.

| Name | Concentration |
|---|---|
| Ampicillin (Amp) | 100 µg/mL |
| Apramycin (Apra) | 30.0 µg/mL |
| Chloramphenicol (Cam) | 25.0 µg/mL |
| Kanamycin (Kan) | 50.0 µg/mL |
| Nalidixic Acid (NA) | 25.0 µg/mL |
| Thiostreptone (Thio) | 25.0 µg/mL |

#### 1.6 Polymerase chain reaction (PCR) cycling conditions

**Table S4.** Program for colony PCR using Taq polymerase.

| Step | Time | Temperature |
|---|---|---|
| 1 | 5 min | 94 °C |
| 2 | 45 s | 94 °C |
| 3 | 30 s | 52-72 °C (according to melting point of primer) |
| 4 | 30 s | 68 °C |
| 5 | 34 cycles (step 2-4) | |
| 6 | 5 min | 68 °C |
| 7 | ∞ | 16 °C |

**Table S5.** Program for long amplicon PCR using Q5 polymerase.

| Step | Time | Temperature |
|---|---|---|
| 1 | 45 s | 98 °C |
| 2 | 10 s | 98 °C |
| 3 | 30 s | 52-72 °C (according to melting point of primer) |
| 4 | 30 s | 72 °C |
| 5 | 30 cycles (step 2-4) | |
| 6 | 5 min | 72 °C |
| 7 | ∞ | 16 °C |

#### 2 Genetic Deletion, Plasmid Cloning, and Construct Verification

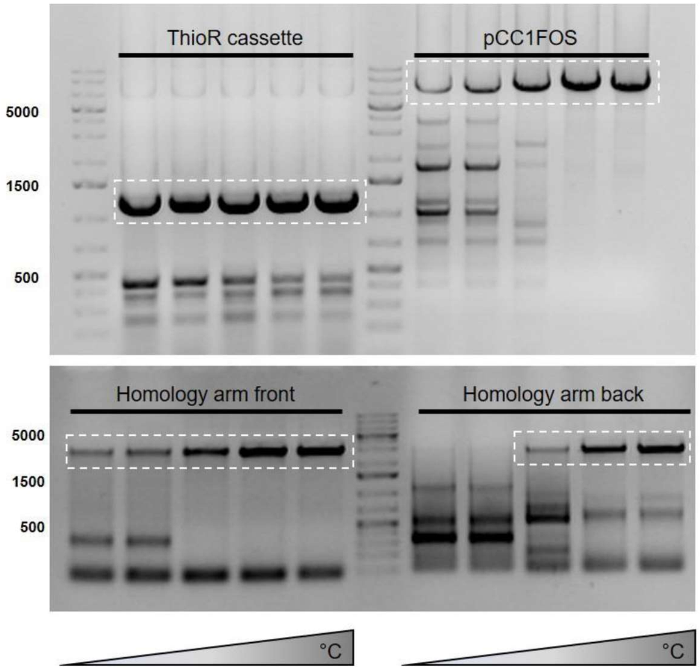

**Figure S1.** Construction of pCC1FOS-HRup-*thioR*-HRdown. Gradient PCR of the ThioR cassette (1176 bp), the pCC1FOS backbone (7205 bp), and the two homology arms (3030 bp).

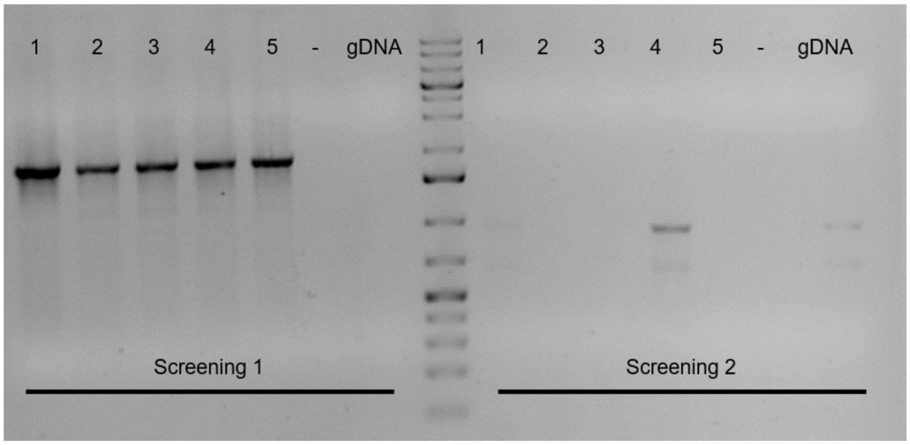

**Figure S2.** Positive and negative screening of the gDNA of the ΔPoTeM strain. Five exconjugants were examined to validate the accomplished homologous recombination to delete the native *S. albus* PoTeM cluster. Screening 1 was performed using a forward primer binding at the end of the front homology arm and a reverse primer binding at the beginning of the back homology arm (1876 bp). Screening 2 was performed using primers binding inside the knocked-out PoTeM BGC, detecting false positive results of screening 1 due to the presents of the non-integrated K.O.-plasmid (exconjugant 1 and 4).

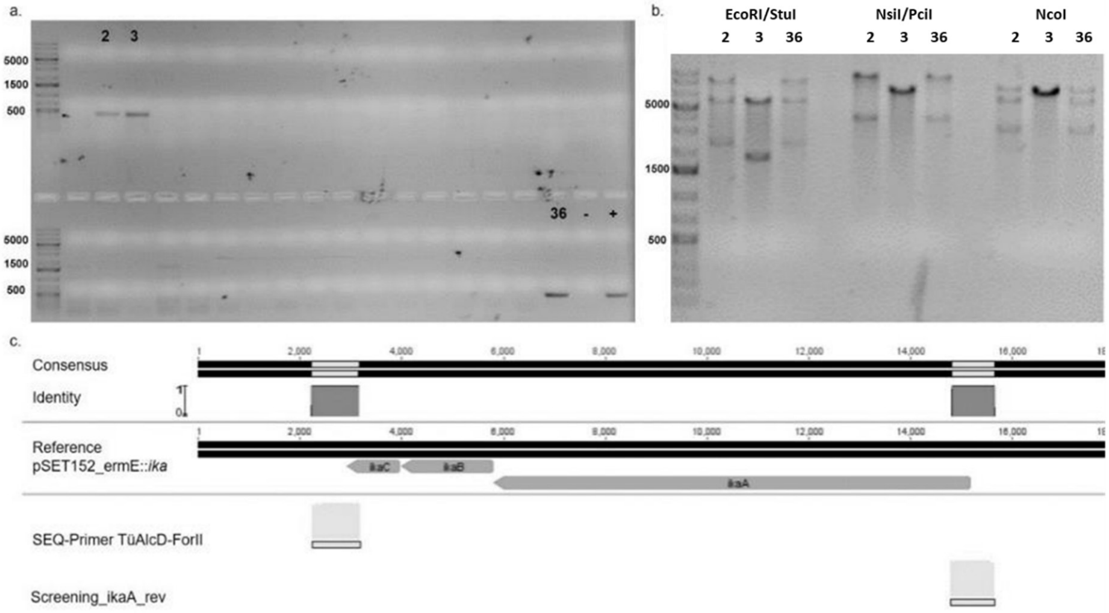

**Figure S3.** Cloning verification for pSET152_ermE::*ika*. **a**. Results of the colony PCR with possibly positive clones 2, 3, and 36; negative control conducted with water; positive control 1 µL of Gibson assembly reaction. **b**. Results of analytical restriction digest with all clones; clone 2 showed the expected restriction pattern and was submitted for sequencing. **c**. Sequencing results of clone 2 with primers: SEQ-Primer TüAlcD-ForII and Screening _IKA_rev, verifying a successful cloning of the *ika* BGC into pSET152_ermE.

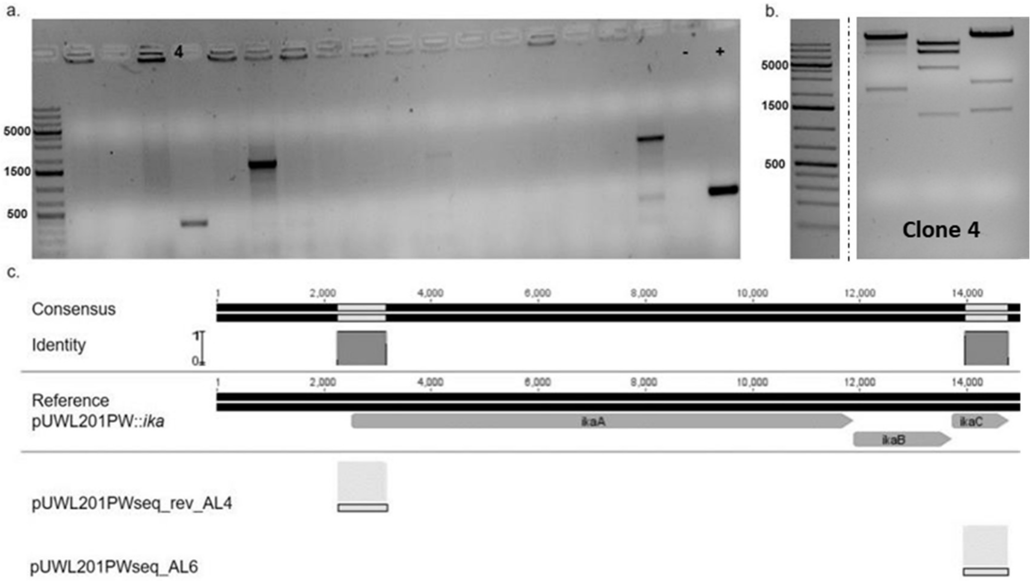

**Figure S4.** Cloning verification for pUWL201PW::*ika*. **a**. Results of colony PCR with possibly positive clone 4, negative control conducted with water; positive control 1 µL Gibson assembly reaction mixture. **b**. Results of analytical restriction digestion (dotted line indicates excision of unrelated restriction digest) with ScaI (left lane), NcoI (central lane), and StoI/AseI (right lane) for clone 4; clone 4 showed the expected restriction pattern and was submitted for sequencing. **c**. Sequencing results of clone 4 with primers: pUWL201Pwseq_rev_AL4 and pUWL2019Pwseq_AL6, verifying a successful cloning of the *ika* BGC into pUWL201PW.

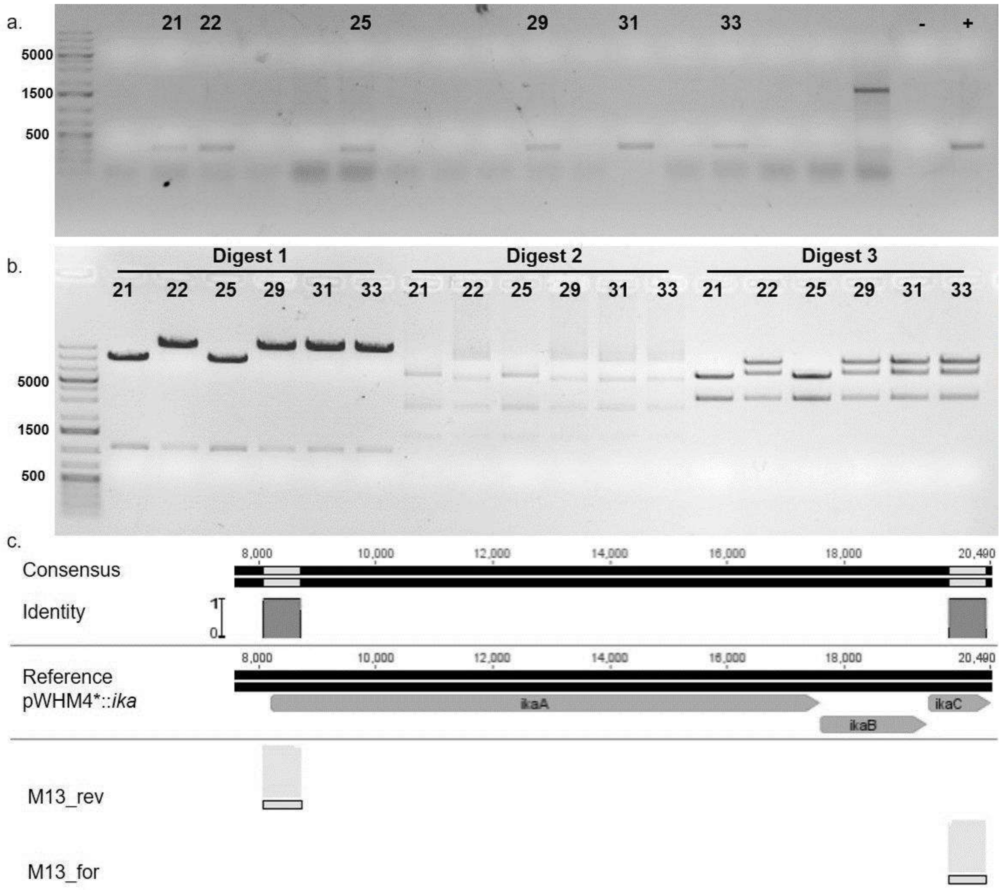

**Figure S5.** Cloning verification for pWHM4*::*ika*. a. Results of colony PCR with possibly positive clones 21, 22, 25, 29, 31, and 33; negative control was conducted with water; positive control 1 µL of Gibson assembly reaction mixture. b. Results of analytical restriction digest of all clones with EcoRV/HindIII (digest 1), BsmI (digest 2), and StuI/XbaI (digest 3); clone 22 showed the expected restriction pattern and was submitted for sequencing. c. Sequencing result of clone 22 with primers: M13_rev and M13_for, verifying a successful cloning of the ika BGC into pWHM4*.

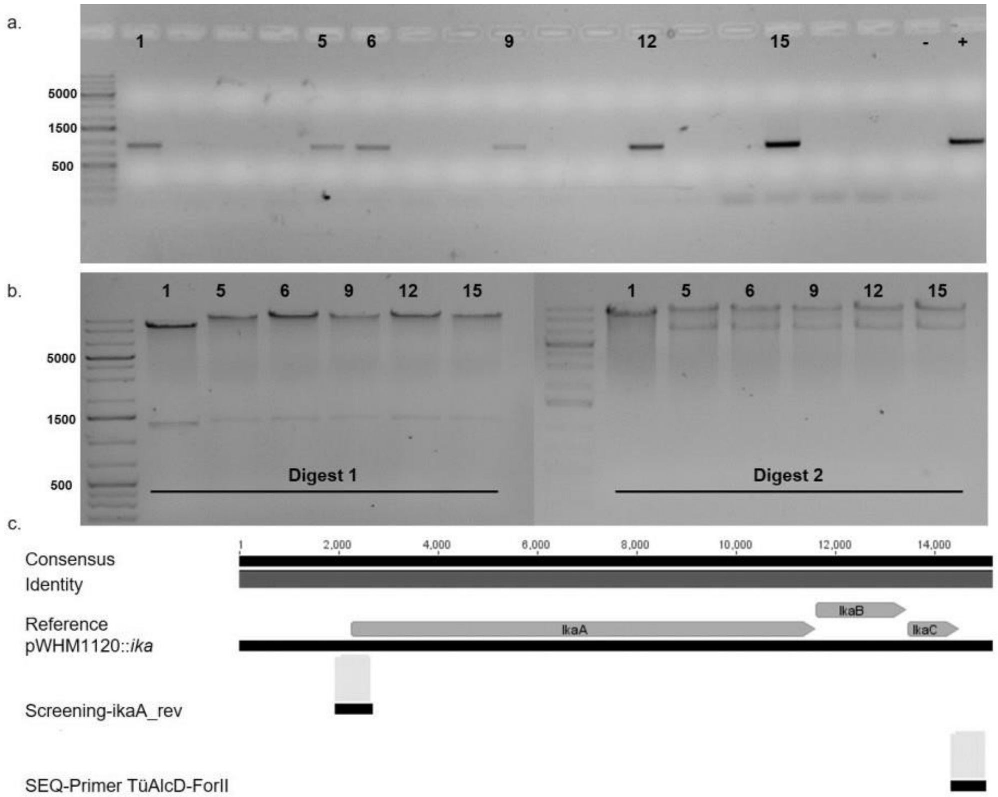

**Figure S6.** Cloning verification for pWHM1120::*ika*. **a**. Results of colony PCR with possibly positive clones 1, 5, 6, 9, 12, and 15; negative control conducted with water; positive control 1 µl Gibson assembly reaction mixture. **b**. Results of analytical restriction digest of all clones using ScaI (digest 1) and EcoRV/NsiI (digest 2); clone 15 showed the expected restriction pattern and was submitted for sequencing. **c**. Sequencing results for clone 15 with primers: Screening-ikaA_rev and SEQPrimer TüAlcD-ForII, verifying a successful cloning of the *ika* BGC into pWHM1120.

### 3 HPLC and HPLC-MS Data

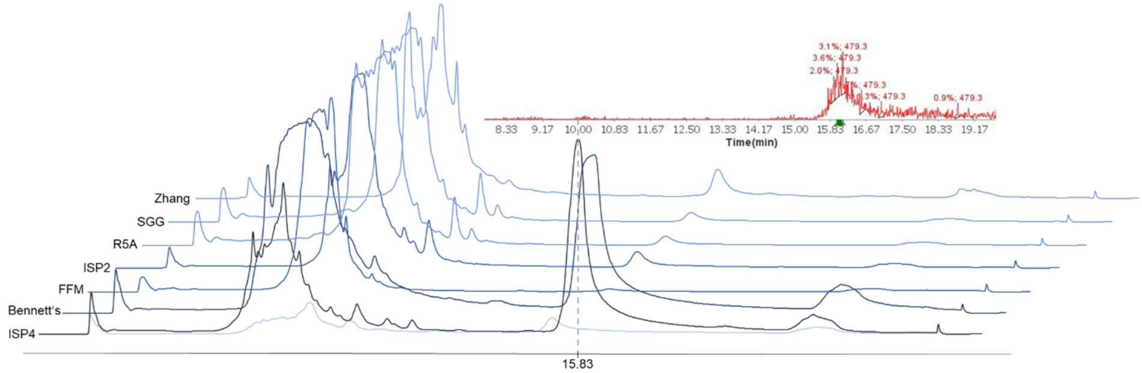

**Figure S7.** Representative HPLC-UV chromatograms at 280 nm (depicted from 8–19 min) for the analysis of supernatant of expression cultures in all tested media, exemplarily shown for *S. albus* KO5 pSEt152_ermE::*ika*. MS trace at *m/z* = 479.4 corresponding to ikarugamycin (**1**) at 15.83 min shown in red.

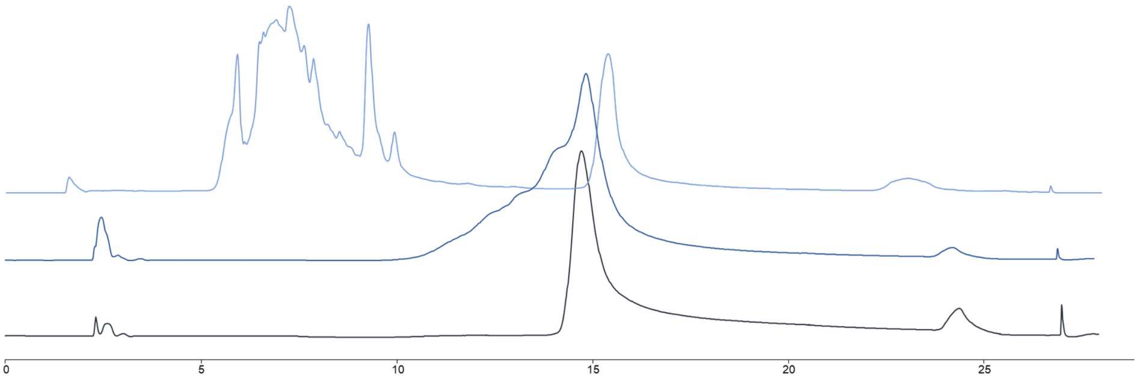

**Figure S8.** HPLC-UV chromatograms at 280 nm monitoring purification progress of **1** (exemplarily shown for extraction from *S. albus* DSM40313 pSET152_ermE::*ika*). Top: raw extract of the supernatant using Bennett's medium (3 d); middle: after HPLC pre-purification; bottom: after final precipitation step.

#### 4 Calibration Curve of Ikarugamycin (1)

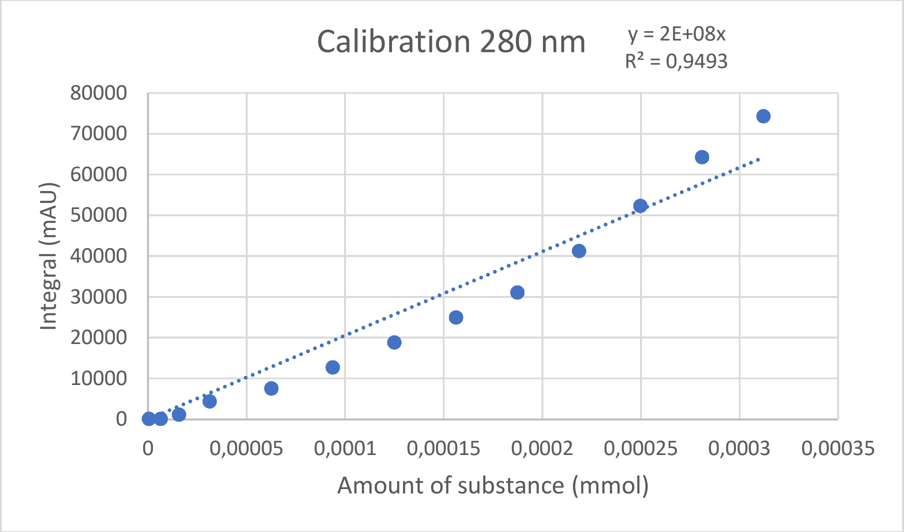

**Figure S9.** Calibration curve for quantification of **1** recorded by HPLC (280 nm) in the range of $3.12 \times 10^{-7}$ mmol to $3.12 \times 10^{-4}$ mmol.

#### 5 Evaluation of Expression conditions

Pre-screening of six different *Streptomyces* host strains in eight different media each. Extraction of 50 mL main culture after 3 (blue), 5 (orange), and 7 (grey) days of cultivation. Combined yields of culture media and cell extracts are shown.

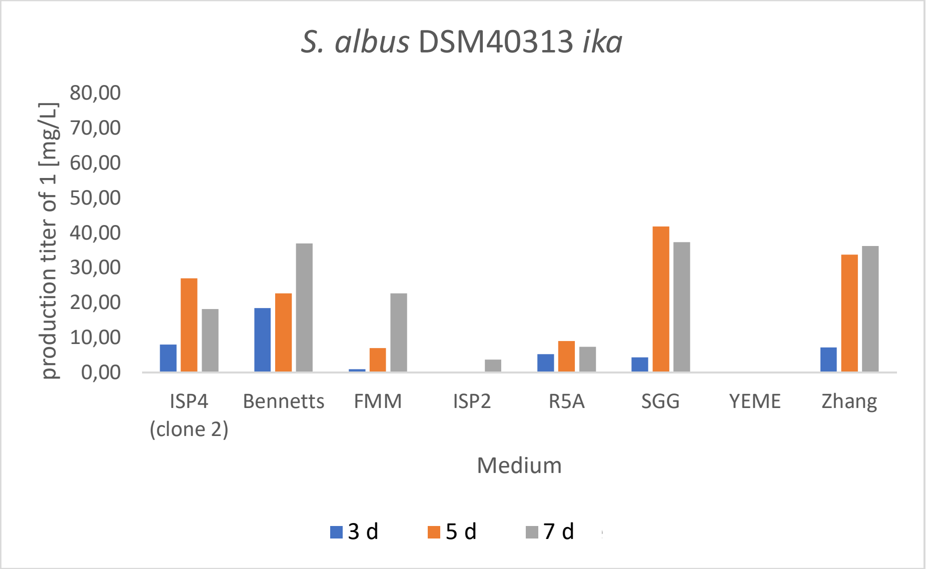

**Figure S10.** Production titers of **1** in *S. albus* DSM40313.

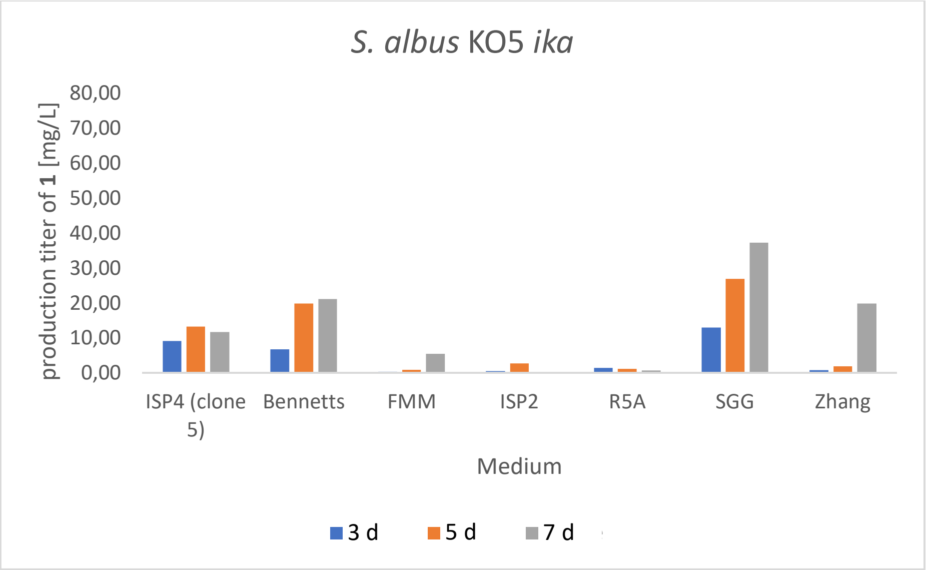

**Figure S11.** Production titers of **1** in *S. albus* KO5.

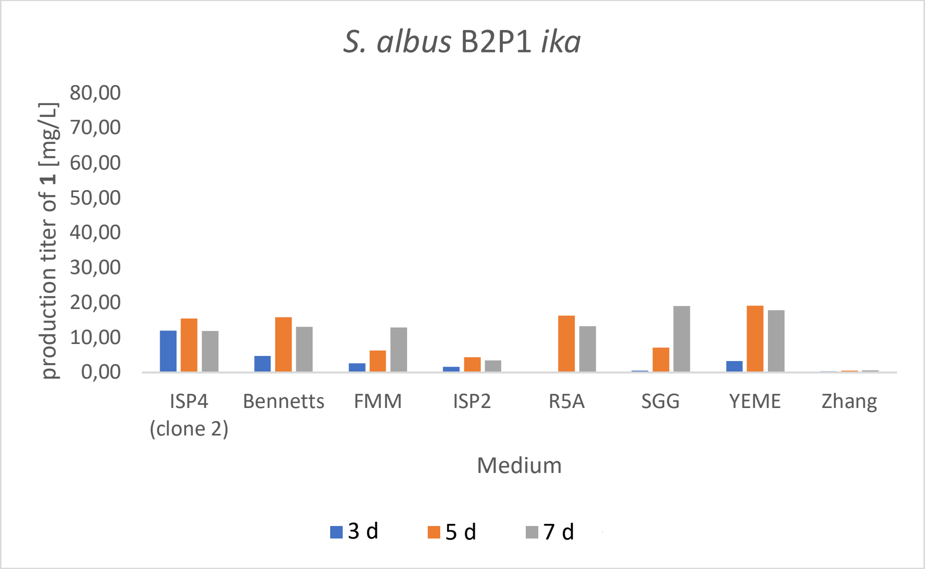

**Figure S12.** Production titers of **1** in *S. albus* B2P1.

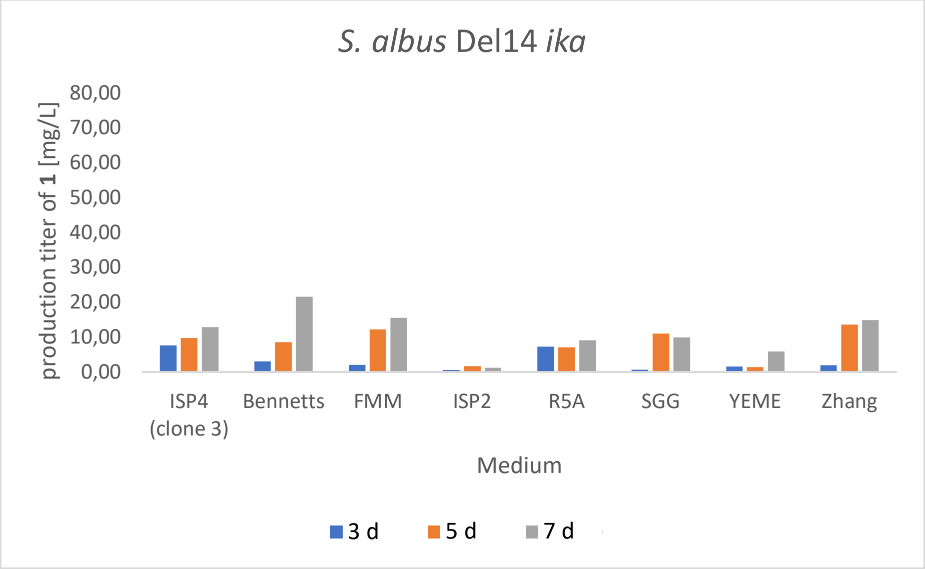

**Figure S13.** Production titers of **1** in *S. albus* Del14.

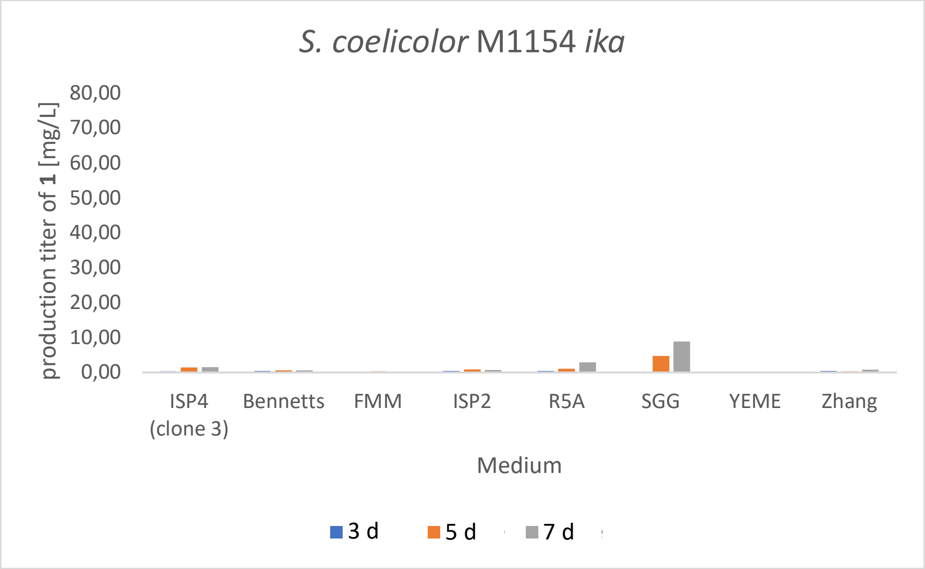

**Figure S14.** Production titers of **1** in *S. coelicolor* M1154.

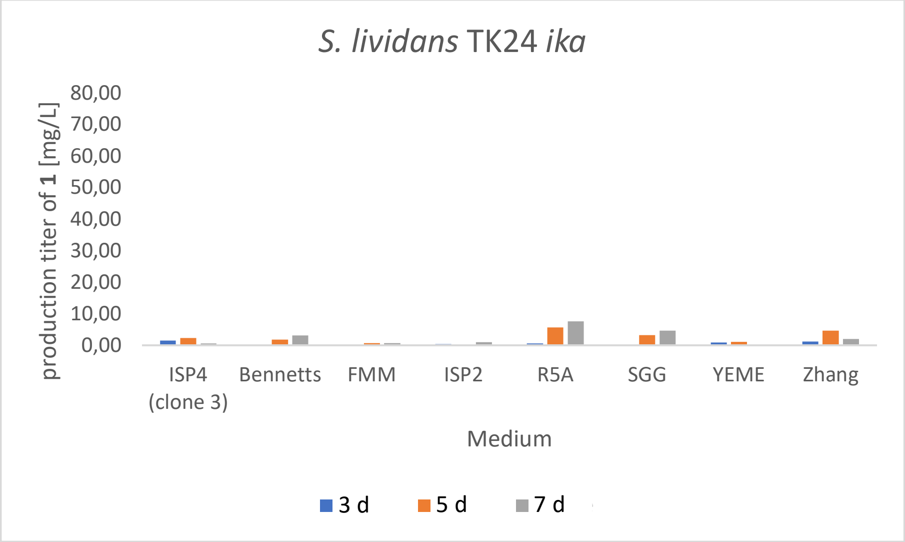

**Figure S15.** Production titers of **1** in *S. lividans* TK24.

In-depth screening of the four best-performing *Streptomyces* strains from pre-screening experiments in four/five different media. Extraction of 50 mL main cultures on days 5, 7 and 9. Titers were determined using biological triplicates.

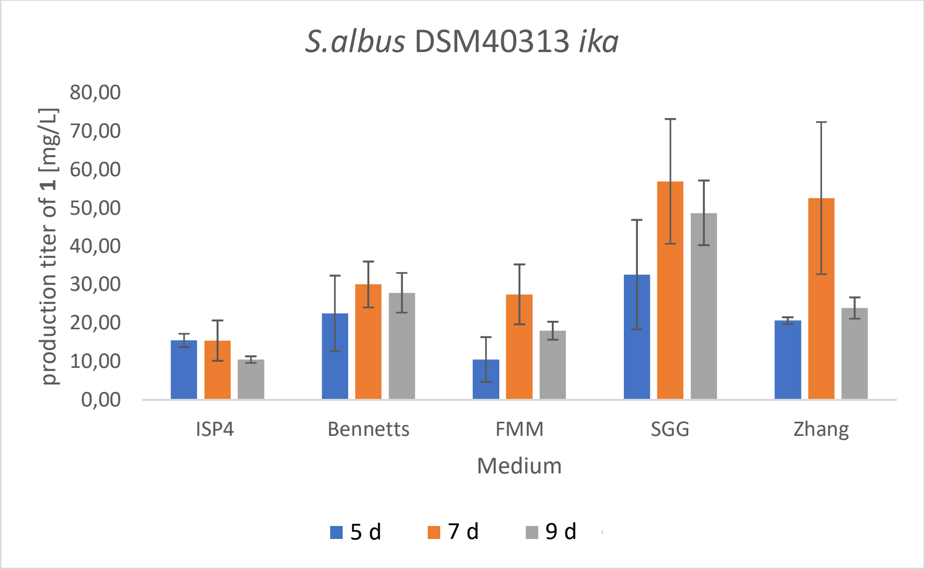

**Figure S16.** Production titer of **1** in *S. albus* DSM40313.

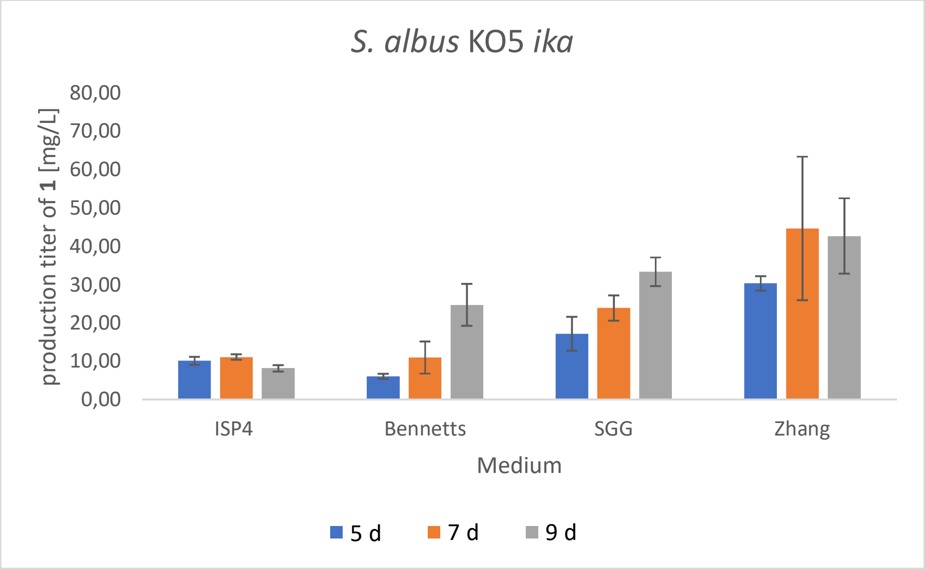

**Figure S17.** Production titer of **1** in *S. albus* KO5.

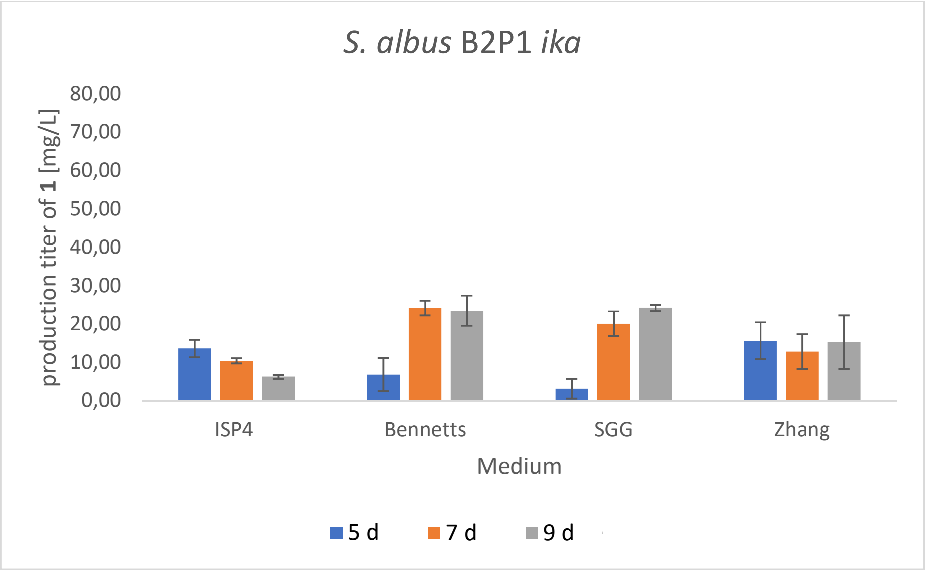

**Figure S18.** Production titer of **1** in *S. albus* B2P1.

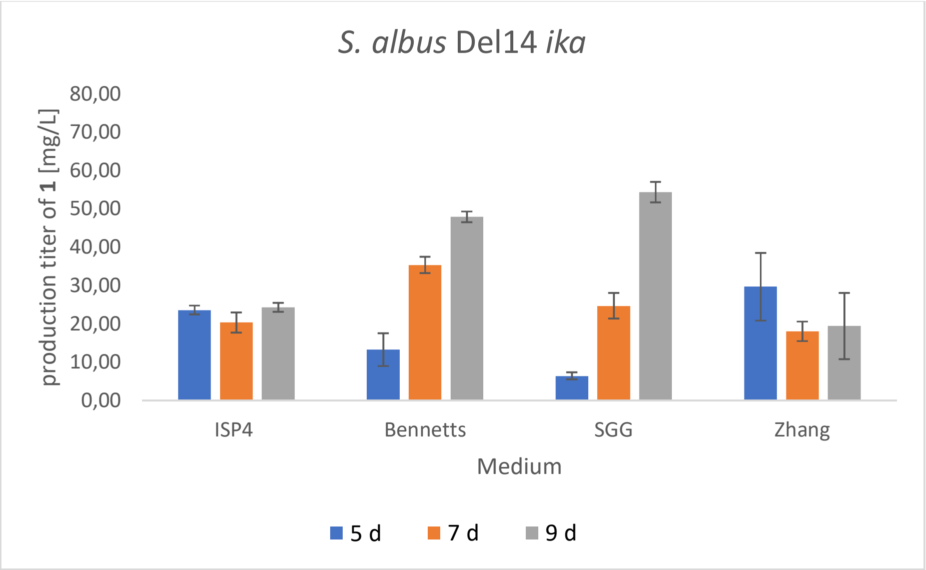

**Figure S191.** Production titer of **1** in *S. albus* Del14.

Scale-up fermentation experiments from 50 mL to 200 mL for three different best-performing strains in the best medium for each strain, as determined above. Extraction after the optimized time for each strain. Titers were determined using biological triplicates.

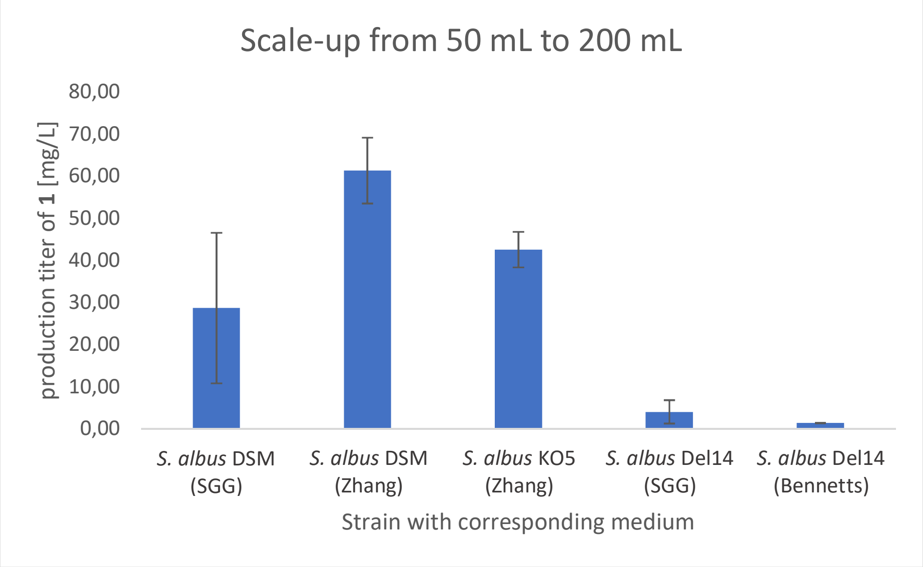

**Figure S20.** Production titer of **1** in three strains in their respective optimized medium.

Determination of the best inoculation ratios for the two best-producing strains. Titers were determined in biological triplicates.

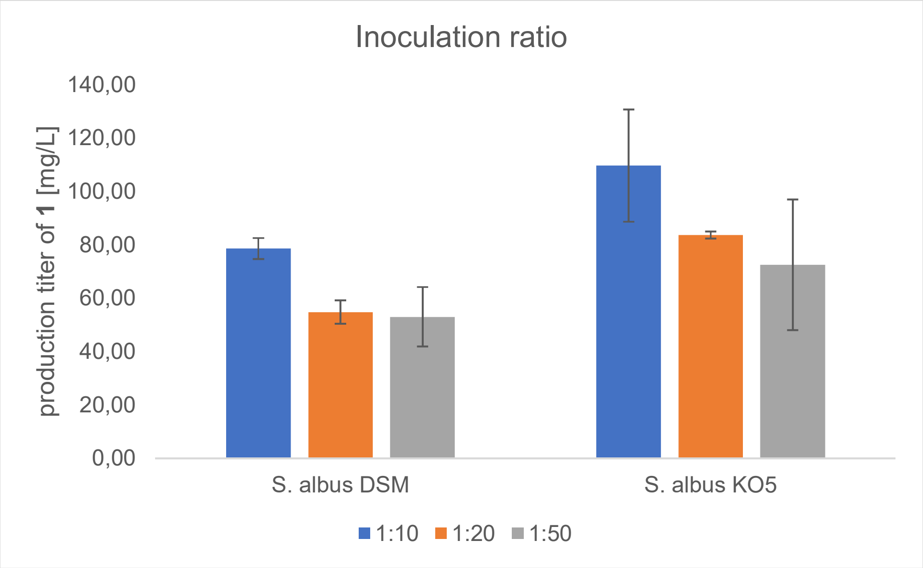

**Figure S21.** Production titer of **1** using best-producing strains for different inoculation ratios of pre- versus main culture.

#### 6 Characterization of the Isolated Ikarugamycin (1)

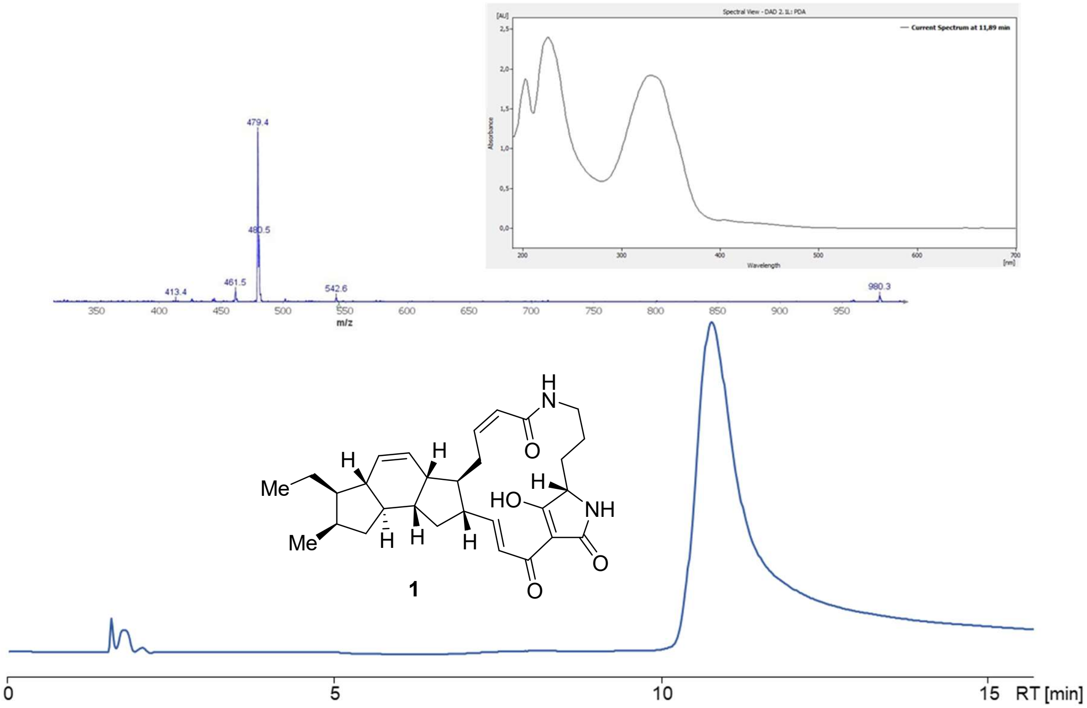

**Figure S22.** HPLC chromatogram of purified **1** (bottom) with MS trace (top; *m/z* = 479.4 [M+H]$^{+}$) and the corresponding UV spectrum (from DAD-HPLC; top right).

#### NMR data

**$^{1}$H-NMR** (600 MHz, DMSO-$d_6$)**:** δ [ppm] = 8.69 (bs, 1H), 7.87 (t, *J* = 5.6 Hz, 1H), 6.98 (d, *J* = 15.5 Hz, 1H), 6.64 (dd, *J* = 15.5, 10.2 Hz, 1H), 5.98 (td, *J* = 11.1, 3.8 Hz, 1H), 5.91 (bdt, *J* = 9.9, 2.8 Hz, 1H), 5.76 (bd, *J* = 11.2 Hz, 1H), 5.73 (dt, *J* = 9.9, 2.8 Hz, 1H), 3.83 (dd, *J* = 5.8, 2.0 Hz, 1H), 3.57–3.51 (m, 1H), 3.26–3.19 (m, 1H), 2.53–2.50 (m, 1H)*, 2.48–2.42 (m, 1H), 2.41–2.33 (m, 1H), 2.27–2.20 (m, 2H), 2.12–2.06 (m, 1H), 2.06–2.00 (m, 2H), 1.88–1.80 (m, 1H), 1.75–1.69 (m, 1H), 1.56–1.43 (m, 3H), 1.38–1.29 (m, 3H), 1.23–1.20 (m, 1H), 1.17–1.12 (m, 1H), 1.10–1.03 (m, 1H), 0.91 (t, *J* = 7.1 Hz, 3H), 0.86 (d, *J* = 7.1 Hz, 3H), 0.66 (ddd, *J* = 12.0, 12.0, 6.9 Hz, 1H).

*Overlap with solvent residual peak.

**$^{13}$C{$^{1}$H}-NMR** (150.9 MHz, DMSO-$d_6$): δ [ppm] = 195.8, 175.2, 171.4**, 165.5, 150.3*, 139.3, 130.6, 128.8, 124.3, 122.0*, 100.8, 61.1, 49.6, 48.3, 48.2, 46.7, 46.5, 42.6, 41.2, 38.2, 38.1, 36.1, 32.5, 26.8, 24.8, 21.2, 20.4, 17.7, 13.2.

*Taken from HSQC. ** Taken from HMBC.

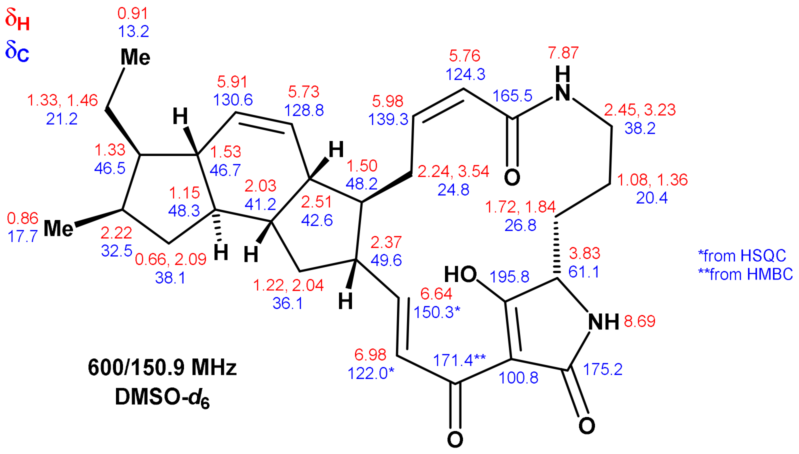

**Figure S23.** Complete $^{1}$H- (red) /$^{13}$C- (blue) NMR signal annotation of ikarugamycin (**1**) as determined by 2D NMR. For $^{1}$H multiplets, signal center is given.

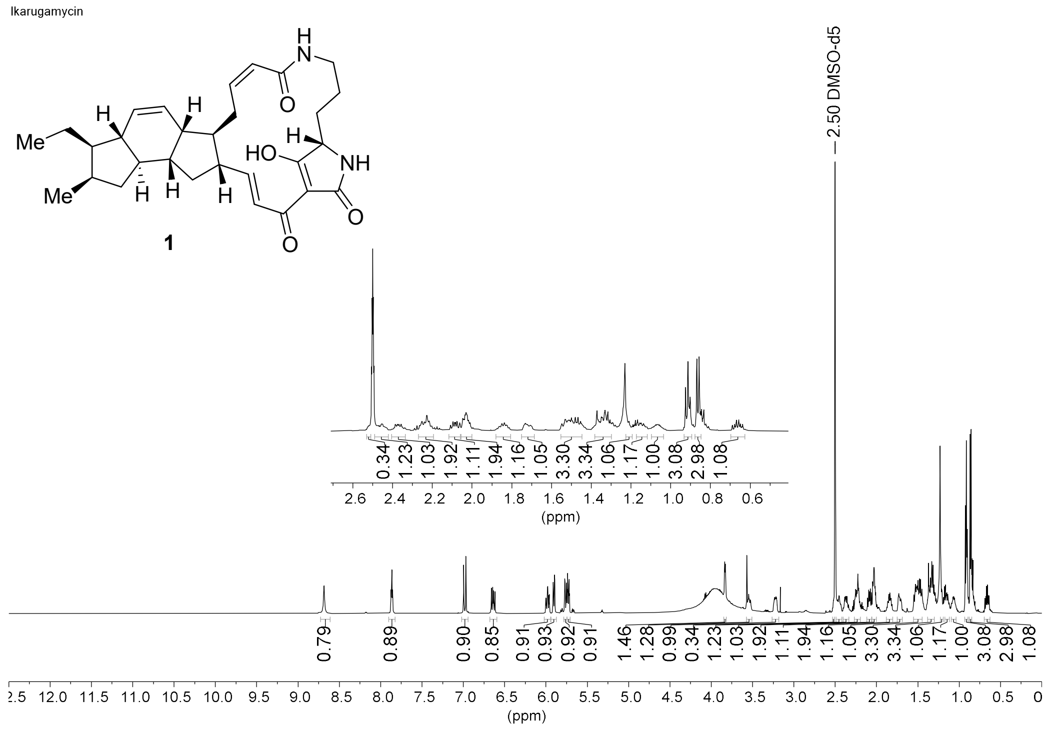

**Figure S24.** $^{1}$H-NMR spectrum (600 MHz, DMSO-$d_6$) of purified ikarugamycin (**1**) obtained by recombinant production.

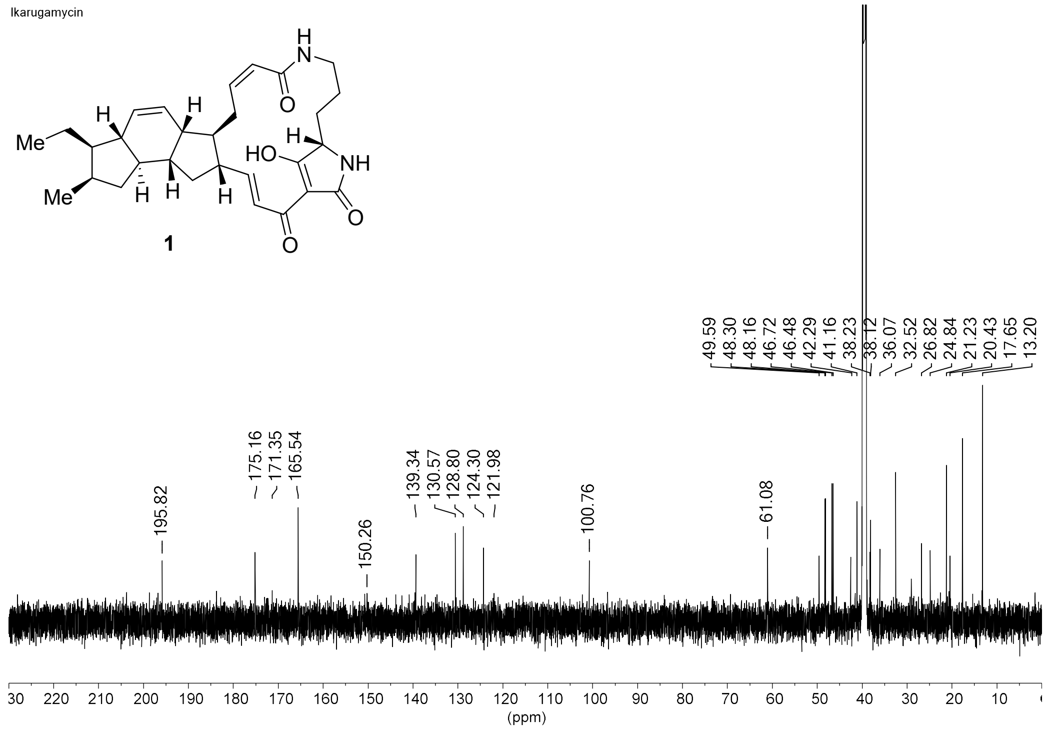

**Figure S25.** $^{13}$C{$^{1}$H} NMR spectrum (151 MHz, DMSO-$d_6$) of purified ikarugamycin (**1**) obtained by recombinant production.

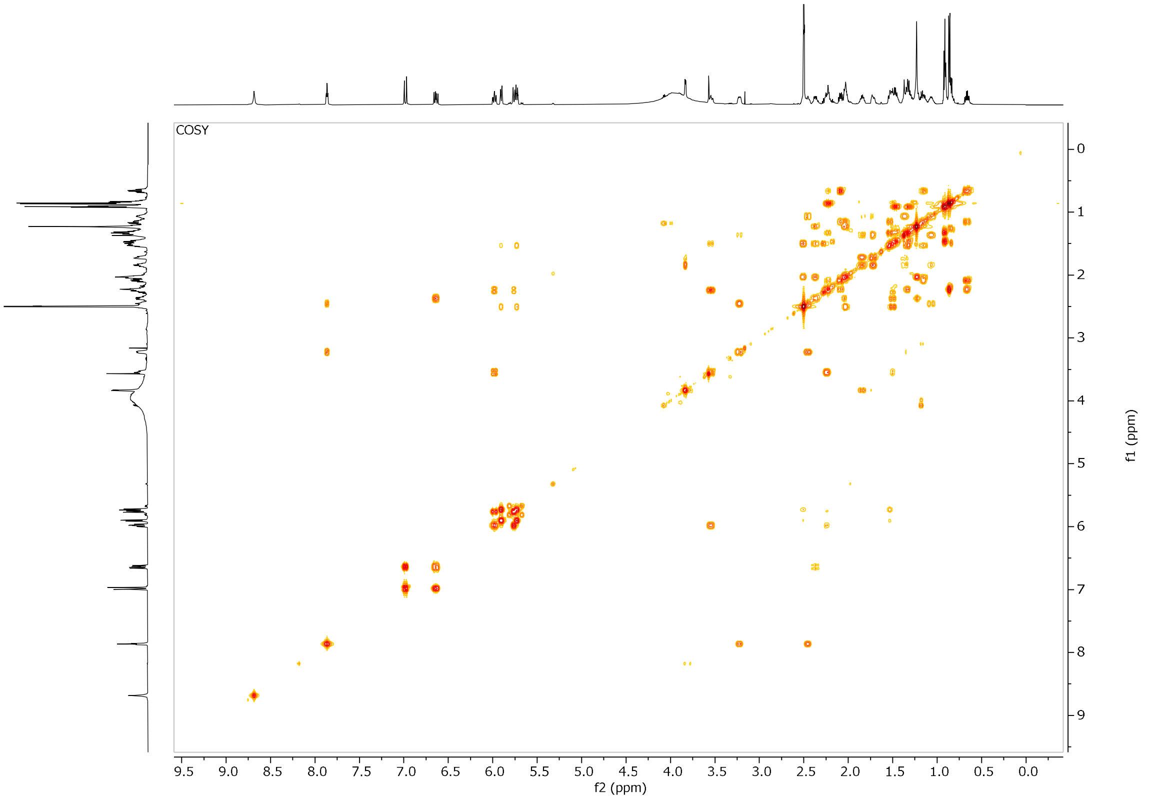

**Figure S26.** $^1$H–$^1$H-COSY NMR spectrum (DMSO-$d_6$) of purified ikarugamycin (**1**) obtained by recombinant production.

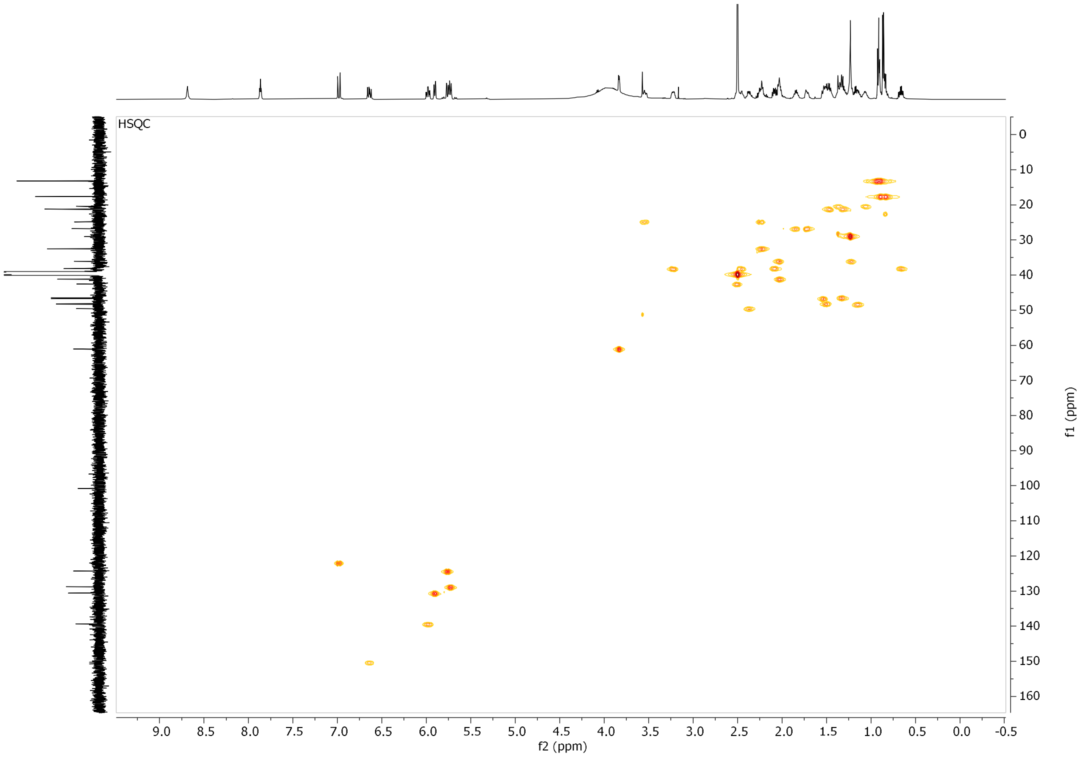

**Figure S27.** $^1$H–$^{13}$C-HSQC NMR spectrum (DMSO-$d_6$) of purified ikarugamycin (**1**) obtained by recombinant production.

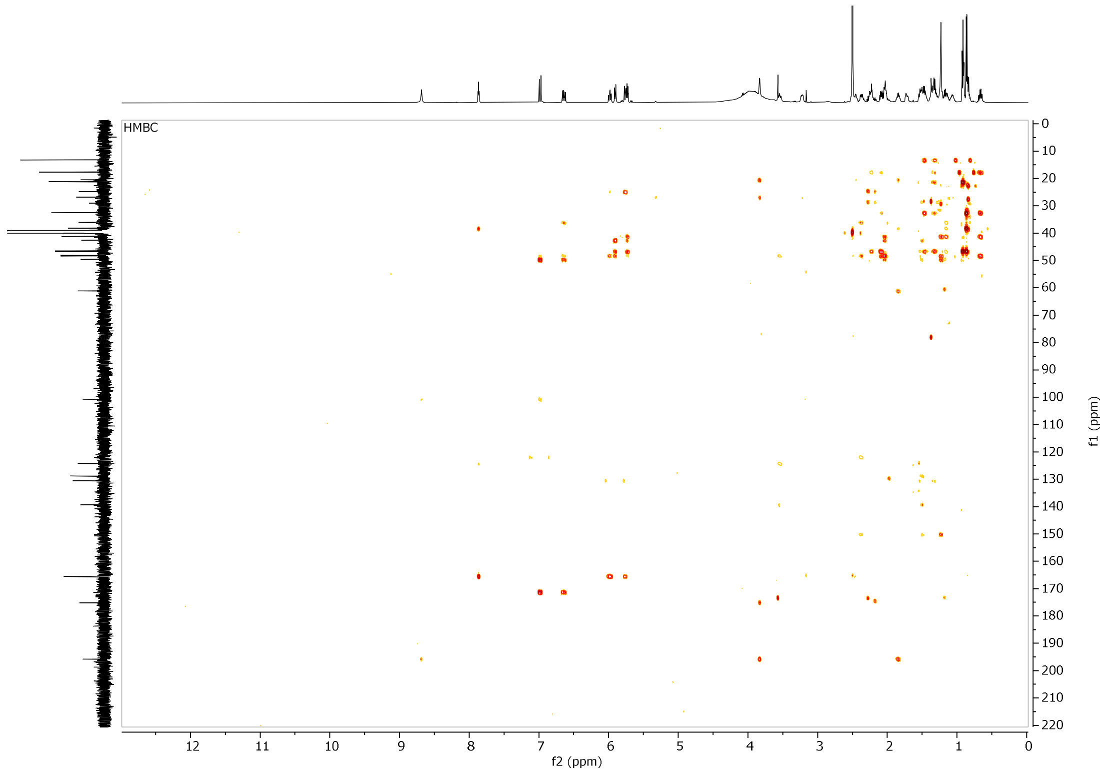

**Figure S28.** $^{1}H$–$^{13}C$-HMBC NMR spectrum (DMSO-$d_6$) of purified ikarugamycin (**1**) obtained by recombinant production.

#### 7 Supplementary Data of Employed Expression Constructs

##### 7.1 pSET152_ermE::*ika*

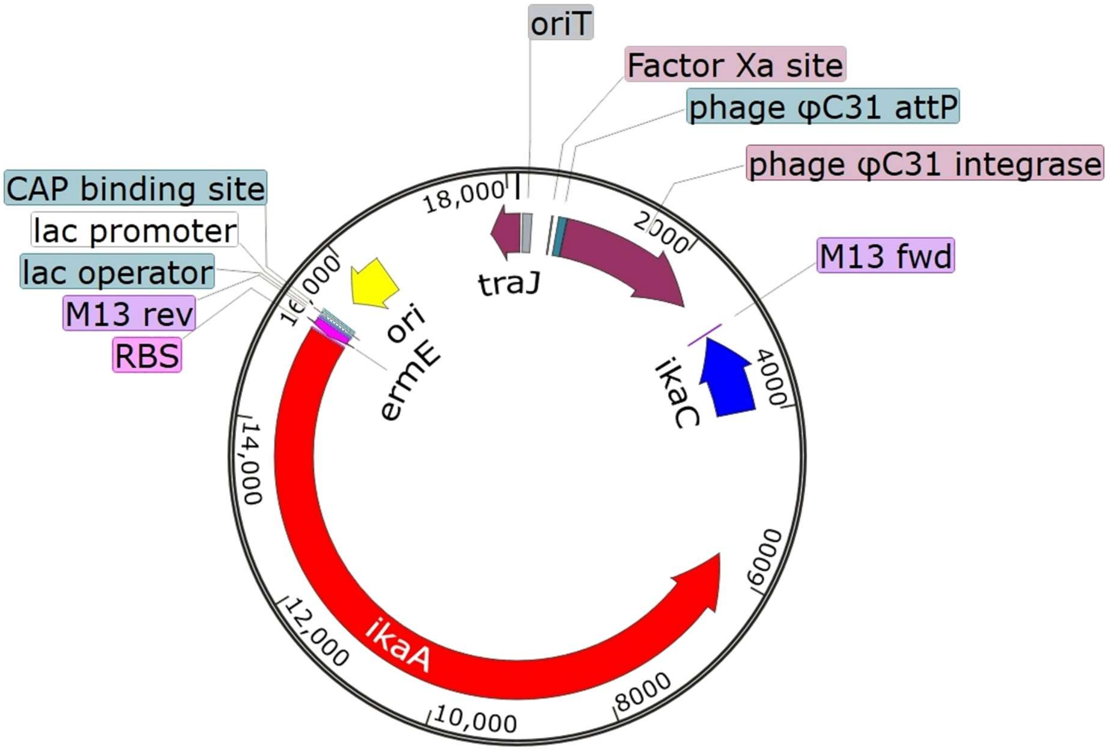

**Figure S2.** Vector map of the expression construct of *ika* in pSET152_ermE.

- DNA sequence of pSET 152_ermE::*ika* [18109 bp]:

CTGCCCTTCCTGGTTGGCTTGGTTTCATCAGCCATCCGCTTGCCCTCATCTGTTACGCCGGCGGTAGCCGGCCAG
CCTCGCAGAGCAGGATTCCCGTTGAGCACCGCCAGGTGCGAATAAGGGACAGTGAAGAAGGAACACCCGCTCGCG
GGTGGGCCTACTTCACCTATCCTGCCCGGCTGACGCCGTTGGATACACCAAGGAAAGTCTACACGAACCCTTTGG
CAAAATCCTGTATATCGTGCGAAAAAGGATGGATATACCGAAAAAATCGCTATAATGACCCCGAAGCAGGGTTAT
GCAGCGGAAAAGATCCGTCGACCTGCAGGCATGCAAGCTCTAGCGATTCCAGACGTCCCGAAGGCGTGGCGCGGC
TTCCCCGTGCCGGAGCAATCGCCCTGGGTGGGTTACACGACGCCCCTCTATGGCCCGTACTGACGGACACACCGA
AGCCCCGGCGGCAACCCTCAGCGGATGCCCCGGGGCTTCACGTTTTCCCAGGTCAGAAGCGGTTTTCGGGAGTAG
TGCCCCAACTGGGGTAACCTTTGAGTTCTCTCAGTTGGGGGCGTAGGGTCGCCGACATGACACAAGGGGTTGTGA
CCGGGGTGGACACGTACGCGGGTGCTTACGACCGTCAGTCGCGCGAGCGCGAAAATTCGAGCGCAGCAAGCCCAG
CGACACAGCGTAGCGCCAACGAAGACAAGGCGGCCGACCTTCAGCGCGAAGTCGAGCGCGACGGGGGCCGGTTCA
GGTTCGTCGGGCATTTCAGCGAAGCGCCGGGCACGTCGGCGTTCGGGACGGCGGAGCGCCCGGAGTTCGAACGCA
TCCTGAACGAATGCCGCGCCGGGCGGCTCAACATGATCATTGTCTATGACGTGTCGCGCTTCTCGCGCCTGAAGG
TCATGGACGCGATTCCGATTGTCTCGGAATTGCTCGCCCTGGGCGTGACGATTGTTTCCACTCAGGAAGGCGTCT
TCCGGCAGGGAAACGTCATGGACCTGATTCACCTGATTATGCGGCTCGACGCGTCGCACAAAGAATCTTCGCTGA
AGTCGGCGAAGATTCTCGACACGAAGAACCTTCAGCGCGAATTGGGCGGGTACGTCGGCGGGAAGGCGCCTTACG
GCTTCGAGCTTGTTTCGGAGACGAAGGAGATCACGCGCAACGGCCGAATGGTCAATGTCGTCATCAACAAGCTTG
CGCACTCGACCACTCCCCTTACCGGACCCTTCGAGTTCGAGCCCGACGTAATCCGGTGGTGGTGGCGTGAGATCA
AGACGCACAAACACCTTCCCTTCAAGCCGGGCAGTCAAGCCGCCATTCACCCGGGCAGCATCACGGGGCTTTGTA
AGCGCATGGACGCTGACGCCGTGCCGACCCGGGGCGAGACGATTGGGAAGAAGACCGCTTCAAGCGCCTGGGACC
CGGCAACCGTTATGCGAATCCTTCGGGACCCGCGTATTGCGGGCTTCGCCGCTGAGGTGATCTACAAGAAGAAGC
CGGACGGCACGCCGACCACGAAGATTGAGGGTTACCGCATTCAGCGCGACCCGATCACGCTCCGGCCGGTCGAGC
TTGATTGCGGACCGATCATCGAGCCCGCTGAGTGGTATGAGCTTCAGGCGTGGTTGGACGGCAGGGGGCGCGGCA

AGGGGCTTTCCCGGGGGCAAGCCATTCTGTCCGCCATGGACAAGCTGTACTGCGAGTGTGGCGCCGTCATGACTT
CGAAGCGCGGGGAAGAATCGATCAAGGACTCTTACCGCTGCCGTCGCCGGAAGGTGGTCGACCCGTCCGCACCTG
GGCAGCACGAAGGCACGTGCAACGTCAGCATGGCGGCACTCGACAAGTTCGTTGCGGAACGCATCTTCAACAAGA
TCAGGCACGCCGAAGGCGACGAAGAGACGTTGGCGCTTCTGTGGGAAGCCGCCCGACGCTTCGGCAAGCTCACTG
AGGCGCCTGAGAAGAGCGGCGAACGGGCGAACCTTGTTGCGGAGCGCGCCGACGCCCTGAACGCCCTTGAAGAGC
TGTACGAAGACCGCGCGGCAGGCGCGTACGACGGACCCGTTGGCAGGAAGCACTTCCGGAAGCAACAGGCAGCGC
TGACGCTCCGGCAGCAAGGGGCGGAAGAGCGGCTTGCCGAACTTGAAGCCGCCGAAGCCCCGAAGCTTCCCCTTG
ACCAATGGTTCCCCGAAGACGCCGACGCTGACCCGACCGGCCCTAAGTCGTGGTGGGGGCGCGCGTCAGTAGACG
ACAAGCGCGTGTTCGTCGGGCTCTTCGTAGACAAGATCGTTGTCACGAAGTCGACTACGGGCAGGGGGCAGGGAA
CGCCCATCGAGAAGCGCGCTTCGATCACGTGGGCGAAGCCGCCGACCGACGACGACGAAGACGACGCCCAGGACG
GCACGGAAGACGTAGCGGCGTAGCGAGACACCCGGGAAGCCTGATCTACGTCTGTCGAGAAGTTTCTGATCGAAA
AGTTCGACAGCGTCTCCGACCTGATGCAGCTCTCGGAGGGCGAAGAATCTCGTGCTTTCAGCTTCGATGTAGGAG
GGCGTGGATATGTCCTGCGGGTAAATAGCTGCGCCGATGGTTTCTACAAAGATCGTTATGTTTATCGGCACTTTG
CATCGGCCGCGCTCCCGATTCCGGAAGTGCTTGACATTGGGGAATTTATGCGGTGTGAAATACCGCACAGATGCG
TAAGGAGAAAATACCGCATCAGGCGCCATTCGCCATTCAGGCTGCGCAACTGTTGGGAAGGGCGATCGGTGCGGG
CCTCTTCGCTATTACGCCAGCTGGCGAAAGGGGGATGTGCTGCAAGGCGATTAAGTTGGGTAACGCCAGGGTTTT
CCCAGTCACGACGTTGTAAAACGACGGCCAGTGCCAAGCTTGGGCTGCAGGTCGACTCTAGAGGCCTCTACAG
GGCGACCAGGACCTTGCCGCGGTTGCCCTCCTGGCCGGTGTACAGGGAGCGGTACGCGTCCGGGAGGGCGTCGAA
GCCCTGGTGGATGGTCTGGTGGTAGGCCAGCTCACCCTTGCGGATCAGCCCGCCCACCTCCTCGTGCAGCGCGTC
GACCATCTCCTCCGTGTACCACTCGTCGGCGAAGATCCCCCGGATCGTGGTGCGCGGGAACATGATGTACGGCAG
CAGCCGCGGCCCCGTCCAGTCCCCGTTGACCGTGGTGGCCCACTGCCAGCACACCGCCACCTGGGAGTGCACGTT
GAGCATCGTGAACACCGCGTCCGTCACGGTGCCGCCCAGGTTGTCGAAGTACTTGTCGATTCCGTTGGGCGCTGC
CGCCGCCAGCGCCTCGCGCACCGTGTCCGTGTCGTCGCCCTGCCGGTAGTTCACCACCGCGTCGAAGCCCAGCTG
CGTCAGATACGCGGCCTTCCCCGGCGAGGAGGTGGTGCCCACCACCCGGGCCCCGGCCCGCTTGGCGAGCTGGCC
CACCAGGGTGCCCACCGTCCCGGACGCCCCGCTGATCACCACCGTGTCCTCGGAGCCCACGGTCAGGAACGTCTT
CATCGCGCCGAACGCGGTGATGCCCGGGGTGCCCATCACGCCCAGCGCCGTGGACAGCGGCAGCGCCTCGTCGTA
CCGCCGCGGGTCCAGCTTGCGGAACGGCGGCAACTGGACGGCGAACTCGGCGCCGTCGTCGTTCCATCCGCTGGG
CCCCCCGGTGGACACCAGATGGCTGCACCAGCCGCCGTACCCCTGCACCAGGTCCCCGACCTGGAAGCGGGCGCG
CGGCCCCGCCACGGCCACCTCCATGATCGAGTCACCGCGCACGGTGTCCCCGATCGGCGTCTGGAGCGAGAGCCC
CACCAGATACGGGTCCACCGACACGTACCGGGTGCGCAGCAGCATCTGATCGTCCGCCAGCGAGGCGGGATCGAA
CTCCCGCGTCACCTTCCGGTAGATCCGGTCCGTGTCCGGGACCCCCGGGATGTGCTCGGCGATCACCCATGTGTG
TACCTGCACTACACGCTCCTCGACGATGTGACGGTGTTGTTGGCGCCGGTGGCGGTCTCGCCGCCGGCGGTCTGG
GCGGGGAACCCGGCCGCCAGATCCGGCACGCCCGGCTGCTTGCCCACCGGGATGGCGACCTGGGTGGGCGGCGGC
GCGCTCTCGTCCACGCTCGCCGTGAACTCCCGCCCGTCGTCCCGGCAGATCACCTGCGTGACCTGCCGGCCCGAG
GCCACCGCCCGGATCAGCCCGCCGACGGTGACCCACACCCCGGCCAGATAGAAGTTGGAGAGCCCCGGCAGCACC
GGGCCGTTCTTGTTGATCTCCACCTCCACGGTCTCCCCGCCGTCCACGAACGGCTGCCAGCCGGGGAACCCGCCG
TTGTAGGTGCCCGTGTAGCGCACCTGCGTCAGCGGCGTGGACACGTCCCGCACGGCGACCGCGTCCTTGAGACCG
GGGAACCGCTCGTCCAGGAAGTTCTCGATGGTGATCCGCGCCTGCCGCTTGGCCTGGGTGTACGCCTTGCCGTGC
TTCACCGGCAGGGTGTGCAGCACCTGACCGCGCCGCACCCGGCCCGCCTGTTCCGGCACGTCGTCGCGCAGCGCC
CGCCACGGCTCGGCCTCCGAGAAGTACGTGGCGAAGATGACCGTGGTCTCGCGCGGCGACAGCTCCGGGTAGTGG
CAGCTGCGGAACTGCACGTTCATGCTGGGATGCCGCATGCCGGTGAGCTTCTCCGCCATGCTGTCCTCCAGCACG
TACGTGGTGCACGGCTCGCCCTCGGGGAACGGCCGGCGCAGCCCCAGGAACAGCGAGACATAGCCGGGGGAGATC
GTGCCCACCTCGTCGATCGTCTCGGTGAGCAGCTTGCGCCAGGTGTCGTTGAGATACCGGCCGCCGAGCATCTCC
ATGGCGGTGGTGTGCAGATCGGCCGCCGACACCACGATGTCCGCGCGGAACTCGCGGCCGTCGGTGAGCCGCACT
CCCACCGCCTTGTCGTGCTCGACGAGGATCTTCTCCACCTTGGCGTTGTAGGTGATCTCCCCGCCGAGCCCCAGG
TAGCGCCGCTCCACGGACCGGGCCAGCTCCAGCGAGCCGCCCTCGGGCACCCCCGCCGAGCCGTTGGCGTGCGAG
GCCAGCTGGAACCAGAACGGCAGGACGGGGAAGTCGGCGTGCTTCTCGTACAGCACGTAGTTGAAGGCCTCGCGC
AGCACCGGGTGCTGGAACTTCTCCGCGTAGTCCGTCATCAGCTCGGTGATGGACTTGCGGATGGCGTTGAAGTAC
GGCAGGAACGAGGCCAGCATCTTCCACCGTTCCCACCGCCCCATCAGCCCCACCGGCTTGAGGAACGGGTAGACC
GACAGCGCCTTCTGGAAGGTGCGCACCCCCTCGCAGAAGTTCTTGATGCGGCGGGCGTCGGCCGGGGAGATCTCC
AGCAGGTGCGCCTGGAGCCGGTCCGGGTCGGAGTAGAAGTACACCGGCTGGCCGCCGCGCACCCGCACGATGTTG
AAGACGTCGAACTGGCGCATCTCCTTGCCCTGCAACGCCCCCAGTTCCATCCAGATCTGGTACATCTCGTTGCCG
GGACCGCTGCCCAGCAGCCAGCTGACGCACCAGTCGAAGGTGAAGTCCCCGCGCTCCCAGGCGGTGCAGGAACCG
CCCGGGATCTCGTGCATCTCGAAGACCCGCGTCGCGTAGCCGTTCATCTGCGCGTAGCAGCCGGTGGACAGGCCC

CCCAGGCCGCCGCCGATGATGATCATCGACTGCCTGCCCGGGGTGCCGGAGGTGGTGGGGGATGACATGGCGCTG
TGCTCCTTGGTGTCGACCGCCGGCTGAACGAAAGGCGTCATACCTCGCCATCACCGCCGGTGAGAATGCCGCGCG
CCAGGCGCGCGTTGTGCTCGACCGTCCCCGGATCGAGCATGTCGGCGTGCCGGCCCACCCCGCGCAGCACGTCGG
TTCCGGTACGGGAGGCGCCGTGCCAGCTGCCCTCGGCGCCGCTGTCGTACGCGGCCGCGTTCTCCTCGTCACTGA
TCACCGCGATCCGGGCCCCGGTGGTGCCGGGGTTGGCGGTGCGGCCGGTGAACTCCAGGTAGTCCTTGGCCTGTT
CGCGCGTCTTCTCCGCGACCAGCGCCGAGCCGGTGTGCTTGCGCAGATGCTCGGCCAGCTCGCGCTCGAAGACGG
CCAGGTGCTCCGGGCCCAGCTCGTAGGACTCCGTCACCCGCAGCGAGTCCATGATGACGACGTGGCGCACGGTGC
GGCCGCGCCGCTCCAGCTCCTTGGCCACCTCGAAGGCGAGGTTGCCGCCCAGCGAGTAGCCGAGCAGGTCGATCT
CGCCCTCCGGCCGGTGCCCGGCCACCAGGTCGGCGTACCCGCTTACCTTGTCCTCGCCCATCAGGTAGTTGAAGG
CGAGGAACTCGAACTCCGGCAGCCGCGCCGCGAACTCCCGGTAGACCAGGCCGTGGCCACCGGCCGGCGGGAAGC
AGAACACCGTGCCCGCCGCGGCGTTCTCGTTGAACCGCAGATACGGCAGCGACCCCTCGATCTCCCCGGTGACGA
TCCGCTCGACCGTCTTGGCCATGCCGTGCAGCGTCGTCACCTGGAACAGCCGGCTGACCGGGATGGAGATCCCGA
ACTCGGCCTGCAGGTGGTAGATCAGCTCGATCAGCCGGATGGAGCTGCCACCGGTCTCGAAGAAGTCGTGGCCCA
GACCGGGCCCGGGGGCCTCGATGCCCAGCAGCCGCTGCCAGTGCTCGGCCATCCGGGTCTCGTACAGCGTGACGG
GGGCCTCGTACACCGGCCCGTCCGCGCCGTCGCCGGTGCGCGGGGCGGGCAGCGCCGCCACGTCCACCTTGCCGT
TCGGGGTGAGCGGCAGGGCGGGCAGCTCGGTGAAGTGGGTGGGGATCATGAACGTCGGCAGATAGTCCGCCAGGC
GCCGGCGCACCTCGCGCCAGTCCAGCACGGCGCCCGGGGCCGCCACCCCGTACGCGCACAGCACGTTCTCGCCGC
CCGCGTCCGGCCGCACGGTGACCTGCGCCTGGGCCAGCTCGGGGCAGGCCGCCAGGTGCGACTCGATCTCCCCGA
TCTCGATGCGGTGCCCGCGCACCTTGATCTGCGAGTCGGCCCGGCCCAGCAGATGGACGGTGCCGGCCGCGTCCC
AGCGCGCGAGGTCACCGGTGCGGTACAGCCGTACCGGAGCGTCGCCGGCCAGGGCGCGGGTGAGGAACCGCTCCC
CGGTCAGCGCCTCGTCGCCGAGGTAGCCGAGCGCCACCCCGGTGCCGCCGATCCACAGCTCGCCGGGGACACCGG
GCGGCACCGGCTCGCCGCGCGAGTCCAGGATGTACAGCGCGCTGCCCGGGAACGGCCGCCCGATGGGCACCATCC
GGCCGCCCTCCAGGTCATCCGCGGGACCCTCGAACCAGGCGCTGTCGATGGTGGCCTCGGTGAGCCCGTACGAGT
TCACCAGGCGCTGTGCGCCCAGCGCGCGCAGCCGCTCGTACTCCTCGGCCTTCCAGGAGTCCGAGCCCACGATCA
GCAGCCGCAGGAAGTCCAGCCGGGCGCCGGTGTCCTCGCAGTGCCGCACCAGGGTGCGCACCACGGCGGGCACGA
ACTCGCCGCAGTCCACCCGTTCGGCGCGCATCGTCTCGTACAGCCGGGCGGTGTTGAACAGCAGCTCCCGGCCGA
CCAGCACCAGCGTGCCGCCCGAGCACAGGGCTCGGGTCAGGTCGCCGGTGAAGACGTCGAAGGAGGGGCTGGCCA
TCTGGAGATGGACCCGGATGCCGCCCTCCTCCAGGCGGTAGGCGTCGCGCCATCCGGCGTACACCGAGGCCAGAT
TGCGGTGGCTGACCGCGACCGCCTTGGGGCGCCCGGTGGAGCCCGAGGTGTAGATGACGTAGGCGGGGGAGTCGG
GCCCCGGCTCGGCGTCCGGCCCCGCTTCGCCGGCCTCGCCCGCGAGCAGTTCCTCCAGGGTGACCACGGTGCCCG
GCAGCCCCTCGGCCGCGCCGCCCGTCCCGCCGATCACCAGCGTGGCCCCGGCGTTGCGCACCATGTACGCGAGCC
GGTCGGCCGGATAGTCCGGGTCCAGCGGCAGATAGGCGCCGCCCGCCTTGAGGACCGCCAGCAGGGCGGTGATCA
GCTCGGGCGACTTCTCCAGGCACAGCGCGACGACGGTGCCCTCGCGCACGCCGCGCGCCCGCAGCCGCCCGGCCA
GTTCGCCGGCGCGCTCCTCCAGTTCGCCGTACGTCAGCCGCCGGGTCTCCCCGCTCTCGGCGGGCGCGGCCACCG
CGATCGCCTGCGGGGTACGGTGCGCCGCCTCGGCGATCAGCCGGTGCACCGGCACCGGCGCGTCCTGCGCGCCCT
GCCCGGCCCCGCTCCACTCGGTGAGGATCCGCTCCCGCTCGCCGCCCGACAGCATCCGCAGTCCACCGGTGGCGG
CGTCGGCGGGCGCCGCGGTCAGCGACTCCAGGAGCTGCGTGTAGTGCCCGGCGAGCCGCCGGATCGTCTCCGCCT
CGAAGAGGTCGGTGTTGTACTTGAAGACGCAGTGGAACCGCCCGTCCGCCTCCTCCTCGTACGCGGACAGCGTCA
GGTCGAACTGGCCCTCCTCCTCGGGCAGCTCGATGTACTCCAGCTTGTAGCCGTACTTCTCGGTGGCCACCTTGT
GGTGCAGCAGGATGAACATCGCCTGGAAGACCGCCGACCGGCTCGGGTCGTGGGCCAGCCCCAGCTGCTCCACCA
GCAGCGTGAACGGGTACTCCTGGTGGTCCAGGCCGCCCAGCACCGTGGTGCGCACCTGGTCCAGCAGCTCGGCGA
CCGTGGGGTCACCGGCCAGCGAGGCGTGCAGCGGCAGCGGGTTCACGAAGTACCCGTAGACGGCGCCGAACTCCT
CCTGGGTGCGGCCGGTGACGGGGGAGCCGACGATGATGTCGTCCTGCCCCGCATAGCGGTGCAGCAGCAGGTAGT
ACGCGCTCAGCAGCACCATGAAGACGGTGACGTTGTGCTCCCGCGCCAGCGCGTGCACCCGGGCGCTCAACTCCG
CGTCCAGGGCGAAGAACTCGGACGCCCCGTTGTGGGTGAGCACCGCCGGGCGCGGCTTGTCGGTGGGCAGCGCCA
GCACCGGCACCTCGTCCGGCAGCTGCCCCCGCCAGTACGCGAGCATCTTCTGCGCCTCGCGGCCGGCCAGGAACG
CGTTCTGCCAGTTGAGGAAGTCCAGATAGCGGGCGGACACCGGCGGCAGTTCGACGTCGTGGCCCTGCCGCAGCC
CCTCGTACAGGGACAGCAGTTCCTCGATGAAGGTGAAGGTGGAGATGGCGTCCGAGATGATGTGGTGGACGGCCT
TGGTGATGACCCAGCGGTCCGGGCCGCGCCGGAAGAGGCGGAACCGGATCAGCGGATCGGTCGCCAGGTCGTACG
GCTTGCGGTACTCCTCGATGATCATCCGGTAGATGTCGTCCCACGCGCGGTCCTCGACGTCGAAGAGCGCGATGT
CCTCCTTGATCTCCGGGGAGATCCGCTGCACCGCCTGCCCCTCCACCAGCAGGAAGTTCGCCCGCAGCACGGGAT
GCCGGGCCAGCAGCCGGCGAAACGCCTCGAACATCAGGTCCGGGTCCAGCTCGACCCGCACCTCGACGGCGCCGC
CGATGTTGTACGCGAAGCCGTCCGGGTTCAGCTGCTTCAGGAACCACAGCGCCTTCTGGTTCTGCGTCAGCGGGA
ACTCGGCCTCGTCCTCGAAGCGCTCCACCGCCGTCACCGCGGCGCCGCTGCCCTCCTCGGCCAGCAACTCCTCCA

GGCCCTCGTGCAGCTGGGTGATCAGGTCCCCGGCCGGCGCGCTGGACAGCAGCGCCACCACCGGCAGCGCGATGC
CCACCTCGGCCACCACCCGCGCTCGCAGCTCCATCGCCAGCAGCGAGTCGAGCCCCAGCAGATTCAGGCTGACCG
CCGGATCCACCTGCTCGGCCCGCACCCGCAGCACACCCGCCACCAGCGTCGTGAACCGCTCGGTCAGCAGCAGCC
GCCGCTTGTCCGCGGTGGCCTCCCGGAACGCGTCCAGGAAACTGCCGTCGCCCTCGGACCCCGGTCCCTGGGCGG
TGGCCGCCAGCTCCGTGACCAGCCGCGGCGGCGCCGCGTACCAGGACATGAACACCGGCCAGTCCACGACCGTGG
CCACCAGCAGCTGTGCCCGGTCCTGCCCGATGACCCGCTCCAGCACCGCCATGCCCGCCTCGGGCGCCAGCGAGG
ACATGCCCCGGCTGTTGCGGTAGTGGTCGATCAGGCCCAGTTCCTCGATCATGCCGGTGGCCCACGGGCCCCAGT
CCAGCGCCAGCGCCGGCAGCCCCTGCGCGCGGCGGTGGTGCGCCAGCGCGTCCAGGAAGGCGTTCCCCGCCGCGT
AGTTGGTCTGTCCGGCCGTCGTCAGCCAGGCCGCGACCGAGGCGAACAGCACGAAGTGCTCCAGCGGTTCGCCGC
TCAGCTGCCGGTGCAGCAGCGCCGCGCCCACCACCTTCGGGTCGTGGACGGCGTCGAACACCTCCCGGTCCATCT
CCGGCACCAGGGTGTCGCGCACCTGCCCCGCCAGATGGAACACCCCGCGGATCGGCGGCCCCTGGGCGCGCCGGT
ACCCGGCGAGCCAGCCGGCCAGCGCGTCCTCGTCGGTGATGTCCAGCGGCGCGAGAATCGGCTGCGCGCCCAGCG
CCTCCAGCTCCTTGAGGAAGGCCACGTGCCGCCCGGCCGGCGAGTTCGGGTCCTGGTCGGCCCAGCGCTCGCGCT
CCGGCAGCCGGGTGCGGCCCACCAGGATCAGCCGCCGCGCCCCGCGCCTGACCAGCGTGCGGCACAGCAGCCTGC
CGAGCGCGCCGAACGCGCCGGTCACCAGATAGCTGCCGTCCGGGCGCAGCCGCAGGGGCAGCGGCCTGCTCAGCC
CCTCGGCGGCCACCAGCCGGCTGGTGTGCCGCTCCCCGGCGCGCAGCGCGATCTCGTCCTCGCGCTCGGCCGCGC
CGGTGGGGTCGGCCAGCTCGCGCAGCAGCGCGTACGCGTCCTCCTCGACGCCGCCGTCGGCCGCCAGGTCGATCA
GCTTGCCGCCGCGCCCGGCCAGTTCCTGCTGCCACAGCACCCGGCCGATGCCCCAGGCGGGCGCGCCCAGCGGCT
CCACCGGCTCACCGGGGACCACGCACTGGGCGGCTCTGGTGACGATGTGCACCGGGGTGCCGCCGTGCCGCTCCG
GGTCGGCGAGCAGGGCCTGAGTGAGGGCGATCAGGGCGTACGCGCCGGTGGAGGCGATGTCCGCGAACCGTCCGC
GCGGGGCGTCGGCCAGCGCCGGCCGGTCCAGGTTCCACAGGTGGACGACGCCGTCCACCTGCCCGAGATCGGTGA
GCAACCGCCGCAGGTCATCCGCGGATCCGGGGACGACGGTGGCGGTCCTCGCCGTGCGGTCCAGGCCGTACGCGG
CACCGGGCCGGACCAGATGGGCCTCCCCGCCGGCCTCGCCGATCAGCGCGGCCAGCCGCTGCGCGACCCCGCCCG
CGTCGGCGAACAGTACGTGCCGCCCGGCCGCCGGCGCGGCGGACGCCGCCTCGGGCAGCGGGCACGGCACCCAGC
TCGGTTCGGTGAGCCAGCTGTCGATGGTGGACAGCCCCACGGTGGTGGCGGCCTTCTCCACATCGGCGGCGCGGA
AGCCGGCGACGCGGCCCAGCGGCGTACCGTCGGCGCCGTCGTACACGGCGATGTCACCGGTGAGTTCGTCCTCGT
CGTCGCCGGTGACGGTGGCGTGCACCCACAGTTCGCGGTCGCCGACCGGGTCCAGCCGTACCTCGGCGATGGACA
GCGGCAGCCGGATGCCGGTGCCCCCGGGCCCGGCGGGCGCGGTGAGCAGCTGCGGGGTCAGCAGCGACTGGAAGC
AGGAGTCGAGCAGCACCGGATGCATGTGGTGCGCCGCCGCGTCCGGGGTGAGCCCCTGCGGCGGACGGATCCGGG
CCAGGGCCTCGCCCTCGCCGATCCACACCTCCTCGATGCCCTGGAAGGCGGGGCCGTAGTGGTAGCCGAGCGCGG
CCAGTTCGGCGTAGCAGTCGGGGCCGCTCAGGTGGCGGGCGGCGCGGGCCCGGACGGCGACGGTGTCCAGCGGCG
CGGTCAGCCGGCGGCGCTGGGCGGCCCGTACCGTACCGGTGGCGTGCACGGTCGGCTCGGCGCCGGCGGCGCCCA
CGGTGGCGATGGAGAACGCGGCGGCGTCGGAGGAGAAGGACAGCTGCACCGTCTGCGGCTCGCCGTCCGGCAGGA
ACAGCGCCTTGCGCAGCTCGATGCCGGCCAGCGCGGCGGTGCTGTGCTCATCACCGGTCAGCGCCCGCATGGCCT
GCGCGGCCATCTCCAGATAGCCGGCGGCCGGGAACAGCACGGTGCCCTGGATGCGGTGGTCCTCCAGGTACGGGG
CGGCCTCCGCGTCCAGCTTCACCTCCCACACCGGCTCGGCGCTCGCGGTGCGGCGGCCCAGCAGCGGGTGGTCGC
GGTGGCCGAGCCGGATCTGCGCGACCGGGGCCGGCTCGACCCAGTACCGGTCGCGCCGGAACGGGTAGCGCGGCA
GTTCGGCCGGCCGCCCGGCGGGGTGCAGGGCGTGCCAGTCCACGGCGAAGCCGAGGCTGTGCAGCGCGGCGAGCG
ACAGGGTGAGGCGTTCGCGCTCGTCCGCCTTGCGGCGGATGGAGGCCAGGGTGACGCTGTGCGCGTCGCGGGCCT
CGCAGCACTCGCGCAGGGAGTGGGCGAGCACGGGGTGCGGGCCGATCTCCAGGAAGACGCCGTAGCCGTCGTCCA
GCAGCCGGTCCACGGCGGCCCGGAAGTGCACGGCCTCGCGCACATTGCGCCACCAGTAGTCGGCGTCCAGCTCCG
TGCCCTGCGCGACGGTGCCCGGCAGCGCGGTGAGGTACAGCGGCAACTTCGCCTGCTGCGGCTTGAGATCGGCGA
GCGAGGTCAGCAGCTCGTCCTTGATCAGCTCCATGCGGGGGCTGTGGTACGGACGCCGACCTCCAGGAAGCGGG
CGAAGATGTCCTCGGCGCCCAGCTCGGCGGCGATCACCTCCAGCGCCTCGGTGTCCCCGGCCAGGGTGATGGAGG
TGGGGCTGTTGACGGCGGCGATGGAGACCCGGTCGCCGTGCGGGCGCACCCGGCGGGCGGCCTCCGCCTCGGTGA
GGCTGACGGCGAGCATGGAGCCGGTGCCGATGAGCTTCTGCTGGAGCCGGCTGCGGTGCACCACGATCTTCACGG
CCTCGGGGAGGGTGTACACCCCGGCCTCGTAGAACGCGGCGACCTCACCGGTGCTGTGCCCGGTGACGGCGTCGG
GCTGGATCCCCTTGCTGCGCCACAGGGCGGCCAGGGCGATCTGGACGGCGAAGTTGGCGGGCTGGGCGAGCCAGG
TCTCGCTCATCCGGGAGTCGGCCTCGTCGGCGTTCAGCTCCTGGGTGAGGGACCAGCCGGTGAGCGCGGCGATCT
CCTGGTCGCAGCGGTCGATGACCTCCCGGTAGACGGGCTCGCTCGCGTACAACTGGCGGCCCATGGCCCACCACT
GCGGGCCCATGCCGGTGAACACCCACACCAGCCTGCGGTCCAGGCCCTCCCGCTGGGTGCCGGCGACGGCACGCG
GGTGGCTCTCGCCGCGGGCGACGGCGCCGAGCACCTCGTCGAGGGCCTCGGGGGAGTCGTACACGACGGACAGCC
GCTGCTCCAGATGCTGCCGCCGGTGGGCGAGGGTGTAGGCGGCGTCGCCCACCGGCAGTCCCTCGGCGAGCCGTT
CGCGCAGGCCGGTGGCGATGGCGGGAAAGGCTTCGGGGTCGCGGGCGCTGAGCGGCAGCACGGAGTACCCGTCGG

TGGCCGGTGGCGCGGGCTCCCCGACGGTCGGCGGTGCCTCCTGGAGCAGGACGTGGGCGTTGGTCCCGCCGAAGC
CGAAGGAGTTGACGCCGGCCCGGGCCGGCCCGCTGTGCTCGGGCCAGGGGGTGGGCTCGCGGGGGATCTCGTACG
GCAGGGACGCCTCGTCGATCTGCGGGTTGAGCTTCTCCAGGTTGATGTGCGGCGGGATGACCTTGTGCTTGAGGC
TGAGCACCGTCTTGATCAGCCCGGCGATGCCGGCGGCGGACTCGGTGTGCCCGATGTTGGTCTTGACCGAGCCGA
CGTACGTCCGGGCGCCCGGCTCGCGGCCGATGGAGAGCGCGCGGCCGAGGGCGTTGGCCTCCAGCGGGTCGCCGA
CGGGGGTGGAGGTGCCGTGCGCCTCGACGTACTGGAGGCTGCCGGGGGTGACGCCGGCGGCGGCGCAGACCCGCT
CGATCAGGGCGACCTGCGCGTCGGGGTTGGGCACGGTGATGCCGTTGGTGCGGCCGTCCTGGTTGACGCCGCTGC
CGATGATGACGGCGTGGATGGGATCGCCGTCGCGCTGCGCGTCCGCGAGGCGCTTGATGGCGACCATGCCGACGC
CCTCGGCGCGCACGTAGCCGTTGGCGGAGGCGTCCAGGGCGCGGGAGCGGCCGTCGGGGGAGAGGAACCCGCCCT
TGGTCTCGGCGATGGTGTACTGCGGCGCCATGTGCAGCAGGGTGCCGCCGGCCAGGGCGACGGAGGTCTCGCCGC
GGCGCAGGCTCTGGCAGGCGAGGTGGACGGCGACCAGGGAGCCGCTGCACGCGGTGTCGACGGAGACCGAGGGTC
CGCGGAAGTCGAAGCAGTACGAGATCCGGTTGGACACCATCGTCATCATGGTGCCGGTGGCGGTGTGCGCGGCCA
GGGTCTCGAAGCCGAGGTCGGCGAACTGCAGGATCTTGTAGTCGAGGGTGAACGCCCCGACGTACACCCCGACAT
CGCTGCCGGCCAGCTCGGCGGGCTTGAGGCCGCCGTCCTCCAGCGCCTCCCAGGCGACCTCCAGGAGCTTCCGCT
GCTGGGGGTCCATGTGCTCGGCCTCGCGCGGGCTGATGCCGAAGAAGGCGGGGTCGAACTCGTCGAAGCCGTCGA
TGTATCCACCGCGTCCGCCGACCAGCCGGCCGGGCTTGGCCTTGTCGCCGCTGCCCAGGGTGCGGGTGTCGTAGC
GGTCGGCGGGGGTGTCGGTGATGCAGTCCTTGCCGTCGAGGAGGTTGCGCCAGAAGGTCCGGTAGTCGCTGGCGC
CGCCGGGCAGCCGGCAGCCGATGCCGACGATGGCGAACGCGTCGTCCTGGGACGGGACGGGCGCGGGGACTTCGG
GTACGGGGACGGGGGCAGGGTGGTGCATGGAATCCATGAATACATCCTTCCGTACCTCCGTTGCTCCGCTGGATC
CTACCAACCGGCACGATTGTGCCCACAACAGCATCGCGGTGCCACGTGTGGACCGCGTCGGTCAGATCCTCCCCG
CACCTCTCGCCAGCCGTCAAGATCGACCGCGAATTCGTAATCATGTCATAGCTGTTTCCTGTGTGAAATTGTTAT
CCGCTCACAATTCCACACAACATACGAGCCGGAAGCATAAAGTGTAAAGCCTGGGGTGCCTAATGAGTGAGCTAA
CTCACATTAATTGCGTTGCGCTCACTGCCCGCTTTCCAGTCGGGAAACCTGTCGTGCCAGCTGCATTAATGAATC
GGCCAACGCGCGGGGAGAGGCGGTTTGCGTATTGGGCGCTCTTCCGCTTCCTCGCTCACTGACTCGCTGCGCTCG
GTCGTTCGGCTGCGGCGAGCGGTATCAGCTCACTCAAAGGCGGTAATACGGTTATCCACAGAATCAGGGGATAAC
GCAGGAAAGAACATGTGAGCAAAAGGCCAGCAAAAGGCCAGGAACCGTAAAAAGGCCGCGTTGCTGGCGTTTTTC
CATAGGCTCCGCCCCCCTGACGAGCATCACAAAAATCGACGCTCAAGTCAGAGGTGGCGAAACCCGACAGGACTA
TAAAGATACCAGGCGTTTCCCCCTGGAAGCTCCCTCGTGCGCTCTCCTGTTCCGACCCTGCCGCTTACCGGATAC
CTGTCCGCCTTTCTCCCTTCGGGAAGCGTGGCGCTTTCTCATAGCTCACGCTGTAGGTATCTCAGTTCGGTGTAG
GTCGTTCGCTCCAAGCTGGGCTGTGTGCACGAACCCCCCGTTCAGCCCGACCGCTGCGCCTTATCCGGTAACTAT
CGTCTTGAGTCCAACCCGGTAAGACACGACTTATCGCCACTGGCAGCAGCCACTGGTAACAGGATTAGCAGAGCG
AGGTATGTAGGCGGTGCTACAGAGTTCTTGAAGTGGTGGCCTAACTACGGCTACACTAGAAGAACAGTATTTGGT
ATCTGCGCTCTGCTGAAGCCAGTTACCTTCGGAAAAAGAGTTGGTAGCTCTTGATCCGGCAAACAAACCACCGCT
GGTAGCGGTGGTTTTTTTGTTTGCAAGCAGCAGATTACGCGCAGAAAAAAAGGATCTCAAGAAGATCCTTTGATC
TTTTCTACGGGGTCTGACGCTCAGTGGAACGAAAACTCACGTTAAGGGATTTTGGTCATGAGATTATCAAAAAGG
ATCTTCACCTAGATCCTTTTGGTTCATGTGCAGCTCCATCAGCAAAAGGGGATGATAAGTTTATCACCACCGACT
ATTTGCAACAGTGCCGTTGATCGTGCTATGATCGACTGATGTCATCAGCGGTGGAGTGCAATGTCGTGCAATACG
AATGGCGAAAAGCCGAGCTCATCGGTCAGCTTCTCAACCTTGGGGTTACCCCCGGCGGTGTGCTGCTGGTCCACA
GCTCCTTCCGTAGCGTCCGGCCCCTCGAAGATGGGCCACTTGGACTGATCGAGGCCCTGCGTGCTGCGCTGGGTC
CGGGAGGGACGCTCGTCATGCCCTCGTGGTCAGGTCTGGACGACGAGCCGTTCGATCCTGCCACGTCGCCCGTTA
CACCGGACCTTGGAGTTGTCTCTGACACATTCTGGCGCCTGCCAAATGTAAAGCGCAGCGCCCATCCATTTGCCT
TTGCGGCAGCGGGGCCACAGGCAGAGCAGATCATCTCTGATCCATTGCCCCTGCCACCTCACTCGCCTGCAAGCC
CGGTCGCCCGTGTCCATGAACTCGATGGGCAGGTACTTCTCCTCGGCGTGGGACACGATGCCAACACGACGCTGC
ATCTTGCCGAGTTGATGGCAAAGGTTCCCTATGGGGTGCCGAGACACTGCACCATTCTTCAGGATGGCAAGTTGG
TACGCGTCGATTATCTCGAGAATGACCACTGCTGTGAGCGCTTTGCCTTGGCGGACAGGTGGCTCAAGGAGAAGA
GCCTTCAGAAGGAAGGTCCAGTCGGTCATGCCTTTGCTCGGTTGATCCGCTCCCGCGACATTGTGGCGACAGCCC
TGGGTCAACTGGGCCGAGATCCGTTGATCTTCCTGCATCCGCCAGAGGCGGGATGCGAAGAATGCGATGCCGCTC
GCCAGTCGATTGGCTGAGCTCATGAGCGGAGAACGAGATGACGTTGGAGGGGCAAGGTCGCGCTGATTGCTGGGG
CAACACGTGGAGCGGATCGGGGATTGTCTTTCTTCAGCTCGCTGATGATATGCTGACGCTCAATGCCGTTTGGCC
TCCGACTAACGAAAATCCCGCATTTGGACGGCTGATCCGATTGGCACGGCGGACGGCGAATGGCGGAGCAGACGC
TCGTCCGGGGGCAATGAGATATGAAAAAGCCTGAACTCACCGCGACGTATCGGGCCCTGGCCAGCTAGCTAGAGT
CGACCTGCAGGTCCCCGGGGATCGGTCTTGCCTTGCTCGTCGGTGATGTACTTCACCAGCTCCGCGAAGTCGCTC
TTCTTGATGGAGCGCATGGGGACGTGCTTGGCAATCACGCGCACCCCCCGGCCGTTTTAGCGGCTAAAAAAGTCA
TGGCTCTGCCCTCGGGCGGACCACGCCCATCATGACCTTGCCAAGCTCGTCCTGCTTCTCTTCGATCTTCGCCAG

CAGGGCGAGGATCGTGGCATCACCGAACCGCGCCGTGCGCGGGTCGTCGGTGAGCCAGAGTTTCAGCAGGCCGCC
CAGGCGGCCCAGGTCGCCATTGATGCGGGCCAGCTCGCGGACGTGCTCATAGTCCACGACGCCCGTGATTTTGTA
GCCCTGGCCGACGGCCAGCAGGTAGGCCGACAGGCTCATGCCGGCCGCCGCCGCCTTTTCCTCAATCGCTCTTCG
TTCGTCTGGAAGGCAGTACACCTTGATAGGTGGG

#### 7.2 pUWL201PW::*ika*

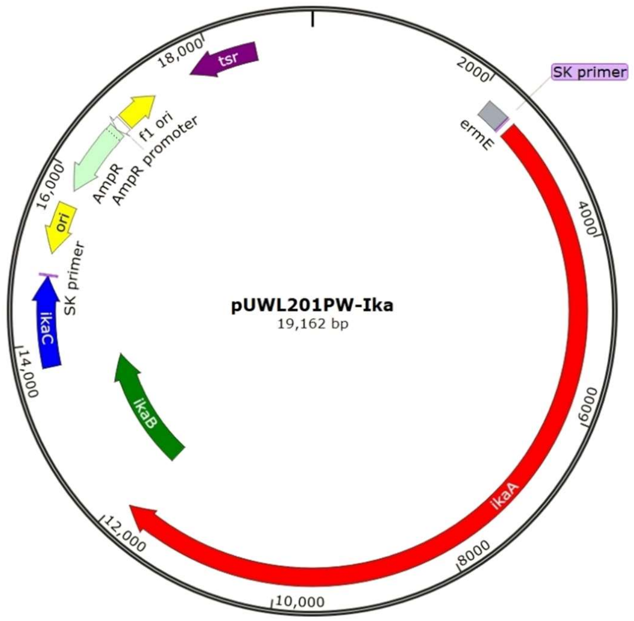

**Figure S3.** Vector map of *ika* in pUWL201PW.

- DNA sequence of pUWL201PW::*ika* [19162 bp]:

```
CGTCAAAGCCCCGCCGGATCACCGGCGGGGCTCTCTTCGGCCCTCCAAGTCACACCAGCCCCAAGGGGCGTCGGG
AGTGGCGGAGGGAACCTCTGGCCCGATTGGTGCCAGGATTCCCACCAGACCAAAGAGCAACGGGCCGGACTTCGC
ACCTCCGACCCGTCCGCTCCCAGACTCGCGCCCCTTAGCCGGGCGAGACAGGAACGTTGCTCGTGCCCAGAGTAC
GGAGCGATGCCGAGGCATTGCCAGATCGGCCCGCCGGGCCCCGCTGCCACTGCGGGACCGCAATTGCCCACACAC
CGGGCAAACGGCCGCGTATCTACTGCTCAGACCGCTGCCGGATGGCAGCGAAGCGGGCGATCGCGCGTGTGACGC
GAGATGCCGCCCGAGGCAAAAGCGAACACCTTGGGAAAGAAACAACAGAGTTTCCCGCACCCCTCCGACCTGCGG
TTTCTCCGGACGGGGTGGATGGGGAGAGCCCGAGAGGCGACAGCCTCTCGGAAGTAGGAAGCACGTCGCGGAGCG
ACGCTGCCCGACTGCGGAAAGCCGCCCGGTACAGCCGCCGCCGGACGCTGTGGCGGATCAGCGGGGACGCCGCGT
GCAAGGGCTGCGGCCGCGCCCTGATGGACCCTGCCTCCGGCGTAATCGTCGCCCAGACGGCGGCCGGAACGTCCG
TGGTCCTGGGCCTGATGCGGTGCGGGCGGATCTGGCTCTGCCCGGTCTGCGCCGCCACGATCCGGCACAAGCGGG
CCGAGGAGATCACCGCCGCCGTGGTCGAGTGGATCAAGCGCGGGGGGACCGCCTACCTGGTCACCTTCACGGCCC
GCCATGGGCACACGGACCGGCTCGCGGACCTCATGGACGCCCTCCAGGGCACCCGGAAGACGCCGGACAGCCCCC
GGCGGCCGGGCGCCTACCAGCGACTGATCACGGGCGGCACGTGGGCCGGACGCCGGGCCAAGGACGGGCACCGGG
CCGCCGACCGCGAGGGCATCCGAGACCGGATCGGGTACGTCGGCATGATCCGCGCGACCGAAGTCACCGTGGGGC
AGATCAACGGCTGGCACCCGCACATCCACGCGATCGTCCTGGTCGGCGGCCGGACCGAGGGGGAGCGGTCCGCGA
AGCAGATCGTCGCCACCTTCGAGCCGACCGGCGCCGCGCTCGACGAGTGGCAGGGGCACTGGCGGTCCGTGTGGA
CCGCCGCCCTGCGCAAGGTCAACCCCGCCTTCACGCCCGACGACCGGCACGGCGTCGACTTCAAGCGGCTGGAGA
CCGAGCGCGACGCCAACGACCTCGCCGAGTACATCGCCAAGACCCAGGACGGGAAGGCGCCCGCCCTCGAACTCG
CCCGCGCCGACCTCAAGACGGCGACCGGCGGGAACGTCGCCCCGTTCGAACTCCTCGGACGGATCGGGGACCTGA
CCGGCGGCATGACCGAGGACGACGCCGCCGGGGTCGGCTCGCTGGAGTGGAACCTCTCGCGCTGGCACGAGTACG
AGCGGGCAACCCGGGGACGCCGGGCCATCGAATGGACCCGCTACCTGCGGCAGATGCTCGGGCTCGACGGCGGCG
ACACCGAGGCCGACGACCTCGATCTGCTCCTGGCGGCCGACGCCGACGGCGGGGAGCTGCGGGCCGGGGTCGCCG
TGACCGAGGACGGATGGCACGCGGTCACCCGCCGCGCCCTCGACCTNCGAGGCGACCCGGGCCGCCGAAGGCAAG
GACGGCAACGAGGATTCGGCGGCCGTGGGCGAACGGGTGCGGGAGGTCCTGGCGCTGGCCGACGCGGCCGACACA
GTGGTGGTGCTCACGGCGGGGGAGGTGGCCGAGGCGTACGCCGACATGCTCGCCGCCCTCGCCCAGCGCCGCGAG
GAAGCAACTGCACGCCGACGGCGAGAGCAGGACGACGACCAGGACGACGACGCCGACGACCGCCAGGAGCGGGCC
GCCCGGCACATCGCCCGGCTCGCAAGTGGGCCCACTTCGCACTAACTCGCTCCCCCCCGCCGTACGTCATCCCGG
TGACGTACGGCGGGGGTCGGTGACGTACGCGGCGACGGCGGCCGGGGTCGAAGCCGCGGGAGTAATCCTGGGATT
```

ACTCGCCCGGGGTCGGCCCCGCCGGCACTTCGTGCAGGCGGTACCAGCCCGACCCGAGCACGCGCCGGCACGCCT
GGTCGATGTCGGACCGGAGTTCGAGGTACGCGGCTTGCAGGTCCAGGAAGGGGACGTCCATGCGAGTGTCCGTTC
GAGTGGCGGCTTGCGCCCGATGCTAGTCGCGGTTGATCGGCGATCGCAGGTGCACGCGGTCGATCTTGACGGCTG
GCGAGAGGTGCGGGGAGGATCTGACCGACGCGGTCCACACGTGGCACCGCGATGCTGTTGTGGGCACAATCGTGC
CGGTTGGTAGGATCGATCCACTAGTTCTAGAAATAATTTTGTTTAACATTAAAGAGGAGAAATTACATATGGCTA
GCATGACGGTATCGATAAGCTTGATATCGAATTCATGGATTCCATGCACCACCCTGCCCCCGTCCCCGTACCCGA
AGTCCCCGCGCCCGTCCCGTCCCAGGACGACGCGTTCGCCATCGTCGGCATCGGCTGCCGGCTGCCCGGCGGCGC
CAGCGACTACCGGACCTTCTGGCGCAACCTCCTCGACGGCAAGGACTGCATCACCGACACCCCCGCCGACCGCTA
CGACACCCGCACCCTGGGCAGCGGCGACAAGGCCAAGCCCGGCCGGCTGGTCGGCGGACGCGGTGGATACATCGA
CGGCTTCGACGAGTTCGACCCCGCCTTCTTCGGCATCAGCCCGCGCGAGGCCGAGCACATGGACCCCCAGCAGCG
GAAGCTCCTGGAGGTCGCCTGGGAGGCGCTGGAGGACGGCGGCCTCAAGCCCGCCGAGCTGGCCGGCAGCGATGT
CGGGGTGTACGTCGGGGCGTTCACCCTCGACTACAAGATCCTGCAGTTCGCCGACCTCGGCTTCGAGACCCTGGC
CGCGCACACCGCCACCGGCACCATGATGACGATGGTGTCCAACCGGATCTCGTACTGCTTCGACTTCCGCGGACC
CTCGGTCTCCGTCGACACCGCGTGCAGCGGCTCCCTGGTCGCCGTCCACCTCGCCTGCCAGAGCCTGCGCCGCGG
CGAGACCTCCGTCGCCCTGGCCGGCGGCACCCTGCTGCACATGGCGCCGCAGTACACCATCGCCGAGACCAAGGG
CGGGTTCCTCTCCCCCGACGGCCGCTCCCGCGCCCTGGACGCCTCCGCCAACGGCTACGTGCGCGCCGAGGGCGT
CGGCATGGTCGCCATCAAGCGCCTCGCGGACGCGCAGCGCGACGGCGATCCCATCCACGCCGTCATCATCGGCAG
CGGCGTCAACCAGGACGGCCGCACCAACGGCATCACCGTGCCCAACCCCGACGCGCAGGTCGCCCTGATCGAGCG
GGTCTGCGCCGCCGCCGGCGTCACCCCCGGCAGCCTCCAGTACGTCGAGGCGCACGGCACCTCCACCCCCGTCGG
CGACCCGCTGGAGGCCAACGCCCTCGGCCGCGCGCTCTCCATCGGCCGCGAGCCGGGCGCCCGGACGTACGTCGG
CTCGGTCAAGACCAACATCGGGCACACCGAGTCCGCCGCCGGCATCGCCGGGCTGATCAAGACGGTGCTCAGCCT
CAAGCACAAGGTCATCCCGCCGCACATCAACCTGGAGAAGCTCAACCCGCAGATCGACGAGGCGTCCCTGCCGTA
CGAGATCCCCCGCGAGCCCACCCCCTGGCCCGAGCACAGCGGGCCGGCCCGGGCCGGCGTCAACTCCTTCGGCTT
CGGCGGGACCAACGCCCACGTCCTGCTCCAGGAGGCACCGCCGACCGTCGGGGAGCCCGCGCCACCGGCCACCGA
CGGGTACTCCGTGCTGCCGCTCAGCGCCCGCGACCCCGAAGCCTTTCCCGCCATCGCCACCGGCCTGCGCGAACG
GCTCGCCGAGGGACTGCCGGTGGGCGACGCCGCCTACACCCTCGCCCACCGGCGGCAGCATCTGGAGCAGCGGCT
GTCCGTCGTGTACGACTCCCCCGAGGCCCTCGACGAGGTGCTCGGCGCCGTCGCCCGCGGCGAGAGCCACCCGCG
TGCCGTCGCCGGCACCCAGCGGGAGGGCCTGGACCGCAGGCTGGTGTGGGTGTTCACCGGCATGGGCCCGCAGTG
GTGGGCCATGGGCCGCCAGTTGTACGCGAGCGAGCCCGTCTACCGGGAGGTCATCGACCGCTGCGACCAGGAGAT
CGCCGCGCTCACCGGCTGGTCCCTCACCCAGGAGCTGAACGCCGACGAGGCCGACTCCCGGATGAGCGAGACCTG
GCTCGCCCAGCCCGCCAACTTCGCCGTCCAGATCGCCCTGGCCGCCCTGTGGCGCAGCAAGGGGATCCAGCCCGA
CGCCGTCACCGGGCACAGCACCGGTGAGGTCGCCGCGTTCTACGAGGCCGGGGTGTACACCCTCCCCGAGGCCGT
GAAGATCGTGGTGCACCGCAGCCGGCTCCAGCAGAAGCTCATCGGCACCGGCTCCATGCTCGCCGTCAGCCTCAC
CGAGGCGGAGGCCGCCCGCCGGGTGCGCCCGCACGGCGACCGGGTCTCCATCGCCGCCGTCAACAGCCCCACCTC
CATCACCCTGGCCGGGGACACCGAGGCGCTGGAGGTGATCGCCGCCGAGCTGGGCGCCGAGGACATCTTCGCCCG
CTTCCTGGAGGTCGGCGTCCCGTACCACAGCCCCCGCATGGAGCTGATCAAGGACGAGCTGCTGACCTCGCTCGC
CGATCTCAAGCCGCAGCAGGCGAAGTTGCCGCTGTACCTCACCGCGCTGCCGGGCACCGTCGCGCAGGGCACGGA
GCTGGACGCCGACTACTGGTGGCGCAATGTGCGCGAGGCCGTGCACTTCCGGGCCGCCGTGGACCGGCTGCTGGA
CGACGGCTACGGCGTCTTCCTGGAGATCGGCCCGCACCCCGTGCTCGCCCACTCCCTGCGCGAGTGCTGCGAGGC
CCGCGACGCGCACAGCGTCACCCTGGCCTCCATCCGCCGCAAGGCGGACGAGCGCGAACGCCTCACCCTGTCGCT
CGCCGCGCTGCACAGCCTCGGCTTCGCCGTGGACTGGCACGCCCTGCACCCCGCCGGGCGGCCGGCCGAACTGCC
GCGCTACCCGTTCCGGCGCGACCGGTACTGGGTCGAGCCGGCCCCGGTCGCGCAGATCCGGCTCGGCCACCGCGA
CCACCCGCTGCTGGGCCGCCGCACCGCGAGCGCCGAGCCGGTGTGGGAGGTGAAGCTGGACGCGGAGGCCGCCCC
GTACCTGGAGGACCACCGCATCCAGGGCACCGTGCTGTTCCCGGCCGCCGGCTATCTGGAGATGGCCGCGCAGGC
CATGCGGGCGCTGACCGGTGATGAGCACAGCACCGCCGCGCTGGCCGGCATCGAGCTGCGCAAGGCGCTGTTCCT
GCCGGACGGCGAGCCGCAGACGGTGCAGCTGTCCTTCTCCTCCGACGCCGCCGCGTTCTCCATCGCCACCGTGGG
CGCCGCCGGCGCCGAGCCGACCGTGCACGCCACCGGTACGGTACGGGCCGCCCAGCGCCGCCGGCTGACCGCGCC
GCTGGACACCGTCGCCGTCCGGGCCCGCGCCGCCCGCCACCTGAGCGGCCCCGACTGCTACGCCGAACTGGCCGC
GCTCGGCTACCACTACGGCCCCGCCTTCCAGGGCATCGAGGAGGTGTGGATCGGCGAGGGCGAGGCCCTGGCCCG
GATCCGTCCGCCGCAGGGGCTCACCCCGGACGCGGCGGCGCACCACATGCATCCGGTGCTGCTCGACTCCTGCTT
CCAGTCGCTGCTGACCCCGCAGCTGCTCACCGCGCCCGCCGGGCCCGGGGGCACCGGCATCCGGCTGCCGCTGTC
CATCGCCGAGGTACGGCTGGACCCGGTCGGCGACCGCGAACTGTGGGTGCACGCCACCGTCACCGGCGACGACGA
GGACGAACTCACCGGTGACATCGCCGTGTACGACGGCGCCGACGGTACGCCGCTGGGCCGCGTCGCCGGCTTCCG
CGCCGCCGATGTGGAGAAGGCCGCCACCACCGTGGGGCTGTCCACCATCGACAGCTGGCTCACCGAACCGAGCTG

GGTGCCGTGCCCGCTGCCCGAGGCGGCGTCCGCCGCGCCGGCGGCCGGGCGGCACGTACTGTTCGCCGACGCGGG
CGGGGTCGCGCAGCGGCTGGCCGCGCTGATCGGCGAGGCCGGCGGGGAGGCCCATCTGGTCCGGCCCGGTGCCGC
GTACGGCCTGGACCGCACGGCGAGGACCGCCACCGTCGTCCCCGGATCCGCGGATGACCTGCGGCGGTTGCTCAC
CGATCTCGGGCAGGTGGACGGCGTCGTCCACCTGTGGAACCTGGACCGGCCGGCGCTGGCCGACGCCCCGCGCGG
ACGGTTCGCGGACATCGCCTCCACCGGCGCGTACGCCCTGATCGCCCTCACTCAGGCCCTGCTCGCCGACCCGGA
GCGGCACGGCGGCACCCCGGTGCACATCGTCACCAGAGCCGCCCAGTGCGTGGTCCCCGGTGAGCCGGTGGAGCC
GCTGGGCGCGCCCGCCTGGGGCATCGGCCGGGTGCTGTGGCAGCAGGAACTGGCCGGGCGCGGCGGCAAGCTGAT
CGACCTGGCGGCCGACGGCGGCGTCGAGGAGGACGCGTACGCGCTGCTGCGCGAGCTGGCCGACCCCACCGGCGC
GGCCGAGCGCGAGGACGAGATCGCGCTGCGCGCCGGGGAGCGGCACACCAGCCGGCTGGTGGCCGCCGAGGGGCT
GAGCAGGCCGCTGCCCCTGCGGCTGCGCCCGGACGGCAGCTATCTGGTGACCGGCGCGTTCGGCGCGCTCGGCAG
GCTGCTGTGCCGCACGCTGGTCAGGCGCGGGGCGCGGCGGCTGATCCTGGTGGGCCGCACCCGGCTGCCGGAGCG
CGAGCGCTGGGCCGACCAGGACCCGAACTCGCCGGCCGGGCGGCACGTGGCCTTCCTCAAGGAGCTGGAGGCGCT
GGGCGCGCAGCCGATTCTCGCGCCGCTGGACATCACCGACGAGGACGCGCTGGCCGGCTGGCTCGCCGGGTACCG
GCGCGCCCAGGGGCCGCCGATCCGCGGGGTGTTCCATCTGGCGGGGCAGGTGCGCGACACCCTGGTGCCGGAGAT
GGACCGGGAGGTGTTCGACGCCGTCCACGACCCGAAGGTGGTGGGCGCGGCGCTGCTGCACCGGCAGCTGAGCGG
CGAACCGCTGGAGCACTTCGTGCTGTTCGCCTCGGTCGCGGCCTGGCTGACGACGGCCGGACAGACCAACTACGC
GGCGGGGAACGCCTTCCTGGACGCGCTGGCGCACCACCGCCGCGCGCAGGGGCTGCCGGCGCTGGCGCTGGACTG
GGGCCCGTGGGCCACCGGCATGATCGAGGAACTGGGCCTGATCGACCACTACCGCAACAGCCGGGGCATGTCCTC
GCTGGCGCCCGAGGCGGGCATGGCGGTGCTGGAGCGGGTCATCGGGCAGGACCGGGCACAGCTGCTGGTGGCCAC
GGTCGTGGACTGGCCGGTGTTCATGTCCTGGTACGCGGCGCCGCCGCGGCTGGTCACGGAGCTGGCGGCCACCGC
CCAGGGACCGGGGTCCGAGGGCGACGGCAGTTTCCTGGACGCGTTCCGGGAGGCCACCGCGGACAAGCGGCGGCT
GCTGCTGACCGAGCGGTTCACGACGCTGGTGGCGGGTGTGCTGCGGGTGCGGGCCGAGCAGGTGGATCCGGCGGT
CAGCCTGAATCTGCTGGGGCTCGACTCGCTGCTGGCGATGGAGCTGCGAGCGCGGGTGGTGGCCGAGGTGGGCAT
CGCGCTGCCGGTGGTGGCGCTGCTGTCCAGCGCGCCGGCCGGGGACCTGATCACCCAGCTGCACGAGGGCCTGGA
GGAGTTGCTGGCCGAGGAGGGCAGCGGCGCCGCGGTGACGGCGGTGGAGCGCTTCGAGGACGAGGCCGAGTTCCC
GCTGACGCAGAACCAGAAGGCGCTGTGGTTCCTGAAGCAGCTGAACCCGGACGGCTTCGCGTACAACATCGGCGG
CGCCGTCGAGGTGCGGGTCGAGCTGGACCCGGACCTGATGTTCGAGGCGTTTCGCCGGCTGCTGGCCCGGCATCC
CGTGCTGCGGGCGAACTTCCTGCTGGTGGAGGGGCAGGCGGTGCAGCGGATCTCCCCGGAGATCAAGGAGGACAT
CGCGCTCTTCGACGTCGAGGACCGCGCGTGGGACGACATCTACCGGATGATCATCGAGGAGTACCGCAAGCCGTA
CGACCTGGCGACCGATCCGCTGATCCGGTTCCGCCTCTTCCGGCGCGGCCCGGACCGCTGGGTCATCACCAAGGC
CGTCCACCACATCATCTCGGACGCCATCTCCACCTTCACCTTCATCGAGGAACTGCTGTCCCTGTACGAGGGGCT
GCGGCAGGGCCACGACGTCGAACTGCCGCCGGTGTCCGCCCGCTATCTGGACTTCCTCAACTGGCAGAACGCGTT
CCTGGCCGGCCGCGAGGCGCAGAAGATGCTCGCGTACTGGCGGGGGCAGCTGCCGGACGAGGTGCCGGTGCTGGC
GCTGCCCACCGACAAGCCGCGCCCGGCGGTGCTCACCCACAACGGGGCGTCCGAGTTCTTCGCCCTGGACGCGGA
GTTGAGCGCCCGGGTGCACGCGCTGGCGCGGGAGCACAACGTCACCGTCTTCATGGTGCTGCTGAGCGCGTACTA
CCTGCTGCTGCACCGCTATGCGGGGCAGGACGACATCATCGTCGGCTCCCCCGTCACCGGCCGCACCCAGGAGGA
GTTCGGCGCCGTCTACGGGTACTTCGTGAACCCGCTGCCGCTGCACGCCTCGCTGGCCGGTGACCCCACGGTCGC
CGAGCTGCTGGACCAGGTGCGCACCACGGTGCTGGGCGGCCTGGACCACCAGGAGTACCCGTTCACGCTGCTGGT
GGAGCAGCTGGGGCTGGCCCACGACCCGAGCCGGTCGGCGGTCTTCCAGGCGATGTTCATCCTGCTGCACCACAA
GGTGGCCACCGAGAAGTACGGCTACAAGCTGGAGTACATCGAGCTGCCCGAGGAGGAGGGCCAGTTCGACCTGAC
GCTGTCCGCGTACGAGGAGGAGGCGGACGGGCGGTTCCACTGCGTCTTCAAGTACAACACCGACCTCTTCGAGGC
GGAGACGATCCGGCGGCTCGCCGGGCACTACACGCAGCTCCTGGAGTCGCTGACCGCGGCGCCCGCCGACGCCGC
CACCGGTGGACTGCGGATGCTGTCGGGCGGCGAGCGGGAGCGGATCCTCACCGAGTGGAGCGGGGCCGGGCAGGG
CGCGCAGGACGCGCCGGTGCCGGTGCACCGGCTGATCGCCGAGGCGGCGCACCGTACCCCGCAGGCGATCGCGGT
GGCCGCGCCCGCCGAGAGCGGGGAGACCCGGCGGCTGACGTACGGCGAACTGGAGGAGCGCGCCGGCGAACTGGC
CGGGCGGCTGCGGGCGCGCGGCGTGCGCGAGGGCACCGTCGTCGCGCTGTGCCTGGAGAAGTCGCCCGAGCTGAT
CACCGCCCTGCTGGCGGTCCTCAAGGCGGGCGGCGCCTATCTGCCGCTGGACCCGGACTATCCGGCCGACCGGCT
CGCGTACATGGTGCGCAACGCCGGGGCCACGCTGGTGATCGGCGGGACGGGCGGCGCGGCCGAGGGGCTGCCGGG
CACCGTGGTCACCCTGGAGGAACTGCTCGCGGGCGAGGCCGGCGAAGCGGGGCCGGACGCCGAGCCGGGGCCCGA
CTCCCCCGCCTACGTCATCTACACCTCGGGCTCCACCGGGCGCCCCAAGGCGGTCGCGGTCAGCCACCGCAATCT
GGCCTCGGTGTACGCCGGATGGCGCGACGCCTACCGCCTGGAGGAGGGCGGCATCCGGGTCCATCTCCAGATGGC
CAGCCCCTCCTTCGACGTCTTCACCGGCGACCTGACCCGAGCCCTGTGCTCGGGCGGCACGCTGGTGCTGGTCGG
CCGGGAGCTGCTGTTCAACACCGCCCGGCTGTACGAGACGATGCGCGCCGAACGGGTGGACTGCGGCGAGTTCGT
GCCCGCCGTGGTGCGCACCCTGGTGCGGCACTGCGAGGACACCGGCGCCCGGCTGGACTTCCTGCGGCTGCTGAT

CGTGGGCTCGGACTCCTGGAAGGCCGAGGAGTACGAGCGGCTGCGCGCGCTGGGCGCACAGCGCCTGGTGAACTC
GTACGGGCTCACCGAGGCCACCATCGACAGCGCCTGGTTCGAGGGTCCCGCGGATGACCTGGAGGGCGGCCGGAT
GGTGCCCATCGGGCGGCCGTTCCCGGGCAGCGCGCTGTACATCCTGGACTCGCGCGGCGAGCCGGTGCCGCCCGG
TGTCCCCGGCGAGCTGTGGATCGGCGGCACCGGGGTGGCGCTCGGCTACCTCGGCGACGAGGCGCTGACCGGGGA
GCGGTTCCTCACCCGCGCCCTGGCCGGCGACGCTCCGGTACGGCTGTACCGCACCGGTGACCTCGCGCGCTGGGA
CGCGGCCGGCACCGTCCATCTGCTGGGCCGGGCCGACTCGCAGATCAAGGTGCGCGGGCACCGCATCGAGATCGG
GGAGATCGAGTCGCACCTGGCGGCCTGCCCCGAGCTGGCCCAGGCGCAGGTCACCGTGCGGCCGGACGCGGGCGG
CGAGAACGTGCTGTGCGCGTACGGGGTGGCGGCCCCGGGCGCCGTGCTGGACTGGCGCGAGGTGCGCCGGCGCCT
GGCGGACTATCTGCCGACGTTCATGATCCCCACCCACTTCACCGAGCTGCCCGCCCTGCCGCTCACCCCGAACGG
CAAGGTGGACGTGGCGGCGCTGCCCGCCCCGCGCACCGGCGACGGCGCGGACGGGCCGGTGTACGAGGCCCCCGT
CACGCTGTACGAGACCCGGATGGCCGAGCACTGGCAGCGGCTGCTGGGCATCGAGGCCCCCGGGCCCGGTCTGGG
CCACGACTTCTTCGAGACCGGTGGCAGCTCCATCCGGCTGATCGAGCTGATCTACCACCTGCAGGCCGAGTTCGG
GATCTCCATCCCGGTCAGCCGGCTGTTCCAGGTGACGACGCTGCACGGCATGGCCAAGACGGTCGAGCGGATCGT
CACCGGGGAGATCGAGGGGTCGCTGCCGTATCTGCGGTTCAACGAGAACGCCGCGGCGGGCACGGTGTTCTGCTT
CCCGCCGGCCGGTGGCCACGGCCTGGTCTACCGGGAGTTCGCGGCGCGGCTGCCGGAGTTCGAGTTCCTCGCCTT
CAACTACCTGATGGGCGAGGACAAGGTAAGCGGGTACGCCGACCTGGTGGCCGGGCACCGGCCGGAGGGCGAGAT
CGACCTGCTCGGCTACTCGCTGGGCGGCAACCTCGCCTTCGAGGTGGCCAAGGAGCTGGAGCGGCGCGGCCGCAC
CGTGCGCCACGTCGTCATCATGGACTCGCTGCGGGTGACGGAGTCCTACGAGCTGGGCCCGGAGCACCTGGCCGT
CTTCGAGCGCGAGCTGGCCGAGCATCTGCGCAAGCACACCGGCTCGGCGCTGGTCGCGGAGAAGACGCGCGAACA
GGCCAAGGACTACCTGGAGTTCACCGGCCGCACCGCCAACCCCGGCACCACCGGGGCCCGGATCGCGGTGATCAG
TGACGAGGAGAACGCGGCCGCGTACGACAGCGGCGCCGAGGGCAGCTGGCACGGCGCCTCCCGTACCGGAACCGA
CGTGCTGCGCGGGGTGGGCCGGCACGCCGACATGCTCGATCCGGGGACGGTCGAGCACAACGCGCGCCTGGCGCG
CGGCATTCTCACCGGCGGTGATGGCGAGGTATGACGCCTTTCGTTCAGCCGGCGGTCGACACCAAGGAGCACAGC
GCCATGTCATCCCCCACCACCTCCGGCACCCCGGGCAGGCAGTCGATGATCATCATCGGCGGCGGCCTGGGGGGC
CTGTCCACCGGCTGCTACGCGCAGATGAACGGCTACGCGACGCGGGTCTTCGAGATGCACGAGATCCCGGGCGGT
TCCTGCACCGCCTGGGAGCGCGGGGACTTCACCTTCGACTGGTGCGTCAGCTGGCTGCTGGGCAGCGGTCCCGGC
AACGAGATGTACCAGATCTGGATGGAACTGGGGGCGTTGCAGGGCAAGGAGATGCGCCAGTTCGACGTCTTCAAC
ATCGTGCGGGTGCGCGGCGGCCAGCCGGTGTACTTCTACTCCGACCCGGACCGGCTCCAGGCGCACCTGCTGGAG
ATCTCCCCGGCCGACGCCCGCCGCATCAAGAACTTCTGCGAGGGGGTGCGCACCTTCCAGAAGGCGCTGTCGGTC
TACCCGTTCCTCAAGCCGGTGGGGCTGATGGGGCGGTGGGAACGGTGGAAGATGCTGGCCTCGTTCCTGCCGTAC
TTCAACGCCATCCGCAAGTCCATCACCGAGCTGATGACGGACTACGCGGAGAAGTTCCAGCACCCGGTGCTGCGC
GAGGCCTTCAACTACGTGCTGTACGAGAAGCACGCCGACTTCCCCGTCCTGCCGTTCTGGTTCCAGCTGGCCTCG
CACGCCAACGGCTCGGCGGGGGTGCCCGAGGGCGGCTCGCTGGAGCTGGCCCGGTCCGTGGAGCGGCGCTACCTG
GGGCTCGGCGGGGAGATCACCTACAACGCCAAGGTGGAGAAGATCCTCGTCGAGCACGACAAGGCGGTGGGAGTG
CGGCTCACCGACGGCCGCGAGTTCCGCGCGGACATCGTGGTGTCGGCGGCCGATCTGCACACCACCGCCATGGAG
ATGCTCGGCGGCCGGTATCTCAACGACACCTGGCGCAAGCTGCTCACCGAGACGATCGACGAGGTGGGCACGATC
TCCCCCGGCTATGTCTCGCTGTTCCTGGGGCTGCGCCGGCCGTTCCCCGAGGGCGAGCCGTGCACCACGTACGTG
CTGGAGGACAGCATGGCGGAGAAGCTCACCGGCATGCGGCATCCCAGCATGAACGTGCAGTTCCGCAGCTGCCAC
TACCCGGAGCTGTCGCCGCGCGAGACCACGGTCATCTTCGCCACGTACTTCTCGGAGGCCGAGCCGTGGCGGGCG
CTGCGCGACGACGTGCCGGAACAGGCGGGCCGGGTGCGGCGCGGTCAGGTGCTGCACACCCTGCCGGTGAAGCAC
GGCAAGGCGTACACCCAGGCCAAGCGGCAGGCGCGGATCACCATCGAGAACTTCCTGGACGAGCGGTTCCCCGGT
CTCAAGGACGCGGTCGCCGTGCGGGACGTGTCCACGCCGCTGACGCAGGTGCGCTACACGGGCACCTACAACGGC
GGGTTCCCCGGCTGGCAGCCGTTCGTGGACGGCGGGGAGACCGTGGAGGTGGAGATCAACAAGAACGGCCCGGTG
CTGCCGGGGCTCTCCAACTTCTATCTGGCCGGGGTGTGGGTCACCGTCGGCGGGCTGATCCGGGCGGTGGCCTCG
GGCCGGCAGGTCACGCAGGTGATCTGCCGGGACGACGGGCGGGAGTTCACGGCGAGCGTGGACGAGAGCGCGCCG
CCGCCCACCCAGGTCGCCATCCCGGTGGGCAAGCAGCCGGGCGTGCCGGATCTGGCGGCCGGGTTCCCCGCCCAG
ACCGCCGGCGGCGAGACCGCCACCGGCGCCAACAACACCGTCACATCGTCGAGGAGCGTGTAGTGCAGGTACACA
CATGGGTGATCGCCGAGCACATCCCGGGGGTCCCGGACACGGACCGGATCTACCGGAAGGTGACGCGGGAGTTCG
ATCCCGCCTCGCTGGCGGACGATCAGATGCTGCTGCGCACCCGGTACGTGTCGGTGGACCCGTATCTGGTGGGGC
TCTCGCTCCAGACGCCGATCGGGGACACCGTGCGCGGTGACTCGATCATGGAGGTGGCCGTGGCGGGGCCGCGCG
CCCGCTTCCAGGTCGGGGACCTGGTGCAGGGGTACGGCGGCTGGTGCAGCCATCTGGTGTCCACCGGGGGGCCCA
GCGGATGGAACGACGACGGCGCCGAGTTCGCCGTCCAGTTGCCGCCGTTCCGCAAGCTGGACCCGCGGCGGTACG
ACGAGGCGCTGCCGCTGTCCACGGCGCTGGGCGTGATGGGCACCCCGGGCATCACCGCGTTCGGCGCGATGAAGA
CGTTCCTGACCGTGGGCTCCGAGGACACGGTGGTGATCAGCGGGGCGTCCGGGACGGTGGGCACCCTGGTGGGCC

AGCTCGCCAAGCGGGCCGGGGCCCGGGTGGTGGGCACCACCTCCTCGCCGGGGAAGGCCGCGTATCTGACGCAGC
TGGGCTTCGACGCGGTGGTGAACTACCGGCAGGGCGACGACACGGACACGGTGCGCGAGGCGCTGGCGGCGGCAG
CGCCCAACGGAATCGACAAGTACTTCGACAACCTGGGCGGCACCGTGACGGACGCGGTGTTCACGATGCTCAACG
TGCACTCCCAGGTGGCGGTGTGCTGGCAGTGGGCCACCACGGTCAACGGGGACTGGACGGGGCCGCGGCTGCTGC
CGTACATCATGTTCCCGCGCACCACGATCCGGGGGATCTTCGCCGACGAGTGGTACACGGAGGAGATGGTCGACG
CGCTGCACGAGGAGGTGGGCGGGCTGATCCGCAAGGGTGAGCTGGCCTACCACCAGACCATCCACCAGGGCTTCG
ACGCCCTCCCGGACGCGTACCGCTCCCTGTACACCGGCCAGGAGGGCAACCGCGGCAAGGTCCTGGTCGCCCTGT
AGGATCCACTAGTTCTAGAAGCTCTGCATTAATGAATCGGCCAACGCGCGGGGAGAGGCGGTTTGCGTATTGGGC
GCTCTTCCGCTTCCTCGCTCACTGACTCGCTGCGCTCGGTCGTTCGGCTGCGGCGAGCGGTATCAGCTCACTCAA
AGGCGGTAATACGGTTATCCACAGAATCAGGGGATAACGCAGGAAAGAACATGTGAGCAAAAGGCCAGCAAAAGG
CCAGGAACCGTAAAAAGGCCGCGTTGCTGGCGTTTTTCCATAGGCTCCGCCCCCCTGACGAGCATCACAAAAATC
GACGCTCAAGTCAGAGGTGGCGAAACCCGACAGGACTATAAAGATACCAGGCGTTTCCCCCTGGAAGCTCCCTCG
TGCGCTCTCCTGTTCCGACCCTGCCGCTTACCGGATACCTGTCCGCCTTTCTCCCTTCGGGAAGCGTGGCGCTTT
CTCATAGCTCACGCTGTAGGTATCTCAGTTCGGTGTAGGTCGTTCGCTCCAAGCTGGGCTGTGTGCACGAACCCC
CCGTTCAGCCCGACCGCTGCGCCTTATCCGGTAACTATCGTCTTGAGTCCAACCCGGTAAGACACGACTTATCGC
CACTGGCAGCAGCCACTGGTAACAGGATTAGCAGAGCGAGGTATGTAGGCGGTGCTACAGAGTTCTTGAAGTGGT
GGCCTAACTACGGCTACACTAGAAGAACAGTATTTGGTATCTGCGCTCTGCTGAAGCCAGTTACCTTCGGAAAAA
GAGTTGGTAGCTCTTGATCCGGCAAACAAACCACCGCTGGTAGCGGTGGTTTTTTTGTTTGCAAGCAGCAGATTA
CGCGCAGAAAAAAAGGATCTCAAGAAGATCCTTTGATCTTTTCTACGGGGTCTGACGCTCAGTGGAACGAAAACT
CACGTTAAGGGATTTTGGTCATGAGATTATCAAAAAGGATCTTCACCTAGATCCTTTTAAATTAAAAATGAAGTT
TTAAATCAATCTAAAGTATATATGAGTAAACTTGGTCTGACAGTTACCAATGCTTAATCAGTGAGGCACCTATCT
CAGCGATCTGTCTATTTCGTTCATCCATAGTTGCCTGACTCCCCGTCGTGTAGATAACTACGATACGGGAGGGCT
TACCATCTGGCCCCAGTGCTGCAATGATACCGCGAGACCCACGCTCACCGGCTCCAGATTTATCAGCAATAAACC
AGCCAGCCGGAAGGGCCGAGCGCAGAAGTGGTCCTGCAACTTTATCCGCCTCCATCCAGTCTATTAATTGTTGCC
GGGAAGCTAGAGTAAGTAGTTCGCCAGTTAATAGTTTGCGCAACGTTGTTGCCATTGCTACAGGCATCGTGGTGT
CACGCTCGTCGTTTGGTATGGCTTCATTCAGCTCCGGTTCCCAACGATCAAGGCGAGTTACATGATCCCCCATGT
TGTGCAAAAAAGCGGTTAGCTCCTTCGGTCCTCCGATCGTTGTCAGAAGTAAGTTGGCCGCAGTGTTATCACTCA
TGGTTATGGCAGCACTGCATAATTCTCTTACTGTCATGCCATCCGTAAGATGCTTTTCTGTGACTGGTGAGTACT
CAACCAAGTCATTCTGAGAATAGTGTATGCGGCGACCGAGTTGCTCTTGCCCGGCGTCAATACGGGATAATACCG
CGCCACATAGCAGAACTTTAAAAGTGCTCATCATTGGAAAACGTTCTTCGGGGCGAAAACTCTCAAGGATCTTAC
CGCTGTTGAGATCCAGTTCGATGTAACCCACTCGTGCACCCAACTGATCTTCAGCATCTTTTACTTTCACCAGCG
TTTCTGGGTGAGCAAAAACAGGAAGGCAAAATGCCGCAAAAAAGGGAATAAGGGCGACACGGAAATGTTGAATAC
TCATACTCTTCCTTTTTCAATATTATTGAAGCATTTATCAGGGTTATTGTCTCATGAGCGGATACATATTTGAAT
GTATTTAGAAAAATAAACAAATAGGGGTTCCGCGCACATTTCCCCGAAAAGTGCCACCTGACGCGCCCTGTAGCG
GCGCATTAAGCGCGGCGGGTGTGGTGGTTACGCGCAGCGTGACCGCTACACTTGCCAGCGCCCTAGCGCCCGCTC
CTTTCGCTTTCTTCCCTTCCTTTCTCGCCACGTTCGCCGGCTTTCCCCGTCAAGCTCTAAATCGGGGGCTCCCTT
TAGGGTTCCGATTTAGTGCTTTACGGCACCTCGACCCCAAAAAACTTGATTAGGGTGATGGTTCACGTAGTGGGC
CATCGCCCTGATAGACGGTTTTTCGCCCTTTGACGTTGGAGTCCACGTTCTTTAATAGTGGACTCTTGTTCCAAA
CTGGAACAACACTCAACCCTATCTCGGTCTATTCTTTTGATTTATAAGGGATTTTGCCGATTTCGGCCTATTGGT
TAAAAAATGAGCTGATTTAACAAAAATTTAACGCGAATTTTAACAAAATATTAACGCTTACAATTTCCATTCGCC
ATTCAGGCTGCGCAACTGTTGGGAAGGGCGATCGGTGCGGGCCTCTTCGCTATTACGCCAGAGCTTGGCCGGATC
TAAAGTTTTGTCGTCTTTCCAGACGTTAGTAAATGAATTTTCTGTATGAGGTTTTGCTAAACAACTTTCAACAGT
TTCAGCGGAGTGAGAATAGAAAGGAACAACTAAAGGAATTGCGAATAATAATTTTTTCACGTTGAAAATCTCCAA
AAAAAAAGGCTCCAAAAGGAGCCTTTAATTGTATCGGTTTATCAGCTTGCTTTCGAGGTGAATTTCTTAAACAGC
TTGATACCGATAGTTGCGCCGACAATGACAACAACCATCGCCCACGCATAACCGATATATTCGGTCGCTGAGGCT
TGCAGGGAGTCAAAGGCCGCTTTTGCGGGATCATCACTGACGAATCGAGGTCGAGGAACCGAGCGTCCGAGGAAC
AGAGGCGCTTATCGGTTGGCCGCGAGATTCCTGTCGATCCTCTCGTGCAGCGCGATTCCGAGGGAAACGGAAACG
TTGAGAGACTCGGTCTGGCTCATCATGGGGATGGAAACCGAGGCGGAAGACGCCTCCTCGAACAGGTCGGAAGGC
CCACCCTTTTCGCTGCCGAACAGCAAGGCCAGCCGATCCGGATTGTCCCCGAGTTCCTTCACGGAAATGTCGCCA
TCCGCCTTGAGCGTCATCAGCTGCATACCGCTGTCCCGAATGAAGGCGATGGCCTCCTCGCGACCGGAGAGAACG
ACGGGAAGGGAGAAGACGTAACCTCGGCTGGCCCTTTGGAGACGCCGGTCCGCGATGCTGGTGATGTCACTGTCG
ACCAGGATGATCCCCGACGCTCCGAGCGCGAGCGACGTGCGTACTATCGCGCCGATGTTCCCGACGATCTTCACC
CCGTCGAGAACGACGACGTCCCCACGCCGGCTCGCGATATCGCCGAACCTGGCCGGGCGAGGGACGCGGGCGATG
CCGAATGTCTTGGCCTTCCGCTCCCCCTTGAACAACTGGTTGACGATCGAGGAGTCGATGAGGCGGACCGGTATG

TTCTGCCGCCCGCACAGATCCAGCAACTCAGATGGAAAAGGACTGCTGTCGCTGCCGTAGACCTCGATGAACTCC
ACCCCGGCCGCGATGCTGTGCATGAGGGGCTCGACGTCCTCGATCAACGTTGTCTTTATGTTGGATCGCGACGGC
TTGGTGACATCGATGATCCGCTGCACCGCGGGATCGGACGGATTTGCGATGGTGTCCAACTCAGTCATGGTCGTC
CTACCGGCTGCTGTGTTCAGTGACGCGATTCCTGGGGTGTGACACCCTACGCGACGATGGCGGATGGCTGCCCTG
ACCGGCAATCACCAACGCAAGGGGAAGTCGTCGCTCTCTGGCAAAGCTCCCCGCTCTTCCCCGTCCGGGACCCGC
GCGGTCGATCCCCGCATATATGAAGTATTCGCCTTGATCAGTCCCGGTGGACGCGCCAGCGGCCCGCCGGAGCGA
CGGACTCCCCGACCTCGATCGTGTCGCCCTGAGCGTCCACGTAGACGTTGCGTGAGAGCAGGACTGGGCCGCCGC
CGACCGCACCGCCCTCACCACCGACCGCGACCGCGCCATGGCCGCCGCCGACGGCCTGGTCGCCGCCGCCGCCCG
CCGGTTCGGCGCCTGACCCGACCAACCCCCGCGGGGCGCCGGCACTTCGTGCTGGCGCCCCGCCCCCACCCACCA
GGAGACCGACCATGACCGACTTCGACGGACGCCTGACCGAGGGGACCGTGAACCTGGTCCAGGACCCCAACGGCG
GTGGCTGGTCCGCCCACTGCGCTGAGCCCGGTTGCGACTGGGCCGACTTCGCCGGACCGCTCGGCTTCCAGGGCC
TCGTGGCCATCGCTCGCCGACACACGCACTGACCGCA

#### 7.3 pWHM4*::*ika*

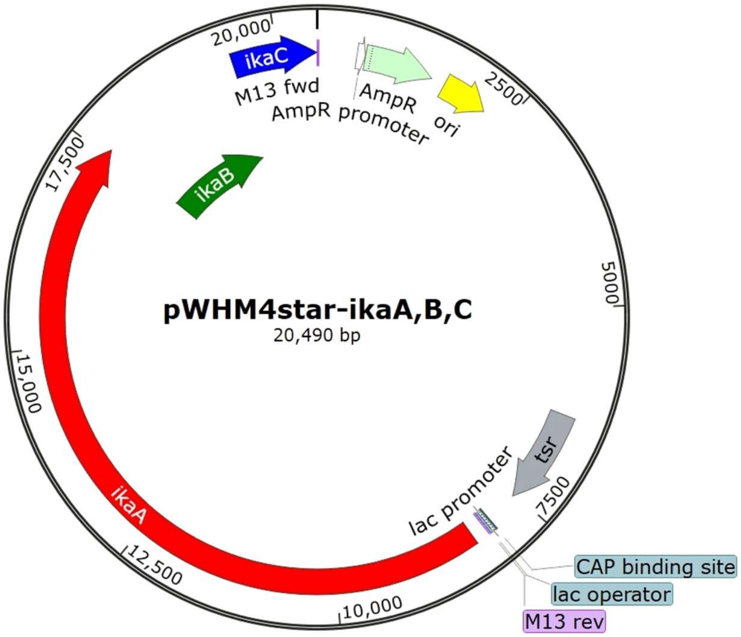

**Figure 4.** Vector map of *ika* in pWHM4*.

- DNA sequence of pWHM4*::*ika* [20490 bp]:

AATTCACTGGCCGTCGTTTTACAACGTCGTGACTGGGAAAACCCTGGCGTTACCCAACTTAATCGCCTTGCAGCA
CATCCCCCTTTCGCCAGCTGGCGTAATAGCGAAGAGGCCCGCACCGATCGCCCTTCCCAACAGTTGCGCAGCCTG
AATGGCGAATGGCGCCTGATGCGGTATTTTCTCCTTACGCATCTGTGCGGTATTTCACACCGCATATGGTGCACT
CTCAGTACAATCTGCTCTGATGCCGCATAGTTAAGCCAGCCCCGACACCCGCCAACACCCGCTGACGCGCCCTGA
CGGGCTTGTCTGCTCCCGGCATCCGCTTACAGACAAGCTGTGACCGTCTCCGGGAGCTGCATGTGTCAGAGGTTT
TCACCGTCATCACCGAAACGCGCGAGACGAAAGGGCCTCGTGATACGCCTATTTTTATAGGTTAATGTCATGATA
ATAATGGTTTCTTAGACGTCAGGTGGCACTTTTCGGGGAAATGTGCGCGGAACCCCTATTTGTTTATTTTTCTAA
ATACATTCAAATATGTATCCGCTCATGAGACAATAACCCTGATAAATGCTTCAATAATATTGAAAAAGGAAGAGT
ATGAGTATTCAACATTTCCGTGTCGCCCTTATTCCCTTTTTTGCGGCATTTTGCCTTCCTGTTTTTGCTCACCCA
GAAACGCTGGTGAAAGTAAAAGATGCTGAAGATCAGTTGGGTGCACGAGTGGGTTACATCGAACTGGATCTCAAC
AGCGGTAAGATCCTTGAGAGTTTTCGCCCCGAAGAACGTTTTCCAATGATGAGCACTTTTAAAGTTCTGCTATGT
GGCGCGGTATTATCCCGTATTGACGCCGGGCAAGAGCAACTCGGTCGCCGCATACACTATTCTCAGAATGACTTG
GTTGAGTACTCACCAGTCACAGAAAAGCATCTTACGGATGGCATGACAGTAAGAGAATTATGCAGTGCTGCCATA
ACCATGAGTGATAACACTGCGGCCAACTTACTTCTGACAACGATCGGAGGACCGAAGGAGCTAACCGCTTTTTTG
CACAACATGGGGGATCATGTAACTCGCCTTGATCGTTGGGAACCGGAGCTGAATGAAGCCATACCAAACGACGAG
CGTGACACCACGATGCCTGTAGCAATGGCAACAACGTTGCGCAAACTATTAACTGGCGAACTACTTACTCTAGCT
TCCCGGCAACAATTAATAGACTGGATGGAGGCGGATAAAGTTGCAGGACCACTTCTGCGCTCGGCCCTTCCGGCT
GGCTGGTTTATTGCTGATAAATCTGGAGCCGGTGAGCGTGGGTCTCGCGGTATCATTGCAGCACTGGGGCCAGAT
GGTAAGCCCTCCCGTATCGTAGTTATCTACACGACGGGGAGTCAGGCAACTATGGATGAACGAAATAGACAGATC
GCTGAGATAGGTGCCTCACTGATTAAGCATTGGTAACTGTCAGACCAAGTTTACTCATATATACTTTAGATTGAT
TTAAAACTTCATTTTTAATTTAAAAGGATCTAGGTGAAGATCCTTTTTGATAATCTCATGACCAAAATCCCTTAA
CGTGAGTTTTCGTTCCACTGAGCGTCAGACCCCGTAGAAAAGATCAAAGGATCTTCTTGAGATCCTTTTTTTCTG
CGCGTAATCTGCTGCTTGCAAACAAAAAAACCACCGCTACCAGCGGTGGTTTGTTTGCCGGATCAAGAGCTACCA
ACTCTTTTTCCGAAGGTAACTGGCTTCAGCAGAGCGCAGATACCAAATACTGTTCTTCTAGTGTAGCCGTAGTTA
GGCCACCACTTCAAGAACTCTGTAGCACCGCCTACATACCTCGCTCTGCTAATCCTGTTACCAGTGGCTGCTGCC
AGTGGCGATAAGTCGTGTCTTACCGGGTTGGACTCAAGACGATAGTTACCGGATAAGGCGCAGCGGTCGGGCTGA
ACGGGGGGTTCGTGCACACAGCCCAGCTTGGAGCGAACGACCTACACCGAACTGAGATACCTACAGCGTGAGCTA
TGAGAAAGCGCCACGCTTCCCGAAGGGAGAAAGGCGGACAGGTATCCGGTAAGCGGCAGGGTCGGAACAGGAGAG

CGCACGAGGGAGCTTCCAGGGGGAAACGCCTGGTATCTTTATAGTCCTGTCGGGTTTCGCCACCTCTGACTTGAG
CGTCGATTTTTGTGATGCTCGTCAGGGGGGCGGAGCCTATGGAAAAACGCCAGCAACGCGGCCTTTTTACGGTTC
CTGGCCTTTTGCTGGCCTTTTGCTCACATGTTCTTTCCTGCGTTATCCCCTGATTCTGTGGATAACCGTATTACC
GCCTTTGAGTGAGCTGATACCGCTCGCCGCAGCCGAACGACCGAGCGCAGCGAGTCAGTGAGCGAGGAAGCGGAA
GAGCGCCCAATACGCAAACCGCCTCTCCCCGCGCGTTGGCCGATTCATTAATGCAGGGTCGCCGGCTTCGTCGGC
GACGCCGTCCTGCTCCCCGGCACCGTCGACGTCGACTCCGGCGACACGGCCCGGACCGCCTTGGGCACGGTGCCC
CTCGCCGCTCCGCCCGGCGGCCTGCGCGCCGGCACGGTCCTGCTCCGTCCGGAACAACTCCGCCTCACCGGCGAG
GACTCCGTCGGCGCGGTCAGGGCAACGGTCACCGACGTCTCGTACTACGGCCACGACGCCATGGATCTGGGCCGC
GCGACGTACTGCTACGTGAGACCCCGCCGCCCGCCGCTGACCAGCACGAAGGGACCGCAGATGGAACTCATGAGA
CCGCCCTACGCGCGCGCGGACCGGTGTAACGCCTGGGCCCGCCACTCCGTCCATACAGCGCCGAGAGAGGAGACC
GCACACGTGGCACAGCAGGCAGTCCGAGCAGGGGGAACAACCGCCCGCCAGCGCCCTACGCTCGGAGACGGCGCG
GGGCCGACCCTCCAAAGGAGGAATGCCCCCGTGCGCCCAGTCGCTTCGCCCCGACCCACGCCCGACTTCCCCGGA
CTGTTGGTCTGCGCCGAACCCCCACGCGCCTTCTTCGCCTGCGCCTGCGGCTACTCGCGAGACGTCGCCGGCCGC
GCCCGTGTGGTCGAGCTGGTGACTCACCACCAGGAGCACGCCGACGTCTGCGTCCTGACGACCGGTGGGGCCTCT
GAGGCATCCGCGCGGCGAGCCACGCGTGAGGCGGGCGATGCGGCAGGATCTACCCGAACAACTCCAGAAAAAGCG
AAACGGCCAGGGGTCCTAATCCCTGACCGCTTCTGACGTCCACCCGGCTACCAACCAAGGGACTCGTCTGTGTCC
AGCGTAAGGGACGCTTTGGCGTCCGCAACCCCCTCTTTGATCGGCGCGCCGACTCAGTCGCCCGCCCGCGTCCCC
GCCGGCCCCCGTTGCCGCTGCGGCGCCGCCCTCACGATCACCCCCGGCAAGCGAGCCAAGGAGTTCTGCTCCGAG
GCGTGCAAGAAGCGGGCCGCGCGGGCCCGGAAGGCCGCCGACCGAGCCGCCGAGGCCGCCCGTGAGACCGCCCCT
GGCAAAAAGGGACGCCTAGGTAAAGGGTTGACAGAGTTTTCCCACCCCCCACTGACCAGCGACTTTGCGGAAGTG
GCAGTCGTGGGAGGAACGGGAGAGTCCGAGAGGGCGGCAGCCCTTTCGGAACAGACCACAAGGCGCGCGGCGACC
GCTGCCGTACGCCGCTACCAGGGCCGCAAGGTGCTCAACCGGGTCTCCGGGATCGACGCCTGCGGCGGGTGCGGG
CGGCGGGTCCTCGACCCGGACACCGGCGTGATCTACGCGAAGTCGAGCCGTGGGTACGTCGTCACGATCGGCCTG
GTCCGCTGCGGGCGGATCTGGTTCTGCCCGGAGTGCTCCTCCGCGATCCGCCGTGGCCGGACCGAGGAGATCAAG
ACCGGTGCTCTGCGGCACCTCGCCGCCGGCGGCACGCTCGCCGTTGTCGTCCTCACCGCCCGGCATAACCAGACC
ACCGACCTCGACAGCCTGGTCGCCGCGCTCTGGGGCGGGCCTCTCCTGGACGACAAGGGCGCCCCGGTCCTCGAC
CGGTCGGGCAAGCCCCGCCGCGCGCCCGGTGCCTACCAGCGGATGCTCACGGCCCCGGCCTTCTACGGCCGCCCT
GAGGCCCGCCGCACCCGGAAGGACGGAACGCAGTACGTCCGTCCCGCTGAGGACGGCATCCGCCACCGGATCGGC
TACATCGGCATGGTCCGCGCGGCTGAGGTCACCCGGTCCAAGAAGAACGGTTACCACCCCCACCTCAACCTGCTG
GTCTTCCTCGGCGGCGAGCTCTCTGGCACCCCGGCCAAGGGTGACGTCGTCGGACACTTTGAGCCCTCCGAGACG
GACCTGGGGGACTGGGAGGACTGGCTCCGGGAGATGTGGGCGGGCGCCCTCAAGCGGGCTGACCCCAAGTTCGAG
CCCTCGACCGACTGCGACACCCCCGGCTGCAAGTGCAAGGGCAAGGGCCACGGCGTGATGGTCTCGATCGTCCGG
TCAGCTGACGACGTCGCGCTGATCGAGTACCTCACCAAGAACCAGGACGGGAAGCGAGAGCGGCCCGACTCCGTC
GACCAGGACCTCGAAGCCGCCGGCGCAGCTGCGATGGAGACCGCCCGTCTGGACTCCAAGACCGGCCGGGGCCGG
AAGTCCATGACGCCGTTCCAGATCCTCTACCGACTGTGGGACATCGAGGTCGCCGGGCTCGACCCCGACATGGCC
GAGGGCTACGGCACGCCGAAGCAGCTGCGCGCGTGGTGGGCCCAGTACGAGGAAGCCCTCGCCGGACGACGCGCG
ATCGAGTGGACCCGAGGCCTGCGGCGGCACGTCGACCTCGACGGTGACGACGACGAGGAGACCGACCTCCAGTAC
GTCTACGAGCCGGAGGCCGCGCCGCTCGACGGTGGCGTCGTCCTCACCTCCGACGCGATGCGCCTGGTCGTCGGA
GCCGACGCTGAACTCGACCTCGACGACGTCGTCCGCGCGGAGGCGTACTACTCCGCGGTTGACGTCGTCACCGGT
CTCGGAGGACGTGCGGATCACGTGCGGGTCGCTACCGCCGAGGAACTCGCGGAGGTGCAGGAGGTGCTGTTCGCG
CGGACGCAGGAACGCGCCGAGGAGAGCAGGCGCCAGCGCCGAATCGCGGAGCACGAGGCCGAGCAGGCCGCCGCG
CATCGGAAGCGGCAGGAGCTCGCGCGGTGCCTCGGGCTGCTCGTACGGCAGCGCGGCGGCACGCAGGACGACTCG
GCGGCGGACAACTTCGTCGCGCACATCCACGCGAACCGCTGACGACAACCCCCGGTTGTCGCCCCGACCGTCCCC
GGTCGACGCCGAAAACCCCGGCGTGACCGAGGCGGCGGGGCGTAACTGGTACCAGTTACCTATAACTCGCTTCCG
CTCCCTACTGCGAGAAGCCCCGGGGCCCGAGAGGTCGGCGGCCCACGATGACGGGCCGCTCGATCCGCTCCGGGA
CCTCCACCCAGTCACCGTCCAGCGCGCGCTCGATCGCCGTCAGCCGACGCTTATCTCGCTCGCGGATCTGGACAT
CGGGGCCTGCGCCGAGAGGGCGTCGTGCTGTGCTCATGGCGCCAGTCTTCCGTGCCGTGAAGGCGTACGGCTTCG
CCGCCGACTTCGTCGTGACCTGAGCGGTCACACCTCAGGCATCACGAATGCCCTCGGGTGCCGACGCCCCGCCGG
AACGGTGGTCCAAGGCGTGCACACGTGCCCGTGCACACGCGTGCACGCGAGCCGTATGGCCCCGGCCATACCTTG
GTACTTCCGGAGCCGTCGTACCGGTCGCAAGCCCGGCGCTCTGACCTGGTCGAGGTACATCTCGTGGAACTTCGT
GGAACCGTCCTACGCGCGCGCGTGCAACGGTTCCTCGCCGTGTGCTGTCGGCTCTCGTGAACGGCCGGTTCGTCG
TGATGCATTGGTGTGGGCGATGAGTTCGAGCGCGTACTCCTGGTCGTCTTCGTGCTCGATCGCGGCCGCGCTCGT
TGGTCCGGGCATGAACGGCGACCAGGAGCACGAGCAGCCCACCGAGTTCGCACAGATCGAAGCGCGGATGATCCG
GACTGCGCTCACCGACGAGGAGTGGGCGGAGCTGATCGAGCACCGGCCGAGTCTGCCTGCCTCGCTCGCGGCGTA

CGAGCGGTGGGAGGCCGAGGTCGAGACGAAGGCCGGGTGACGGTGGCCGCGCGGCACGTACTAAGGCGCGGCGCA
TTCTGGTGACCGTGCTCCTGGTCGTCATCGGCGCCGGCGCCACCTCATCAGCGTCGCGGCCAGCTCCGGCAGCTC
GCGGATCACGGCGACGACGTCGGAGCGCTCCGCCCGCAGGATCGCGGTGATGGCGACGGCGGTGATCGCCAGGAT
CAACCTCACGGTCAACTCGGGTGGCGGCCGGGGCGGTCAGTCCCCCGCCGCCTCGCGTACGCCCCACTTCATGCT
GGGGGAGCCGACGTTCGCCTCGACCGATGACGTCCGCAACTACTGCAACGGCGTCCGCGCTCTGATGCTCCAGTG
CGCGATCAAGGCGAATACTTCATATGCGGGGATCGACCGCGCGGGTCCCGGACGGGGAAGAGCGGGGAGCTTTGC
CAGAGAGCGACGACTTCCCCTTGCGTTGGTGATTGCCGGTCAGGGCAGCCATCCGCCATCGTCGCGTAGGGTGTC
ACACCCCAGGAATCGCGTCACTGAACACAGCAGCCGGTAGGACGACCATGACTGAGTTGGACACCATCGCAAATC
CGTCCGATCCCGCGGTGCAGCGGATCATCGATGTCACCAAGCCGTCGCGATCCAACATAAAGACAACGTTGATCG
AGGACGTCGAGCCCCTCATGCACAGCATCGCGGCCGGGGTGGAGTTCATCGAGGTCTACGGCAGCGACAGCAGTC
CTTTTCCATCTGAGTTGCTGGATCTGTGCGGGCGGCAGAACATACCGGTCCGCCTCATCGACTCCTCGATCGTCA
ACCAGTTGTTCAAGGGGGAGCGGAAGGCCAAGACATTCGGCATCGCCCGCGTCCCTCGCCCGGCCAGGTTCGGCG
ATATCGCGAGCCGGCGTGGGACGTCGTCGTTCTCGACGGGGTGAAGATCGTCGGGAACATCGGCGCGATAGTAC
GCACGTCGCTCGCGCTCGGAGCGTCGGGGATCATCCTGGTCGACAGTGACATCACCAGCATCGCGGACCGGCGTC
TCCAAAGGGCCAGCCGAGGTTACGTCTTCTCCCTTCCCGTCGTTCTCTCCGGTCGCGAGGAGGCCATCGCCTTCA
TTCGGGACAGCGGTATGCAGCTGATGACGCTCAAGGCGGATGGCGACATTTCCGTGAAGGAACTCGGGGACAATC
CGGATCGGCTGGCCTTGCTGTTCGGCAGCGAAAAGGGTGGGCCTTCCGACCTGTTCGAGGAGGCGTCTTCCGCCT
CGGTTTCCATCCCCATGATGAGCCAGACCGAGTCTCTCAACGTTTCCGTTTCCCTCGGAATCGCGCTGCACGAGA
GGATCGACAGGAATCTCGCGGCCAACCGATAAGCGCCTCTGTTCCTCGGACGCTCGGTTCCTCGACCTCGATTCG
TCAGTGATGATCACCTCACACGGCAGCGATCACCACTGACATATCGAGGTCAACGGTCGTGGTCCGGGCGGGCAC
TCCTCGAAGGCGCGGCCGACGCCCTTGAACGACTCGATGGCGCTGACGCCTTCCGCGATCGCCTCGACTGTGAAA
ACGCTCGTGGACGTCGGCGGGTAGCCGATGAGTTCCATCGCCTTGCGCATCCCCTTGAGCATGGAGAACGCGTCC
ACCGGGTAGTCACGAAGGTACTTGCGAAGGCTTCGCGCCGTCCGAGACCGTGAGCCGGCGCAAGCCGGTGTGGCG
CCGGTCCCTGGCACGACAGGTTTCCCGACTGGAAAGCGGGCAGTGAGCGCAACGCAATTAATGTGAGTTAGCTCA
CTCATTAGGCACCCCAGGCTTTACACTTTATGCTTCCGGCTCGTATGTTGTGTGGAATTGTGAGCGGATAACAAT
TTCACACAGGAAACAGCTATGACCATGATTACGCCAAGCTTGCGAGTGTCCGTTCGAGTCGAGAGGTGCGGGGAG
GATCTGACCGACGCGGTCCACACGTGGCACCGCGATGCTGTTGTGGGCACAATCGTGCCGGTTGGTAGGATCCAG
ACCTGCAGGTCGACTCTAGAGGATCCCCGGGTACCGAGCTCGAATTCATGGATTCCATGCACCACCCTGCCCCCG
TCCCCGTACCCGAAGTCCCCGCGCCCGTCCCGTCCCAGGACGACGCGTTCGCCATCGTCGGCATCGGCTGCCGGC
TGCCCGGCGGCGCCAGCGACTACCGGACCTTCTGGCGCAACCTCCTCGACGGCAAGGACTGCATCACCGACACCC
CCGCCGACCGCTACGACACCCGCACCCTGGGCAGCGGCGACAAGGCCAAGCCCGGCCGGCTGGTCGGCGGACGCG
GTGGATACATCGACGGCTTCGACGAGTTCGACCCCGCCTTCTTCGGCATCAGCCCGCGCGAGGCCGAGCACATGG
ACCCCCAGCAGCGGAAGCTCCTGGAGGTCGCCTGGGAGGCGCTGGAGGACGGCGGCCTCAAGCCCGCCGAGCTGG
CCGGCAGCGATGTCGGGGTGTACGTCGGGGCGTTCACCCTCGACTACAAGATCCTGCAGTTCGCCGACCTCGGCT
TCGAGACCCTGGCCGCGCACACCGCCACCGGCACCATGATGACGATGGTGTCCAACCGGATCTCGTACTGCTTCG
ACTTCCGCGGACCCTCGGTCTCCGTCGACACCGCGTGCAGCGGCTCCCTGGTCGCCGTCCACCTCGCCTGCCAGA
GCCTGCGCCGCGGCGAGACCTCCGTCGCCCTGGCCGGCGGCACCCTGCTGCACATGGCGCCGCAGTACACCATCG
CCGAGACCAAGGGCGGGTTCCTCTCCCCCGACGGCCGCTCCCGCGCCCTGGACGCCTCCGCCAACGGCTACGTGC
GCGCCGAGGGCGTCGGCATGGTCGCCATCAAGCGCCTCGCGGACGCGCAGCGCGACGGCGATCCCATCCACGCCG
TCATCATCGGCAGCGGCGTCAACCAGGACGGCCGCACCAACGGCATCACCGTGCCCAACCCCGACGCGCAGGTCG
CCCTGATCGAGCGGGTCTGCGCCGCCGCCGGCGTCACCCCCGGCAGCCTCCAGTACGTCGAGGCGCACGGCACCT
CCACCCCCGTCGGCGACCCGCTGGAGGCCAACGCCCTCGGCCGCGCGCTCTCCATCGGCCGCGAGCCGGGCGCCC
GGACGTACGTCGGCTCGGTCAAGACCAACATCGGGCACACCGAGTCCGCCGCCGGCATCGCCGGGCTGATCAAGA
CGGTGCTCAGCCTCAAGCACAAGGTCATCCCGCCGCACATCAACCTGGAGAAGCTCAACCCGCAGATCGACGAGG
CGTCCCTGCCGTACGAGATCCCCCGCGAGCCCACCCCCTGGCCCGAGCACAGCGGGCCGGCCCGGGCCGGCGTCA
ACTCCTTCGGCTTCGGCGGGACCAACGCCCACGTCCTGCTCCAGGAGGCACCGCCGACCGTCGGGGAGCCCGCGC
CACCGGCCACCGACGGGTACTCCGTGCTGCCGCTCAGCGCCCGCGACCCCGAAGCCTTTCCCGCCATCGCCACCG
GCCTGCGCGAACGGCTCGCCGAGGGACTGCCGGTGGGCGACGCCGCCTACACCCTCGCCCACCGGCGGCAGCATC
TGGAGCAGCGGCTGTCCGTCGTGTACGACTCCCCCGAGGCCCTCGACGAGGTGCTCGGCGCCGTCGCCCGCGGCG
AGAGCCACCCGCGTGCCGTCGCCGGCACCCAGCGGGAGGGCCTGGACCGCAGGCTGGTGTGGGTGTTCACCGGCA
TGGGCCCGCAGTGGTGGGCCATGGGCCGCCAGTTGTACGCGAGCGAGCCCGTCTACCGGGAGGTCATCGACCGCT
GCGACCAGGAGATCGCCGCGCTCACCGGCTGGTCCCTCACCCAGGAGCTGAACGCCGACGAGGCCGACTCCCGGA
TGAGCGAGACCTGGCTCGCCCAGCCCGCCAACTTCGCCGTCCAGATCGCCCTGGCCGCCCTGTGGCGCAGCAAGG
GGATCCAGCCCGACGCCGTCACCGGGCACAGCACCGGTGAGGTCGCCGCGTTCTACGAGGCCGGGGTGTACACCC

TCCCCGAGGCCGTGAAGATCGTGGTGCACCGCAGCCGGCTCCAGCAGAAGCTCATCGGCACCGGCTCCATGCTCG
CCGTCAGCCTCACCGAGGCGGAGGCCGCCCGCCGGGTGCGCCCGCACGGCGACCGGGTCTCCATCGCCGCCGTCA
ACAGCCCCACCTCCATCACCCTGGCCGGGGACACCGAGGCGCTGGAGGTGATCGCCGCCGAGCTGGGCGCCGAGG
ACATCTTCGCCCGCTTCCTGGAGGTCGGCGTCCCGTACCACAGCCCCCGCATGGAGCTGATCAAGGACGAGCTGC
TGACCTCGCTCGCCGATCTCAAGCCGCAGCAGGCGAAGTTGCCGCTGTACCTCACCGCGCTGCCGGGCACCGTCG
CGCAGGGCACGGAGCTGGACGCCGACTACTGGTGGCGCAATGTGCGCGAGGCCGTGCACTTCCGGGCCGCCGTGG
ACCGGCTGCTGGACGACGGCTACGGCGTCTTCCTGGAGATCGGCCCGCACCCCGTGCTCGCCCACTCCCTGCGCG
AGTGCTGCGAGGCCCGCGACGCGCACAGCGTCACCCTGGCCTCCATCCGCCGCAAGGCGGACGAGCGCGAACGCC
TCACCCTGTCGCTCGCCGCGCTGCACAGCCTCGGCTTCGCCGTGGACTGGCACGCCCTGCACCCCGCCGGGCGGC
CGGCCGAACTGCCGCGCTACCCGTTCCGGCGCGACCGGTACTGGGTCGAGCCGGCCCCGGTCGCGCAGATCCGGC
TCGGCCACCGCGACCACCCGCTGCTGGGCCGCCGCACCGCGAGCGCCGAGCCGGTGTGGGAGGTGAAGCTGGACG
CGGAGGCCGCCCCGTACCTGGAGGACCACCGCATCCAGGGCACCGTGCTGTTCCCGGCCGCCGGCTATCTGGAGA
TGGCCGCGCAGGCCATGCGGGCGCTGACCGGTGATGAGCACAGCACCGCCGCGCTGGCCGGCATCGAGCTGCGCA
AGGCGCTGTTCCTGCCGGACGGCGAGCCGCAGACGGTGCAGCTGTCCTTCTCCTCCGACGCCGCCGCGTTCTCCA
TCGCCACCGTGGGCGCCGCCGGCGCCGAGCCGACCGTGCACGCCACCGGTACGGTACGGGCCGCCCAGCGCCGCC
GGCTGACCGCGCCGCTGGACACCGTCGCCGTCCGGGCCCGCGCCGCCCGCCACCTGAGCGGCCCCGACTGCTACG
CCGAACTGGCCGCGCTCGGCTACCACTACGGCCCCGCCTTCCAGGGCATCGAGGAGGTGTGGATCGGCGAGGGCG
AGGCCCTGGCCCGGATCCGTCCGCCGCAGGGGCTCACCCCGGACGCGGCGGCGCACCACATGCATCCGGTGCTGC
TCGACTCCTGCTTCCAGTCGCTGCTGACCCCGCAGCTGCTCACCGCGCCCGCCGGGCCCGGGGGCACCGGCATCC
GGCTGCCGCTGTCCATCGCCGAGGTACGGCTGGACCCGGTCGGCGACCGCGAACTGTGGGTGCACGCCACCGTCA
CCGGCGACGACGAGGACGAACTCACCGGTGACATCGCCGTGTACGACGGCGCCGACGGTACGCCGCTGGGCCGCG
TCGCCGGCTTCCGCGCCGCCGATGTGGAGAAGGCCGCCACCACCGTGGGGCTGTCCACCATCGACAGCTGGCTCA
CCGAACCGAGCTGGGTGCCGTGCCCGCTGCCCGAGGCGGCGTCCGCCGCGCCGGCGGCCGGGCGGCACGTACTGT
TCGCCGACGCGGGCGGGGTCGCGCAGCGGCTGGCCGCGCTGATCGGCGAGGCCGGCGGGGAGGCCCATCTGGTCC
GGCCCGGTGCCGCGTACGGCCTGGACCGCACGGCGAGGACCGCCACCGTCGTCCCCGGATCCGCGGATGACCTGC
GGCGGTTGCTCACCGATCTCGGGCAGGTGGACGGCGTCGTCCACCTGTGGAACCTGGACCGGCCGGCGCTGGCCG
ACGCCCCGCGCGGACGGTTCGCGGACATCGCCTCCACCGGCGCGTACGCCCTGATCGCCCTCACTCAGGCCCTGC
TCGCCGACCCGGAGCGGCACGGCGGCACCCCGGTGCACATCGTCACCAGAGCCGCCCAGTGCGTGGTCCCCGGTG
AGCCGGTGGAGCCGCTGGGCGCGCCCGCCTGGGGCATCGGCCGGGTGCTGTGGCAGCAGGAACTGGCCGGGCGCG
GCGGCAAGCTGATCGACCTGGCGGCCGACGGCGGCGTCGAGGAGGACGCGTACGCGCTGCTGCGCGAGCTGGCCG
ACCCCACCGGCGCGGCCGAGCGCGAGGACGAGATCGCGCTGCGCGCCGGGGAGCGGCACACCAGCCGGCTGGTGG
CCGCCGAGGGGCTGAGCAGGCCGCTGCCCCTGCGGCTGCGCCCGGACGGCAGCTATCTGGTGACCGGCGCGTTCG
GCGCGCTCGGCAGGCTGCTGTGCCGCACGCTGGTCAGGCGCGGGGCGCGGCGGCTGATCCTGGTGGGCCGCACCC
GGCTGCCGGAGCGCGAGCGCTGGGCCGACCAGGACCCGAACTCGCCGGCCGGGCGGCACGTGGCCTTCCTCAAGG
AGCTGGAGGCGCTGGGCGCGCAGCCGATTCTCGCGCCGCTGGACATCACCGACGAGGACGCGCTGGCCGGCTGGC
TCGCCGGGTACCGGCGCGCCCAGGGGCCGCCGATCCGCGGGGTGTTCCATCTGGCGGGGCAGGTGCGCGACACCC
TGGTGCCGGAGATGGACCGGGAGGTGTTCGACGCCGTCCACGACCCGAAGGTGGTGGGCGCGGCGCTGCTGCACC
GGCAGCTGAGCGGCGAACCGCTGGAGCACTTCGTGCTGTTCGCCTCGGTCGCGGCCTGGCTGACGACGGCCGGAC
AGACCAACTACGCGGCGGGGAACGCCTTCCTGGACGCGCTGGCGCACCACCGCCGCGCGCAGGGGCTGCCGGCGC
TGGCGCTGGACTGGGGCCCGTGGGCCACCGGCATGATCGAGGAACTGGGCCTGATCGACCACTACCGCAACAGCC
GGGGCATGTCCTCGCTGGCGCCCGAGGCGGGCATGGCGGTGCTGGAGCGGGTCATCGGGCAGGACCGGGCACAGC
TGCTGGTGGCCACGGTCGTGGACTGGCCGGTGTTCATGTCCTGGTACGCGGCGCCGCCGCGGCTGGTCACGGAGC
TGGCGGCCACCGCCCAGGGACCGGGGTCCGAGGGCGACGGCAGTTTCCTGGACGCGTTCCGGGAGGCCACCGCGG
ACAAGCGGCGGCTGCTGCTGACCGAGCGGTTCACGACGCTGGTGGCGGGTGTGCTGCGGGTGCGGGCCGAGCAGG
TGGATCCGGCGGTCAGCCTGAATCTGCTGGGGCTCGACTCGCTGCTGGCGATGGAGCTGCGAGCGCGGGTGGTGG
CCGAGGTGGGCATCGCGCTGCCGGTGGTGGCGCTGCTGTCCAGCGCGCCGGCCGGGGACCTGATCACCCAGCTGC
ACGAGGGCCTGGAGGAGTTGCTGGCCGAGGAGGGCAGCGGCGCCGCGGTGACGGCGGTGGAGCGCTTCGAGGACG
AGGCCGAGTTCCCGCTGACGCAGAACCAGAAGGCGCTGTGGTTCCTGAAGCAGCTGAACCCGGACGGCTTCGCGT
ACAACATCGGCGGCGCCGTCGAGGTGCGGGTCGAGCTGGACCCGGACCTGATGTTCGAGGCGTTTCGCCGGCTGC
TGGCCCGGCATCCCGTGCTGCGGGCGAACTTCCTGCTGGTGGAGGGGCAGGCGGTGCAGCGGATCTCCCCGGAGA
TCAAGGAGGACATCGCGCTCTTCGACGTCGAGGACCGCGCGTGGGACGACATCTACCGGATGATCATCGAGGAGT
ACCGCAAGCCGTACGACCTGGCGACCGATCCGCTGATCCGGTTCCGCCTCTTCCGGCGCGGCCCGGACCGCTGGG
TCATCACCAAGGCCGTCCACCACATCATCTCGGACGCCATCTCCACCTTCACCTTCATCGAGGAACTGCTGTCCC
TGTACGAGGGGCTGCGGCAGGGCCACGACGTCGAACTGCCGCCGGTGTCCGCCCGCTATCTGGACTTCCTCAACT

GGCAGAACGCGTTCCTGGCCGGCCGCGAGGCGCAGAAGATGCTCGCGTACTGGCGGGGGCAGCTGCCGGACGAGG
TGCCGGTGCTGGCGCTGCCCACCGACAAGCCGCGCCCGGCGGTGCTCACCCACAACGGGGCGTCCGAGTTCTTCG
CCCTGGACGCGGAGTTGAGCGCCCGGGTGCACGCGCTGGCGCGGGAGCACAACGTCACCGTCTTCATGGTGCTGC
TGAGCGCGTACTACCTGCTGCTGCACCGCTATGCGGGGCAGGACGACATCATCGTCGGCTCCCCCGTCACCGGCC
GCACCCAGGAGGAGTTCGGCGCCGTCTACGGGTACTTCGTGAACCCGCTGCCGCTGCACGCCTCGCTGGCCGGTG
ACCCCACGGTCGCCGAGCTGCTGGACCAGGTGCGCACCACGGTGCTGGGCGGCCTGGACCACCAGGAGTACCCGT
TCACGCTGCTGGTGGAGCAGCTGGGGCTGGCCCACGACCCGAGCCGGTCGGCGGTCTTCCAGGCGATGTTCATCC
TGCTGCACCACAAGGTGGCCACCGAGAAGTACGGCTACAAGCTGGAGTACATCGAGCTGCCCGAGGAGGAGGGCC
AGTTCGACCTGACGCTGTCCGCGTACGAGGAGGAGGCGGACGGGCGGTTCCACTGCGTCTTCAAGTACAACACCG
ACCTCTTCGAGGCGGAGACGATCCGGCGGCTCGCCGGGCACTACACGCAGCTCCTGGAGTCGCTGACCGCGGCGC
CCGCCGACGCCGCCACCGGTGGACTGCGGATGCTGTCGGGCGGCGAGCGGGAGCGGATCCTCACCGAGTGGAGCG
GGGCCGGGCAGGGCGCGCAGGACGCGCCGGTGCCGGTGCACCGGCTGATCGCCGAGGCGGCGCACCGTACCCCGC
AGGCGATCGCGGTGGCCGCGCCCGCCGAGAGCGGGGAGACCCGGCGGCTGACGTACGGCGAACTGGAGGAGCGCG
CCGGCGAACTGGCCGGGCGGCTGCGGGCGCGCGGCGTGCGCGAGGGCACCGTCGTCGCGCTGTGCCTGGAGAAGT
CGCCCGAGCTGATCACCGCCCTGCTGGCGGTCCTCAAGGCGGGCGGCGCCTATCTGCCGCTGGACCCGGACTATC
CGGCCGACCGGCTCGCGTACATGGTGCGCAACGCCGGGGCCACGCTGGTGATCGGCGGGACGGGCGGCGCGGCCG
AGGGGCTGCCGGGCACCGTGGTCACCCTGGAGGAACTGCTCGCGGGCGAGGCCGGCGAAGCGGGGCCGGACGCCG
AGCCGGGGCCCGACTCCCCCGCCTACGTCATCTACACCTCGGGCTCCACCGGGCGCCCCAAGGCGGTCGCGGTCA
GCCACCGCAATCTGGCCTCGGTGTACGCCGGATGGCGCGACGCCTACCGCCTGGAGGAGGGCGGCATCCGGGTCC
ATCTCCAGATGGCCAGCCCCTCCTTCGACGTCTTCACCGGCGACCTGACCCGAGCCCTGTGCTCGGGCGGCACGC
TGGTGCTGGTCGGCCGGGAGCTGCTGTTCAACACCGCCCGGCTGTACGAGACGATGCGCGCCGAACGGGTGGACT
GCGGCGAGTTCGTGCCCGCCGTGGTGCGCACCCTGGTGCGGCACTGCGAGGACACCGGCGCCCGGCTGGACTTCC
TGCGGCTGCTGATCGTGGGCTCGGACTCCTGGAAGGCCGAGGAGTACGAGCGGCTGCGCGCGCTGGGCGCACAGC
GCCTGGTGAACTCGTACGGGCTCACCGAGGCCACCATCGACAGCGCCTGGTTCGAGGGTCCCGCGGATGACCTGG
AGGGCGGCCGGATGGTGCCCATCGGGCGGCCGTTCCCGGGCAGCGCGCTGTACATCCTGGACTCGCGCGGCGAGC
CGGTGCCGCCCGGTGTCCCCGGCGAGCTGTGGATCGGCGGCACCGGGGTGGCGCTCGGCTACCTCGGCGACGAGG
CGCTGACCGGGGAGCGGTTCCTCACCCGCGCCCTGGCCGGCGACGCTCCGGTACGGCTGTACCGCACCGGTGACC
TCGCGCGCTGGGACGCGGCCGGCACCGTCCATCTGCTGGGCCGGGCCGACTCGCAGATCAAGGTGCGCGGGCACC
GCATCGAGATCGGGGAGATCGAGTCGCACCTGGCGGCCTGCCCCGAGCTGGCCCAGGCGCAGGTCACCGTGCGGC
CGGACGCGGGCGGCGAGAACGTGCTGTGCGCGTACGGGGTGGCGGCCCCGGGCGCCGTGCTGGACTGGCGCGAGG
TGCGCCGGCGCCTGGCGGACTATCTGCCGACGTTCATGATCCCCACCCACTTCACCGAGCTGCCCGCCCTGCCGC
TCACCCCGAACGGCAAGGTGGACGTGGCGGCGCTGCCCGCCCCGCGCACCGGCGACGGCGCGGACGGGCCGGTGT
ACGAGGCCCCCGTCACGCTGTACGAGACCCGGATGGCCGAGCACTGGCAGCGGCTGCTGGGCATCGAGGCCCCCG
GGCCCGGTCTGGGCCACGACTTCTTCGAGACCGGTGGCAGCTCCATCCGGCTGATCGAGCTGATCTACCACCTGC
AGGCCGAGTTCGGGATCTCCATCCCGGTCAGCCGGCTGTTCCAGGTGACGACGCTGCACGGCATGGCCAAGACGG
TCGAGCGGATCGTCACCGGGGAGATCGAGGGGTCGCTGCCGTATCTGCGGTTCAACGAGAACGCCGCGGCGGGCA
CGGTGTTCTGCTTCCCGCCGGCCGGTGGCCACGGCCTGGTCTACCGGGAGTTCGCGGCGCGGCTGCCGGAGTTCG
AGTTCCTCGCCTTCAACTACCTGATGGGCGAGGACAAGGTAAGCGGGTACGCCGACCTGGTGGCCGGGCACCGGC
CGGAGGGCGAGATCGACCTGCTCGGCTACTCGCTGGGCGGCAACCTCGCCTTCGAGGTGGCCAAGGAGCTGGAGC
GGCGCGGCCGCACCGTGCGCCACGTCGTCATCATGGACTCGCTGCGGGTGACGGAGTCCTACGAGCTGGGCCCGG
AGCACCTGGCCGTCTTCGAGCGCGAGCTGGCCGAGCATCTGCGCAAGCACACCGGCTCGGCGCTGGTCGCGGAGA
AGACGCGCGAACAGGCCAAGGACTACCTGGAGTTCACCGGCCGCACCGCCAACCCCGGCACCACCGGGGCCCGGA
TCGCGGTGATCAGTGACGAGGAGAACGCGGCCGCGTACGACAGCGGCGCCGAGGGCAGCTGGCACGGCGCCTCCC
GTACCGGAACCGACGTGCTGCGCGGGGTGGGCCGGCACGCCGACATGCTCGATCCGGGGACGGTCGAGCACAACG
CGCGCCTGGCGCGCGGCATTCTCACCGGCGGTGATGGCGAGGTATGACGCCTTTCGTTCAGCCGGCGGTCGACAC
CAAGGAGCACAGCGCCATGTCATCCCCCACCACCTCCGGCACCCCGGGCAGGCAGTCGATGATCATCATCGGCGG
CGGCCTGGGGGGCCTGTCCACCGGCTGCTACGCGCAGATGAACGGCTACGCGACGCGGGTCTTCGAGATGCACGA
GATCCCGGGCGGTTCCTGCACCGCCTGGGAGCGCGGGGACTTCACCTTCGACTGGTGCGTCAGCTGGCTGCTGGG
CAGCGGTCCCGGCAACGAGATGTACCAGATCTGGATGGAACTGGGGGCGTTGCAGGGCAAGGAGATGCGCCAGTT
CGACGTCTTCAACATCGTGCGGGTGCGCGGCGGCCAGCCGGTGTACTTCTACTCCGACCCGGACCGGCTCCAGGC
GCACCTGCTGGAGATCTCCCCGGCCGACGCCCGCCGCATCAAGAACTTCTGCGAGGGGGTGCGCACCTTCCAGAA
GGCGCTGTCGGTCTACCCGTTCCTCAAGCCGGTGGGGCTGATGGGGCGGTGGGAACGGTGGAAGATGCTGGCCTC
GTTCCTGCCGTACTTCAACGCCATCCGCAAGTCCATCACCGAGCTGATGACGGACTACGCGGAGAAGTTCCAGCA
CCCGGTGCTGCGCGAGGCCTTCAACTACGTGCTGTACGAGAAGCACGCCGACTTCCCCGTCCTGCCGTTCTGGTT

CCAGCTGGCCTCGCACGCCAACGGCTCGGCGGGGGTGCCCGAGGGCGGCTCGCTGGAGCTGGCCCGGTCCGTGGA
GCGGCGCTACCTGGGGCTCGGCGGGGAGATCACCTACAACGCCAAGGTGGAGAAGATCCTCGTCGAGCACGACAA
GGCGGTGGGAGTGCGGCTCACCGACGGCCGCGAGTTCCGCGCGGACATCGTGGTGTCGGCGGCCGATCTGCACAC
CACCGCCATGGAGATGCTCGGCGGCCGGTATCTCAACGACACCTGGCGCAAGCTGCTCACCGAGACGATCGACGA
GGTGGGCACGATCTCCCCCGGCTATGTCTCGCTGTTCCTGGGGCTGCGCCGGCCGTTCCCCGAGGGCGAGCCGTG
CACCACGTACGTGCTGGAGGACAGCATGGCGGAGAAGCTCACCGGCATGCGGCATCCCAGCATGAACGTGCAGTT
CCGCAGCTGCCACTACCCGGAGCTGTCGCCGCGCGAGACCACGGTCATCTTCGCCACGTACTTCTCGGAGGCCGA
GCCGTGGCGGGCGCTGCGCGACGACGTGCCGGAACAGGCGGGCCGGGTGCGGCGCGGTCAGGTGCTGCACACCCT
GCCGGTGAAGCACGGCAAGGCGTACACCCAGGCCAAGCGGCAGGCGCGGATCACCATCGAGAACTTCCTGGACGA
GCGGTTCCCCGGTCTCAAGGACGCGGTCGCCGTGCGGGACGTGTCCACGCCGCTGACGCAGGTGCGCTACACGGG
CACCTACAACGGCGGGTTCCCCGGCTGGCAGCCGTTCGTGGACGGCGGGGAGACCGTGGAGGTGGAGATCAACAA
GAACGGCCCGGTGCTGCCGGGGCTCTCCAACTTCTATCTGGCCGGGGTGTGGGTCACCGTCGGCGGGCTGATCCG
GGCGGTGGCCTCGGGCCGGCAGGTCACGCAGGTGATCTGCCGGGACGACGGGCGGGAGTTCACGGCGAGCGTGGA
CGAGAGCGCGCCGCCGCCCACCCAGGTCGCCATCCCGGTGGGCAAGCAGCCGGGCGTGCCGGATCTGGCGGCCGG
GTTCCCCGCCCAGACCGCCGGCGGCGAGACCGCCACCGGCGCCAACAACACCGTCACATCGTCGAGGAGCGTGTA
GTGCAGGTACACACATGGGTGATCGCCGAGCACATCCCGGGGGTCCCGGACACGGACCGGATCTACCGGAAGGTG
ACGCGGGAGTTCGATCCCGCCTCGCTGGCGGACGATCAGATGCTGCTGCGCACCCGGTACGTGTCGGTGGACCCG
TATCTGGTGGGGCTCTCGCTCCAGACGCCGATCGGGGACACCGTGCGCGGTGACTCGATCATGGAGGTGGCCGTG
GCGGGGCCGCGCGCCCGCTTCCAGGTCGGGGACCTGGTGCAGGGGTACGGCGGCTGGTGCAGCCATCTGGTGTCC
ACCGGGGGGCCCAGCGGATGGAACGACGACGGCGCCGAGTTCGCCGTCCAGTTGCCGCCGTTCCGCAAGCTGGAC
CCGCGGCGGTACGACGAGGCGCTGCCGCTGTCCACGGCGCTGGGCGTGATGGGCACCCCGGGCATCACCGCGTTC
GGCGCGATGAAGACGTTCCTGACCGTGGGCTCCGAGGACACGGTGGTGATCAGCGGGGCGTCCGGGACGGTGGGC
ACCCTGGTGGGCCAGCTCGCCAAGCGGGCCGGGGCCCGGGTGGTGGGCACCACCTCCTCGCCGGGGAAGGCCGCG
TATCTGACGCAGCTGGGCTTCGACGCGGTGGTGAACTACCGGCAGGGCGACGACACGGACACGGTGCGCGAGGCG
CTGGCGGCGGCAGCGCCCAACGGAATCGACAAGTACTTCGACAACCTGGGCGGCACCGTGACGGACGCGGTGTTC
ACGATGCTCAACGTGCACTCCCAGGTGGCGGTGTGCTGGCAGTGGGCCACCACGGTCAACGGGGACTGGACGGGG
CCGCGGCTGCTGCCGTACATCATGTTCCCGCGCACCACGATCCGGGGGATCTTCGCCGACGAGTGGTACACGGAG
GAGATGGTCGACGCGCTGCACGAGGAGGTGGGCGGGCTGATCCGCAAGGGTGAGCTGGCCTACCACCAGACCATC
CACCAGGGCTTCGACGCCCTCCCGGACGCGTACCGCTCCCTGTACACCGGCCAGGAGGGCAACCGCGGCAAGGTC
CTGGTCGCCCTGTAG

#### 7.4 pWHM1120::*ika*

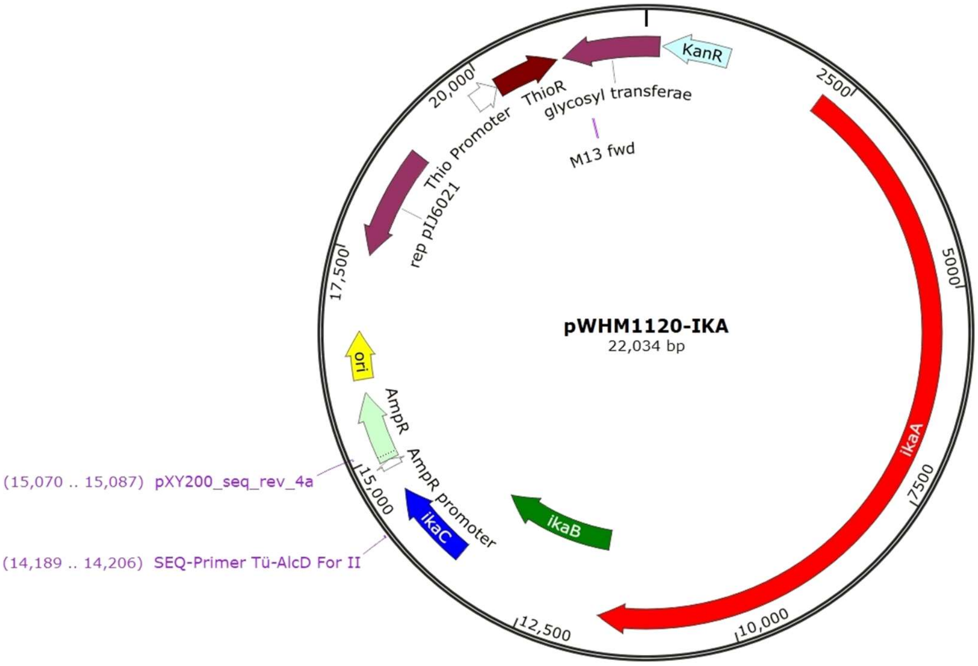

**Figure 5.** Vector map of *ika* in pWHM1120.

- DNA sequence of pWHM1120::*ika* [22034 bp]:

GGCAGTCCCGGAAAGCCCAGGGTCAGCACGAGCTGGTCGACGCCCGCCCTGGCCAGGCTGCGGGACAACGCCAGG
ATCTGCACCTGCATACCGCCCACCGGATCGAACTCGGCCGGCCAGTGGTCGACGTAGTCGTGGTGGAAAAAGGGA
GTCAGCCGAAGTACACGCACGACGCCATCCGATTCTTCCGGAAGAGTGCCGAAAAGCTATTTCTGAATGACGTAA
ATCAGCTCGTTGCCGATCTCCAGTCGCTGAATGCGGCACGACCGCTCTCTGGCCTGGGACTCAAAACTTTGTGAA
TAGTTCTGAAACATCCCCTTCGATCGCTGACCGAGAGACTTGGTCGGGAAGGTTACCACGATAATCGGCGAGTTG
ACAATGTCAATCACTTCCCAGCCGGAGCCTCGTCGCTGAGTCTCCAGACAGGGGAGCGTCTTCAGCAATAGCGTG
ACGTCGGTCGGTTCGTCAAGGCGGTCCTCAAGGACGTCGACCACACTCGTGCGGTGCGCGACGCCCAACCTCGTC
AGGGCGGCGTCCACGAAGTCGATCAGCCGGGCGTCGATGTCGGAGGCGACGTAGACGGTCTCGTCCGACAGGCCC
ATCCACGGGGCGGCCAGCGGGTTGAGGCCGCAGGCAAGGTCACGCAGCGTGTTGGGCCGGGGGACGTGGCGGAAG
ACCTCCCGGTAGAACTCCGCCAGGTGGGGCAATCGTTCGCGGGTGGACACGTGCACCGACATCGCGCGCCGCAGG
GCCGCCTGGACCGCCGCGTCGTCACCGTCGTCCACAGCGGAGTCGAGCTGCCGGAGCAACGCTGCGTAGTTGGGC
GGACTGGGCGGCAGGAAGGCGCCGTAGATCTCGTGCAGCCCGCGCTTGGTGCGCTTCACCGCGTCCGGCACGTCG
CCCCGCGTGGCGACGAGGGCGGCGCGGGCCAGCCGCCGCACGGTGGCCGGGGCCACCGTCTGGTAGCGCCGGCTC
TTGGTGATGGCCTGCTCGACCTGCTCGATACGGTCGTCGGGCACAGATGTCGTCATCGAATCCTCCGAAAGATCA
TGAACCGGCAGCGAAGCGGAAAAATTCCCCCACCAATACCCGCTCCGCACACCAAGAGTCATTGCCACGTGTACT
TAAAGTTATTGCAGGAGTATTACCGGGCGGGCATGGGAGCGCTCGCAGCCCGCCCCTCGTCACGTCCATATTGGT
CGGCCTGTCCTGCAAGCCCGGCTTGGCCGGTTTTTGCACGTCGACACTGGTGATTGTGTCTGGCACACTTGGCCG
GGGCCGCGATCCTCGCCGAGCAGGCCGTCGAGGACATCTTCCCCCTCCCCCGCGAGCTGGTGGGCGAGGGCGAGG
TCTTCATGCTCCAGGTCAAGGGCGACTCGATGCTCGACGCCGCCATCTGCGACGGGGACTGGGTCGTGGTCCGGC
AGCAGCCCACCGCCGACAGCGGCGAGATCTACAAGGGCCGCCGGATACAGGGCCGGACGTCACCGACGGCGGGGG
CCACCACGGCGGCGGTCACGGCGGTGACGGTCACGGCGGCGGCCATCACGGCGGCGGTTACGCCGTGTTCGTGGA
CGGCGTCGAACTGCATGTGATGCGCAACGCCGACGGCTCGTGGATCAGCGTCGTCAGCCACTACGAGCCGGTGGA
CACCCCGCGCGCCGCGGCCCGCGCTGCGGTCGACGAGCTCGGTACCCGGGGATCTGTGTTGGCGCACAAATCAAC
GGGGATTACTGTCGTTTAATGTGATTTAAACTGTGAAATAGTATGGTTTTCAGTTATTGAAACGCCGTGAGCGGG
GAAAACTTGCTTTTTCCCGTTTCCGGGGTTGGACAACTGAGCAACGCGAAGGCGTCAGCTACGATGTTCCGGGGA
CTGCTGATCCGGTCAGCAGGTGGAAGAGGGACTGGATTCCAAAGTTCTCAATGCTGCTTGCTGTTCTTGAATGGG
GGGTCGTTGACGACGACATGGCTCGATTGGCGCGACAAGTTGCTGCGATTCTCACCAATAAAAAACGCCCGGCGG

CAACCGAGCGTTCTGAACAAATCCAGATGGAGTTCTGAGGTCATTACTGGACCGGATCGGGGATCTGGGCTGAGG
GAGCCGACGGCACGCGGCGGCTCACGGCGTGGCACGCGGAACGTCCGGGCTTGCACCTCACGTCACGTGAGGAGG
CAGCGTGGACGGCGTCAGAGAAGGGAGCGGACATATGAAGCTTGCATGCCTGCAGGTCGACTCTAGAATGGATTC
CATGCACCACCCTGCCCCCGTCCCCGTACCCGAAGTCCCCGCGCCCGTCCCGTCCCAGGACGACGCGTTCGCCAT
CGTCGGCATCGGCTGCCGGCTGCCCGGCGGCGCCAGCGACTACCGGACCTTCTGGCGCAACCTCCTCGACGGCAA
GGACTGCATCACCGACACCCCCGCCGACCGCTACGACACCCGCACCCTGGGCAGCGGCGACAAGGCCAAGCCCGG
CCGGCTGGTCGGCGGACGCGGTGGATACATCGACGGCTTCGACGAGTTCGACCCCGCCTTCTTCGGCATCAGCCC
GCGCGAGGCCGAGCACATGGACCCCCAGCAGCGGAAGCTCCTGGAGGTCGCCTGGGAGGCGCTGGAGGACGGCGG
CCTCAAGCCCGCCGAGCTGGCCGGCAGCGATGTCGGGGTGTACGTCGGGGCGTTCACCCTCGACTACAAGATCCT
GCAGTTCGCCGACCTCGGCTTCGAGACCCTGGCCGCGCACACCGCCACCGGCACCATGATGACGATGGTGTCCAA
CCGGATCTCGTACTGCTTCGACTTCCGCGGACCCTCGGTCTCCGTCGACACCGCGTGCAGCGGCTCCCTGGTCGC
CGTCCACCTCGCCTGCCAGAGCCTGCGCCGCGGCGAGACCTCCGTCGCCCTGGCCGGCGGCACCCTGCTGCACAT
GGCGCCGCAGTACACCATCGCCGAGACCAAGGGCGGGTTCCTCTCCCCCGACGGCCGCTCCCGCGCCCTGGACGC
CTCCGCCAACGGCTACGTGCGCGCCGAGGGCGTCGGCATGGTCGCCATCAAGCGCCTCGCGGACGCGCAGCGCGA
CGGCGATCCCATCCACGCCGTCATCATCGGCAGCGGCGTCAACCAGGACGGCCGCACCAACGGCATCACCGTGCC
CAACCCCGACGCGCAGGTCGCCCTGATCGAGCGGGTCTGCGCCGCCGCCGGCGTCACCCCCGGCAGCCTCCAGTA
CGTCGAGGCGCACGGCACCTCCACCCCCGTCGGCGACCCGCTGGAGGCCAACGCCCTCGGCCGCGCGCTCTCCAT
CGGCCGCGAGCCGGGCGCCCGGACGTACGTCGGCTCGGTCAAGACCAACATCGGGCACACCGAGTCCGCCGCCGG
CATCGCCGGGCTGATCAAGACGGTGCTCAGCCTCAAGCACAAGGTCATCCCGCCGCACATCAACCTGGAGAAGCT
CAACCCGCAGATCGACGAGGCGTCCCTGCCGTACGAGATCCCCCGCGAGCCCACCCCCTGGCCCGAGCACAGCGG
GCCGGCCCGGGCCGGCGTCAACTCCTTCGGCTTCGGCGGACCAACGCCCACGTCCTGCTCCAGGAGGCACCGCC
GACCGTCGGGGAGCCCGCGCCACCGGCCACCGACGGGTACTCCGTGCTGCCGCTCAGCGCCCGCGACCCCGAAGC
CTTTCCCGCCATCGCCACCGGCCTGCGCGAACGGCTCGCCGAGGGACTGCCGGTGGGCGACGCCGCCTACACCCT
CGCCCACCGGCGGCAGCATCTGGAGCAGCGGCTGTCCGTCGTGTACGACTCCCCCGAGGCCCTCGACGAGGTGCT
CGGCGCCGTCGCCCGCGGCGAGAGCCACCCGCGTGCCGTCGCCGGCACCCAGCGGGAGGGCCTGGACCGCAGGCT
GGTGTGGGTGTTCACCGGCATGGGCCCGCAGTGGTGGGCCATGGGCCGCCAGTTGTACGCGAGCGAGCCCGTCTA
CCGGGAGGTCATCGACCGCTGCGACCAGGAGATCGCCGCGCTCACCGGCTGGTCCCTCACCCAGGAGCTGAACGC
CGACGAGGCCGACTCCCGGATGAGCGAGACCTGGCTCGCCCAGCCCGCCAACTTCGCCGTCCAGATCGCCCTGGC
CGCCCTGTGGCGCAGCAAGGGGATCCAGCCCGACGCCGTCACCGGGCACAGCACCGGTGAGGTCGCCGCGTTCTA
CGAGGCCGGGGTGTACACCCTCCCCGAGGCCGTGAAGATCGTGGTGCACCGCAGCCGGCTCCAGCAGAAGCTCAT
CGGCACCGGCTCCATGCTCGCCGTCAGCCTCACCGAGGCGGAGGCCGCCCGCCGGGTGCGCCCGCACGGCGACCG
GGTCTCCATCGCCGCCGTCAACAGCCCCACCTCCATCACCCTGGCCGGGGACACCGAGGCGCTGGAGGTGATCGC
CGCCGAGCTGGGCGCCGAGGACATCTTCGCCCGCTTCCTGGAGGTCGGCGTCCCGTACCACAGCCCCCGCATGGA
GCTGATCAAGGACGAGCTGCTGACCTCGCTCGCCGATCTCAAGCCGCAGCAGGCGAAGTTGCCGCTGTACCTCAC
CGCGCTGCCGGGCACCGTCGCGCAGGGCACGGAGCTGGACGCCGACTACTGGTGGCGCAATGTGCGCGAGGCCGT
GCACTTCCGGGCCGCCGTGGACCGGCTGCTGGACGACGGCTACGGCGTCTTCCTGGAGATCGGCCCGCACCCCGT
GCTCGCCCACTCCCTGCGCGAGTGCTGCGAGGCCCGCGACGCGCACAGCGTCACCCTGGCCTCCATCCGCCGCAA
GGCGGACGAGCGCGAACGCCTCACCCTGTCGCTCGCCGCGCTGCACAGCCTCGGCTTCGCCGTGGACTGGCACGC
CCTGCACCCCGCCGGGCGGCCGGCCGAACTGCCGCGCTACCCGTTCCGGCGCGACCGGTACTGGGTCGAGCCGGC
CCCGGTCGCGCAGATCCGGCTCGGCCACCGCGACCACCCGCTGCTGGGCCGCCGCACCGCGAGCGCCGAGCCGGT
GTGGGAGGTGAAGCTGGACGCGGAGGCCGCCCCGTACCTGGAGGACCACCGCATCCAGGGCACCGTGCTGTTCCC
GGCCGCCGGCTATCTGGAGATGGCCGCGCAGGCCATGCGGGCGCTGACCGGTGATGAGCACAGCACCGCCGCGCT
GGCCGGCATCGAGCTGCGCAAGGCGCTGTTCCTGCCGGACGGCGAGCCGCAGACGGTGCAGCTGTCCTTCTCCTC
CGACGCCGCCGCGTTCTCCATCGCCACCGTGGGCGCCGCCGGCGCCGAGCCGACCGTGCACGCCACCGGTACGGT
ACGGGCCGCCCAGCGCCGCCGGCTGACCGCGCCGCTGGACACCGTCGCCGTCCGGGCCCGCGCCGCCCGCCACCT
GAGCGGCCCCGACTGCTACGCCGAACTGGCCGCGCTCGGCTACCACTACGGCCCCGCCTTCCAGGGCATCGAGGA
GGTGTGGATCGGCGAGGGCGAGGCCCTGGCCCGGATCCGTCCGCCGCAGGGGCTCACCCCGGACGCGGCGGCGCA
CCACATGCATCCGGTGCTGCTCGACTCCTGCTTCCAGTCGCTGCTGACCCCGCAGCTGCTCACCGCGCCCGCCGG
GCCCGGGGGCACCGGCATCCGGCTGCCGCTGTCCATCGCCGAGGTACGGCTGGACCCGGTCGGCGACCGCGAACT
GTGGGTGCACGCCACCGTCACCGGCGACGACGAGGACGAACTCACCGGTGACATCGCCGTGTACGACGGCGCCGA
CGGTACGCCGCTGGGCCGCGTCGCCGGCTTCCGCGCCGCCGATGTGGAGAAGGCCGCCACCACCGTGGGGCTGTC
CACCATCGACAGCTGGCTCACCGAACCGAGCTGGGTGCCGTGCCCGCTGCCCGAGGCGGCGTCCGCCGCGCCGGC
GGCCGGGCGGCACGTACTGTTCGCCGACGCGGGCGGGGTCGCGCAGCGGCTGGCCGCGCTGATCGGCGAGGCCGG
CGGGGAGGCCCATCTGGTCCGGCCCGGTGCCGCGTACGGCCTGGACCGCACGGCGAGGACCGCCACCGTCGTCCC

CGGATCCGCGGATGACCTGCGGCGGTTGCTCACCGATCTCGGGCAGGTGGACGGCGTCGTCCACCTGTGGAACCT
GGACCGGCCGGCGCTGGCCGACGCCCCGCGCGGACGGTTCGCGGACATCGCCTCCACCGGCGCGTACGCCCTGAT
CGCCCTCACTCAGGCCCTGCTCGCCGACCCGGAGCGGCACGGCGGCACCCCGGTGCACATCGTCACCAGAGCCGC
CCAGTGCGTGGTCCCCGGTGAGCCGGTGGAGCCGCTGGGCGCGCCCGCCTGGGGCATCGGCCGGGTGCTGTGGCA
GCAGGAACTGGCCGGGCGCGGCGGCAAGCTGATCGACCTGGCGGCCGACGGCGGCGTCGAGGAGGACGCGTACGC
GCTGCTGCGCGAGCTGGCCGACCCCACCGGCGCGGCCGAGCGCGAGGACGAGATCGCGCTGCGCGCCGGGGAGCG
GCACACCAGCCGGCTGGTGGCCGCCGAGGGGCTGAGCAGGCCGCTGCCCCTGCGGCTGCGCCCGGACGGCAGCTA
TCTGGTGACCGGCGCGTTCGGCGCGCTCGGCAGGCTGCTGTGCCGCACGCTGGTCAGGCGCGGGGCGCGGCGGCT
GATCCTGGTGGGCCGCACCCGGCTGCCGGAGCGCGAGCGCTGGGCCGACCAGGACCCGAACTCGCCGGCCGGGCG
GCACGTGGCCTTCCTCAAGGAGCTGGAGGCGCTGGGCGCGCAGCCGATTCTCGCGCCGCTGGACATCACCGACGA
GGACGCGCTGGCCGGCTGGCTCGCCGGGTACCGGCGCGCCCAGGGGCCGCCGATCCGCGGGGTGTTCCATCTGGC
GGGGCAGGTGCGCGACACCCTGGTGCCGGAGATGGACCGGGAGGTGTTCGACGCCGTCCACGACCCGAAGGTGGT
GGGCGCGGCGCTGCTGCACCGGCAGCTGAGCGGCGAACCGCTGGAGCACTTCGTGCTGTTCGCCTCGGTCGCGGC
CTGGCTGACGACGGCCGGACAGACCAACTACGCGGCGGGGAACGCCTTCCTGGACGCGCTGGCGCACCACCGCCG
CGCGCAGGGGCTGCCGGCGCTGGCGCTGGACTGGGGCCCGTGGGCCACCGGCATGATCGAGGAACTGGGCCTGAT
CGACCACTACCGCAACAGCCGGGGCATGTCCTCGCTGGCGCCCGAGGCGGGCATGGCGGTGCTGGAGCGGGTCAT
CGGGCAGGACCGGGCACAGCTGCTGGTGGCCACGGTCGTGGACTGGCCGGTGTTCATGTCCTGGTACGCGGCGCC
GCCGCGGCTGGTCACGGAGCTGGCGGCCACCGCCCAGGGACCGGGGTCCGAGGGCGACGGCAGTTTCCTGGACGC
GTTCCGGGAGGCCACCGCGGACAAGCGGCGGCTGCTGCTGACCGAGCGGTTCACGACGCTGGTGGCGGGTGTGCT
GCGGGTGCGGGCCGAGCAGGTGGATCCGGCGGTCAGCCTGAATCTGCTGGGGCTCGACTCGCTGCTGGCGATGGA
GCTGCGAGCGCGGGTGGTGGCCGAGGTGGGCATCGCGCTGCCGGTGGTGGCGCTGCTGTCCAGCGCGCCGGCCGG
GGACCTGATCACCCAGCTGCACGAGGGCCTGGAGGAGTTGCTGGCCGAGGAGGGCAGCGGCGCCGCGGTGACGGC
GGTGGAGCGCTTCGAGGACGAGGCCGAGTTCCCGCTGACGCAGAACCAGAAGGCGCTGTGGTTCCTGAAGCAGCT
GAACCCGGACGGCTTCGCGTACAACATCGGCGGCGCCGTCGAGGTGCGGGTCGAGCTGGACCCGGACCTGATGTT
CGAGGCGTTTCGCCGGCTGCTGGCCCGGCATCCCGTGCTGCGGGCGAACTTCCTGCTGGTGGAGGGGCAGGCGGT
GCAGCGGATCTCCCCGGAGATCAAGGAGGACATCGCGCTCTTCGACGTCGAGGACCGCGCGTGGGACGACATCTA
CCGGATGATCATCGAGGAGTACCGCAAGCCGTACGACCTGGCGACCGATCCGCTGATCCGGTTCCGCCTCTTCCG
GCGCGGCCCGGACCGCTGGGTCATCACCAAGGCCGTCCACCACATCATCTCGGACGCCATCTCCACCTTCACCTT
CATCGAGGAACTGCTGTCCCTGTACGAGGGGCTGCGGCAGGGCCACGACGTCGAACTGCCGCCGGTGTCCGCCCG
CTATCTGGACTTCCTCAACTGGCAGAACGCGTTCCTGGCCGGCCGCGAGGCGCAGAAGATGCTCGCGTACTGGCG
GGGGCAGCTGCCGGACGAGGTGCCGGTGCTGGCGCTGCCCACCGACAAGCCGCGCCCGGCGGTGCTCACCCACAA
CGGGGCGTCCGAGTTCTTCGCCCTGGACGCGGAGTTGAGCGCCCGGGTGCACGCGCTGGCGCGGGAGCACAACGT
CACCGTCTTCATGGTGCTGCTGAGCGCGTACTACCTGCTGCTGCACCGCTATGCGGGGCAGGACGACATCATCGT
CGGCTCCCCCGTCACCGGCCGCACCCAGGAGGAGTTCGGCGCCGTCTACGGGTACTTCGTGAACCCGCTGCCGCT
GCACGCCTCGCTGGCCGGTGACCCCACGGTCGCCGAGCTGCTGGACCAGGTGCGCACCACGGTGCTGGGCGGCCT
GGACCACCAGGAGTACCCGTTCACGCTGCTGGTGGAGCAGCTGGGGCTGGCCCACGACCCGAGCCGGTCGGCGGT
CTTCCAGGCGATGTTCATCCTGCTGCACCACAAGGTGGCCACCGAGAAGTACGGCTACAAGCTGGAGTACATCGA
GCTGCCCGAGGAGGAGGGCCAGTTCGACCTGACGCTGTCCGCGTACGAGGAGGAGGCGGACGGGCGGTTCCACTG
CGTCTTCAAGTACAACACCGACCTCTTCGAGGCGGAGACGATCCGGCGGCTCGCCGGGCACTACACGCAGCTCCT
GGAGTCGCTGACCGCGGCGCCCGCCGACGCCGCCACCGGTGGACTGCGGATGCTGTCGGGCGGCGAGCGGGAGCG
GATCCTCACCGAGTGGAGCGGGGCCGGGCAGGGCGCGCAGGACGCGCCGGTGCCGGTGCACCGGCTGATCGCCGA
GGCGGCGCACCGTACCCCGCAGGCGATCGCGGTGGCCGCGCCCGCCGAGAGCGGGGAGACCCGGCGGCTGACGTA
CGGCGAACTGGAGGAGCGCGCCGGCGAACTGGCCGGGCGGCTGCGGGCGCGCGGCGTGCGCGAGGGCACCGTCGT
CGCGCTGTGCCTGGAGAAGTCGCCCGAGCTGATCACCGCCCTGCTGGCGGTCCTCAAGGCGGGCGGCGCCTATCT
GCCGCTGGACCCGGACTATCCGGCCGACCGGCTCGCGTACATGGTGCGCAACGCCGGGGCCACGCTGGTGATCGG
CGGGACGGGCGGCGCGGCCGAGGGGCTGCCGGGCACCGTGGTCACCCTGGAGGAACTGCTCGCGGGCGAGGCCGG
CGAAGCGGGGCCGGACGCCGAGCCGGGGCCCGACTCCCCCGCCTACGTCATCTACACCTCGGGCTCCACCGGGCG
CCCCAAGGCGGTCGCGGTCAGCCACCGCAATCTGGCCTCGGTGTACGCCGGATGGCGCGACGCCTACCGCCTGGA
GGAGGGCGGCATCCGGGTCCATCTCCAGATGGCCAGCCCCTCCTTCGACGTCTTCACCGGCGACCTGACCCGAGC
CCTGTGCTCGGGCGGCACGCTGGTGCTGGTCGGCCGGGAGCTGCTGTTCAACACCGCCCGGCTGTACGAGACGAT
GCGCGCCGAACGGGTGGACTGCGGCGAGTTCGTGCCCGCCGTGGTGCGCACCCTGGTGCGGCACTGCGAGGACAC
CGGCGCCCGGCTGGACTTCCTGCGGCTGCTGATCGTGGGCTCGGACTCCTGGAAGGCCGAGGAGTACGAGCGGCT
GCGCGCGCTGGGCGCACAGCGCCTGGTGAACTCGTACGGGCTCACCGAGGCCACCATCGACAGCGCCTGGTTCGA
GGGTCCCGCGGATGACCTGGAGGGCGGCCGGATGGTGCCCATCGGGCGGCCGTTCCCGGGCAGCGCGCTGTACAT

CCTGGACTCGCGCGGCGAGCCGGTGCCGCCCGGTGTCCCCGGCGAGCTGTGGATCGGCGGCACCGGGGTGGCGCT
CGGCTACCTCGGCGACGAGGCGCTGACCGGGGAGCGGTTCCTCACCCGCGCCCTGGCCGGCGACGCTCCGGTACG
GCTGTACCGCACCGGTGACCTCGCGCGCTGGGACGCGGCCGGCACCGTCCATCTGCTGGGCCGGGCCGACTCGCA
GATCAAGGTGCGCGGGCACCGCATCGAGATCGGGGAGATCGAGTCGCACCTGGCGGCCTGCCCCGAGCTGGCCCA
GGCGCAGGTCACCGTGCGGCCGGACGCGGGCGGCGAGAACGTGCTGTGCGCGTACGGGGTGGCGGCCCCGGGCGC
CGTGCTGGACTGGCGCGAGGTGCGCCGGCGCCTGGCGGACTATCTGCCGACGTTCATGATCCCCACCCACTTCAC
CGAGCTGCCCGCCCTGCCGCTCACCCCGAACGGCAAGGTGGACGTGGCGGCGCTGCCCGCCCCGCGCACCGGCGA
CGGCGCGGACGGGCCGGTGTACGAGGCCCCCGTCACGCTGTACGAGACCCGGATGGCCGAGCACTGGCAGCGGCT
GCTGGGCATCGAGGCCCCCGGGCCCGGTCTGGGCCACGACTTCTTCGAGACCGGTGGCAGCTCCATCCGGCTGAT
CGAGCTGATCTACCACCTGCAGGCCGAGTTCGGGATCTCCATCCCGGTCAGCCGGCTGTTCCAGGTGACGACGCT
GCACGGCATGGCCAAGACGGTCGAGCGGATCGTCACCGGGGAGATCGAGGGGTCGCTGCCGTATCTGCGGTTCAA
CGAGAACGCCGCGGCGGGCACGGTGTTCTGCTTCCCGCCGGCCGGTGGCCACGGCCTGGTCTACCGGGAGTTCGC
GGCGCGGCTGCCGGAGTTCGAGTTCCTCGCCTTCAACTACCTGATGGGCGAGGACAAGGTAAGCGGGTACGCCGA
CCTGGTGGCCGGGCACCGGCCGGAGGGCGAGATCGACCTGCTCGGCTACTCGCTGGGCGGCAACCTCGCCTTCGA
GGTGGCCAAGGAGCTGGAGCGGCGCGGCCGCACCGTGCGCCACGTCGTCATCATGGACTCGCTGCGGGTGACGGA
GTCCTACGAGCTGGGCCCGGAGCACCTGGCCGTCTTCGAGCGCGAGCTGGCCGAGCATCTGCGCAAGCACACCGG
CTCGGCGCTGGTCGCGGAGAAGACGCGCGAACAGGCCAAGGACTACCTGGAGTTCACCGGCCGCACCGCCAACCC
CGGCACCACCGGGGCCCGGATCGCGGTGATCAGTGACGAGGAGAACGCGGCCGCGTACGACAGCGGCGCCGAGGG
CAGCTGGCACGGCGCCTCCCGTACCGGAACCGACGTGCTGCGCGGGGTGGGCCGGCACGCCGACATGCTCGATCC
GGGGACGGTCGAGCACAACGCGCGCCTGGCGCGCGGCATTCTCACCGGCGGTGATGGCGAGGTATGACGCCTTTC
GTTCAGCCGGCGGTCGACACCAAGGAGCACAGCGCCATGTCATCCCCCACCACCTCCGGCACCCCGGGCAGGCAG
TCGATGATCATCATCGGCGGCGGCCTGGGGGGCCTGTCCACCGGCTGCTACGCGCAGATGAACGGCTACGCGACG
CGGGTCTTCGAGATGCACGAGATCCCGGGCGGTTCCTGCACCGCCTGGGAGCGCGGGGACTTCACCTTCGACTGG
TGCGTCAGCTGGCTGCTGGGCAGCGGTCCCGGCAACGAGATGTACCAGATCTGGATGGAACTGGGGGCGTTGCAG
GGCAAGGAGATGCGCCAGTTCGACGTCTTCAACATCGTGCGGGTGCGCGGCGGCCAGCCGGTGTACTTCTACTCC
GACCCGGACCGGCTCCAGGCGCACCTGCTGGAGATCTCCCCGGCCGACGCCCGCCGCATCAAGAACTTCTGCGAG
GGGGTGCGCACCTTCCAGAAGGCGCTGTCGGTCTACCCGTTCCTCAAGCCGGTGGGGCTGATGGGGCGGTGGGAA
CGGTGGAAGATGCTGGCCTCGTTCCTGCCGTACTTCAACGCCATCCGCAAGTCCATCACCGAGCTGATGACGGAC
TACGCGGAGAAGTTCCAGCACCCGGTGCTGCGCGAGGCCTTCAACTACGTGCTGTACGAGAAGCACGCCGACTTC
CCCGTCCTGCCGTTCTGGTTCCAGCTGGCCTCGCACGCCAACGGCTCGGCGGGGGTGCCCGAGGGCGGCTCGCTG
GAGCTGGCCCGGTCCGTGGAGCGGCGCTACCTGGGGCTCGGCGGGGAGATCACCTACAACGCCAAGGTGGAGAAG
ATCCTCGTCGAGCACGACAAGGCGGTGGGAGTGCGGCTCACCGACGGCCGCGAGTTCCGCGCGGACATCGTGGTG
TCGGCGGCCGATCTGCACACCACCGCCATGGAGATGCTCGGCGGCCGGTATCTCAACGACACCTGGCGCAAGCTG
CTCACCGAGACGATCGACGAGGTGGGCACGATCTCCCCCGGCTATGTCTCGCTGTTCCTGGGGCTGCGCCGGCCG
TTCCCCGAGGGCGAGCCGTGCACCACGTACGTGCTGGAGGACAGCATGGCGGAGAAGCTCACCGGCATGCGGCAT
CCCAGCATGAACGTGCAGTTCCGCAGCTGCCACTACCCGGAGCTGTCGCCGCGCGAGACCACGGTCATCTTCGCC
ACGTACTTCTCGGAGGCCGAGCCGTGGCGGGCGCTGCGCGACGACGTGCCGGAACAGGCGGGCCGGGTGCGGCGC
GGTCAGGTGCTGCACACCCTGCCGGTGAAGCACGGCAAGGCGTACACCCAGGCCAAGCGGCAGGCGCGGATCACC
ATCGAGAACTTCCTGGACGAGCGGTTCCCCGGTCTCAAGGACGCGGTCGCCGTGCGGGACGTGTCCACGCCGCTG
ACGCAGGTGCGCTACACGGGCACCTACAACGGCGGGTTCCCCGGCTGGCAGCCGTTCGTGGACGGCGGGGAGACC
GTGGAGGTGGAGATCAACAAGAACGGCCCGGTGCTGCCGGGGCTCTCCAACTTCTATCTGGCCGGGGTGTGGGTC
ACCGTCGGCGGGCTGATCCGGGCGGTGGCCTCGGGCCGGCAGGTCACGCAGGTGATCTGCCGGGACGACGGGCGG
GAGTTCACGGCGAGCGTGGACGAGAGCGCGCCGCCGCCCACCCAGGTCGCCATCCCGGTGGGCAAGCAGCCGGGC
GTGCCGGATCTGGCGGCCGGGTTCCCCGCCCAGACCGCCGGCGGCGAGACCGCCACCGGCGCCAACAACACCGTC
ACATCGTCGAGGAGCGTGTAGTGCAGGTACACACATGGGTGATCGCCGAGCACATCCCGGGGGTCCCGGACACGG
ACCGGATCTACCGGAAGGTGACGCGGGAGTTCGATCCCGCCTCGCTGGCGGACGATCAGATGCTGCTGCGCACCC
GGTACGTGTCGGTGGACCCGTATCTGGTGGGGCTCTCGCTCCAGACGCCGATCGGGGACACCGTGCGCGGTGACT
CGATCATGGAGGTGGCCGTGGCGGGGCCGCGCGCCCGCTTCCAGGTCGGGGACCTGGTGCAGGGGTACGGCGGCT
GGTGCAGCCATCTGGTGTCCACCGGGGGGCCCAGCGGATGGAACGACGACGGCGCCGAGTTCGCCGTCCAGTTGC
CGCCGTTCCGCAAGCTGGACCCGCGGCGGTACGACGAGGCGCTGCCGCTGTCCACGGCGCTGGGCGTGATGGGCA
CCCCGGGCATCACCGCGTTCGGCGCGATGAAGACGTTCCTGACCGTGGGCTCCGAGGACACGGTGGTGATCAGCG
GGGCGTCCGGGACGGTGGGCACCCTGGTGGGCCAGCTCGCCAAGCGGGCCGGGGCCCGGGTGGTGGGCACCACCT
CCTCGCCGGGGAAGGCCGCGTATCTGACGCAGCTGGGCTTCGACGCGGTGGTGAACTACCGGCAGGGCGACGACA
CGGACACGGTGCGCGAGGCGCTGGCGGCGGCAGCGCCCAACGGAATCGACAAGTACTTCGACAACCTGGGCGGCA

CCGTGACGGACGCGGTGTTCACGATGCTCAACGTGCACTCCCAGGTGGCGGTGTGCTGGCAGTGGGCCACCACGG
TCAACGGGGACTGGACGGGGCCGCGGCTGCTGCCGTACATCATGTTCCCGCGCACCACGATCCGGGGGATCTTCG
CCGACGAGTGGTACACGGAGGAGATGGTCGACGCGCTGCACGAGGAGGTGGGCGGGCTGATCCGCAAGGGTGAGC
TGGCCTACCACCAGACCATCCACCAGGGCTTCGACGCCCTCCCGGACGCGTACCGCTCCCTGTACACCGGCCAGG
AGGGCAACCGCGGCAAGGTCCTGGTCGCCCTGTAGAGGATCCCCGGGTACCGAGCTCGAATTTATGGTGCACTCT
CAGTACAATCTGCTCTGATGCCGCATAGTTAAGCCAGCCCCGACACCCGCCAACACCCGCTGACGCGCCCTGACG
GGCTTGTCTGCTCCCGGCATCCGCTTACAGACAAGCTGTGACCGTCTCCGGGAGCTGCATGTGTCAGAGGTTTTC
ACCGTCATCACCGAAACGCGCGAGACGAAAGGGCCTCGTGATACGCCTATTTTTATAGGTTAATGTCATGATAAT
AATGGTTTCTTAGACGTCAGGTGGCACTTTTCGGGGAAATGTGCGCGGAACCCCTATTTGTTTATTTTTCTAAAT
ACATTCAAATATGTATCCGCTCATGAGACAATAACCCTGATAAATGCTTCAATAATATTGAAAAAGGAAGAGTAT
GAGTATTCAACATTTCCGTGTCGCCCTTATTCCCTTTTTTGCGGCATTTTGCCTTCCTGTTTTTGCTCACCCAGA
AACGCTGGTGAAAGTAAAAGATGCTGAAGATCAGTTGGGTGCACGAGTGGGTTACATCGAACTGGATCTCAACAG
CGGTAAGATCCTTGAGAGTTTTCGCCCCGAAGAACGTTTTCCAATGATGAGCACTTTTAAAGTTCTGCTATGTGG
CGCGGTATTATCCCGTATTGACGCCGGGCAAGAGCAACTCGGTCGCCGCATACACTATTCTCAGAATGACTTGGT
TGAGTACTCACCAGTCACAGAAAAGCATCTTACGGATGGCATGACAGTAAGAGAATTATGCAGTGCTGCCATAAC
CATGAGTGATAACACTGCGGCCAACTTACTTCTGACAACGATCGGAGGACCGAAGGAGCTAACCGCTTTTTTGCA
CAACATGGGGGATCATGTAACTCGCCTTGATCGTTGGGAACCGGAGCTGAATGAAGCCATACCAAACGACGAGCG
TGACACCACGATGCCTGTAGCAATGGCAACAACGTTGCGCAAACTATTAACTGGCGAACTACTTACTCTAGCTTC
CCGGCAACAATTAATAGACTGGATGGAGGCGGATAAAGTTGCAGGACCACTTCTGCGCTCGGCCCTTCCGGCTGG
CTGGTTTATTGCTGATAAATCTGGAGCCGGTGAGCGTGGGTCTCGCGGTATCATTGCAGCACTGGGGCCAGATGG
TAAGCCCTCCCGTATCGTAGTTATCTACACGACGGGGAGTCAGGCAACTATGGATGAACGAAATAGACAGATCGC
TGAGATAGGTGCCTCACTGATTAAGCATTGGTAACTGTCAGACCAAGTTTACTCATATATACTTTAGATTGATTT
AAAACTTCATTTTTAATTTAAAAGGATCTAGGTGAAGATCCTTTTTGATAATCTCATGACCAAAATCCCTTAACG
TGAGTTTTCGTTCCACTGAGCGTCAGACCCCGTAGAAAAGATCAAAGGATCTTCTTGAGATCCTTTTTTTCTGCG
CGTAATCTGCTGCTTGCAAACAAAAAAACCACCGCTACCAGCGGTGGTTTGTTTGCCGGATCAAGAGCTACCAAC
TCTTTTTCCGAAGGTAACTGGCTTCAGCAGAGCGCAGATACCAAATACTGTTCTTCTAGTGTAGCCGTAGTTAGG
CCACCACTTCAAGAACTCTGTAGCACCGCCTACATACCTCGCTCTGCTAATCCTGTTACCAGTGGCTGCTGCCAG
TGGCGATAAGTCGTGTCTTACCGGGTTGGACTCAAGACGATAGTTACCGGATAAGGCGCAGCGGTCGGGCTGAAC
GGGGGGTTCGTGCACACAGCCCAGCTTGGAGCGAACGACCTACACCGAACTGAGATACCTACAGCGTGAGCTATG
AGAAAGCGCCACGCTTCCCGAAGGGAGAAAGGCGGACAGGTATCCGGTAAGCGGCAGGGTCGGAACAGGAGAGCG
CACGAGGGAGCTTCCAGGGGGAAACGCCTGGTATCTTTATAGTCCTGTCGGGTTTCGCCACCTCTGACTTGAGCG
TCGATTTTTGTGATGCTCGTCAGGGGGGCGGAGCCTATGGAAAAACGCCAGCAACGCGGCCTTTTTACGGTTCCT
GGCCTTTTGCTGGCCTTTTGCTCACATGTTCTTTCCTGCGTTATCCCCTGATTCTGTGGATAACCGTATTACCGC
CTTTGAGTGAGCTGATACCGCTCGCCGCAGCCGAACGACCGAGCGCAGCGAGTCAGTGAGCGAGGAAGCGGAAGA
GCGCCCAATACGCAAACCGCCTCTCCCCGCGCGTTGGCCGATTCATTAATGCAGGGTCGCCGGCTTCGTCGGCGA
CGCCGTCCTGCTCCCCGGCACCGTCGACGTCGACTCCGGCGACACGGCCCGGACCGCCTTGGGCACGGTGCCCCT
CGCCGCTCCGCCCGGCGGCCTGCGCGCCGGCACGGTCCTGCTCCGTCCGGAACAACTCCGCCTCACCGGCGAGGA
CTCCGTCGGCGCGGTCAGGGCAACGGTCACCGACGTCTCGTACTACGGCCACGACGGAGCGGGCGGAACCGTGCT
CTGACCTGCGGCCCGAGTTTCGTCACGTGACGGAATGGAAGGCTGCTGCATTTCGTCACGTGACGTATCTCGGCG
AGCGACTGCCGACGCCACGGCGGACACGATCGCCTCGCGCTGGCGCCGGGCCTCGTACGCCCGCTGGCGGCAGGA
GCGGCGGCAGTAGTCCCGGCTCCGGCCGACGCCGGATTGCTTGATCTCCGAGCCGCACCAGGCGCAGAGCTTCGC
GCCGTCGGCGTCCCTGGGGGTGGTGGTGCTCATGGCCGACGACCGTACGCGGCACGTCTCGTAGCGAGGCGAGTC
GGGCGCGAGGTACCGCCTGCACGAAGTGCCGGCGGGGCCGACCCCGGGCGAGTAATCCCAGGATTACTCCCGCGG
CTTCGACCCCGGCCGCCGTCGCCGCGTACGTCACCGACCCCCGCCGTACGTCACCGGGATGACGTACGGCGGGGG
GGAGCGAGTTAGTGCGAAGTGGGCCCACTTGCGAGCCGGGCGATGTGCCGGGCGGCCCGCTCCTGGCGGTCGTCG
GCGTCGTCGTCCTGGTCGTCGTCCTGCTCTCGCCGTCGGCGTGCAGTTGCTTCCTCGCGGCGCTGGGCGAGGGCG
GCGAGCATGTCGGCGTACGCCTCGGCCACCTCCCCCGCCGTGAGCACCACCACTGTGTCGGCCGCGTCGGCCAGC
GCCAGGACCTCCCGCACCCGTTCGCCCACGGCCGCCGAATCCTCGTTGCCGTCCTTGCCTTCGGCGGCCCGGGTC
GCCTCGAGGTCGAGGGCGCGGCGGGTGACCGCGTGCCATCCGTCCTCGGTCACGGCGACCCCGGCCCGCAGCTCC
CCGCCGTCGGCGTCGGCCGCCAGGAGCAGATCGAGGTCGTCGGCCTCGGTGTCGCCGCCGTCGAGCCCGAGCATC
TGCCGCAGGTAGCGGGTCCATTCGATGGCCCGGCGTCCCCGGGTTGCCCGCTCGTACTCGTGCCAGCGCGAGAGG
TTCCACTCCAGCGAGCCGACCCCGGCGGCGTCGTCCTCGGTCATGCCGCCGGTCAGGTCCCCGATCCGTCCGAGG
AGTTCGAACGGGGCGACGTTCCCGCCGGTCGCCGTCTTGAGGTCGGCGCGGGCGAGTTCGAGGGCGGGCGCCTTC
CCGTCCTGGGTCTTGGCGATGTACTCGGCGAGGTCGTTGGCGTCGCGCTCGGTCTCCAGCCGCTTGAAGTCGACG

CCGTGCCGGTCGTCGGGCGTGAAGGCGGGGTTGACCTTGCGCAGGGCGGCGGTCCACACGGACCGCCAGTGCCCC
TGCCACTCGTCGAGCGCGGCGCCGGTCGGCTCGAAGGTGGCGACGATCTGCTTCGCGGACCGCTCCCCCTCGGTC
CGGCCGCCGACCAGGACGATCGCGTGGATGTGCGGGTGCCAGCCGTTGATCTGCCCCACGGTGACTTCGGTCGCG
CGGATCATGCCGACGTACCCGATCCGGTCTCGGATGCCCTCGCGGTCGGCGGCCCGGTGCCCGTCCTTGGCCCGG
CGTCCGGCCCACGTGCCGCCCGTGATCAGTCGCTGGTAGGCGCCCGGCCGCCGGGGGCTGTCCGGCGTCTTCCGG
GTGCCCTGGAGGGCGTCCATGAGGTCCGCGAGCCGGTCCGTGTGCCCATGGCGGGCCGTGAAGGTGACCAGGTAG
GCGGTCCCCCCGCGCTTGATCCACTCGACCACGGCGGCGGTGATCTCCTCGGCCCGCTTGTGCCGGATCGTGGCG
GCGCAGACCGGGCAGAGCCAGATCCGCCCGCACCGCATCAGGCCCAGGACCACGGACGTTCCGGCCGCCGTCTGG
GCGACGATTACGCCGGAGGCAGGGTCCATCAGGGCGCGGCCGCAGCCCTTGCACGCGGCGTCCCCGCTGATCCGC
CACAGCGTCCGGCGGCGGCTGTACCGGGCGGCTTTCCGCAGTCGGGCAGCGTCGCTCCGCGACGTGCTTCCTACT
TCCGAGAGGCTGTCGCCTCTCGGGCTCTCCCCATCCACCCCGTCCGGAGAAACCGCAGGTCGGAGGGGTGCGGGA
AACTCTGTTGTTTCTTTCCCAAGGTGTTCGCTTTTGCCTCGGGCGGCATCTCGCGTCACACGCGCGATCGCCCGC
TTCGCTGCCATCCGGCAGCGGTCTGAGCAGTAGATACGCGGCCGTTTGCCCGGTGTGTGGGCAATTGCGGTCCCG
CAGTGGCAGCGGGGCCCGGCGGGCCGATCTGGCAATGCCTCGGCATCGCTCCGTACTCTGGGCACGAGCAACGTT
CCTGTCTCGCCCGGCTAAGGGGCGCGAGTCTGGGAGCGGACGGGTCGGAGGTGCGAAGTCCGGCCCGTTGCTCTT
TGGTCTGGTGGGAATCCTGGCACCAATCGGGCCAGAGGTTCCCTCCGCCACTCCCGACGCCCCTTGGGGCTGGTG
TGACTTGGAGGGCCGAAGAGAGCCCCGCCGGTGATCCGGCGGGGCTTTGACGTGCGGTCAGTGCGTGTGTCGGCG
AGCGATGGCCACGAGGCCCTGGAAGCCGAGCGGTCCGGCGAAGTCGGCCCAGTCGCAACCGGGCTCAGCGCAGTG
GGCGGACCAGCCACCGCCGTTGGGGTCCTGGACCAGGTTCACGGTCCCCTCGGTCAGGCGTCCGTCGAAGTCGGT
CATGGTCGGTCTCCTGGTGGGTGGGGGCGGGGCGCCAGCACGAAGTGCCGGCGCCCCGCGGGGGTTGGTCGGGTC
AGGCGCCGAACCGGCGGGCGGCGGCGGCGACCAGGCCGTCGGCGGCGGCCATGGCGCGGTCGCGGTCGGTGGTGA
GGGCGGTGCGGTCGGCGGCGGCCCAGTCCTGCTCTCACGCAACGTCTACGTGGACGCTCAGGGCGACACGATCGA
GGTCGGGGAGTCCGTCGCTCCGGCGGGCCGCTGGCGCGTCCACCGGGACTGATCAAGGCAAATACTTCATATATG
CGGGGATCGACCGCGCGGGTCCCGGACGGGGAAGAGCGGGGAGCTTTGCCAGAGAGCGACGACTTCCCCTTGCGT
TGGTGATTGCCGGTCAGGGCAGCCATCCGCCATCGTCGCGTAGGGTGTCACACCCCAGGAATCGCGTCACTGAAC
ACAGCAGCCGGTAGGACGACCATGACTGAGTTGGACACCATCGCAAATCCGTCCGATCCCGCGGTGCAGCGGATC
ATCGATGTCACCAAGCCGTCGCGATCCAACATAAAGACAACGTTGATCGAGGACGTCGAGCCCCTCATGCACAGC
ATCGCGGCCGGGGTGGAGTTCATCGAGGTCTACGGCAGCGACAGCAGTCCTTTTCCATCTGAGTTGCTGGATCTG
TGCGGGCGGCAGAACATACCGGTCCGCCTCATCGACTCCTCGATCGTCAACCAGTTGTTCAAGGGGGAGCGGAAG
GCCAAGACATTCGGCATCGCCCGCGTCCCTCGCCCGGCCAGGTTCGGCGATATCGCGAGCCGGCGTGGGGACGTC
GTCGTTCTCGACGGGGTGAAGATCGTCGGGAACATCGGCGCGATAGTACGCACGTCGCTCGCGCTCGGAGCGTCG
GGGATCATCCTGGTCGACAGTGACATCACCAGCATCGCGGACCGGCGTCTCCAAAGGGCCAGCCGAGGTTACGTC
TTCTCCCTTCCCGTCGTTCTCTCCGGTCGCGAGGAGGCCATCGCCTTCATTCGGGACAGCGGTATGCAGCTGATG
ACGCTCAAGGCGGATGGCGACATTTCCGTGAAGGAACTCGGGGACAATCCGGATCGGCTGGCCTTGCTGTTCGGC
AGCGAAAAGGGTGGGCCTTCCGACCTGTTCGAGGAGGCGTCTTCCGCCTCGGTTTCCATCCCCATGATGAGCCAG
ACCGAGTCTCTCAACGTTTCCGTTTCCCTCGGAATCGCGCTGCACGAGAGGATCGACAGGAATCTCGCGGCCAAC
CGATAAGCGCCTCTGTTCCTCGGACGCTCGGTTCCTCGACCTCGATTCGTCAGTGATGATCACCCCCGACAGCGG
ATCAAGGGGTTTGCGGGTCCCGGTCGGCGCCGGGCGGGGGAGGCAGGAGCCGCCGACGCTGCCTCTGGGACGGGC
CGGACGGCAGGGGGACCGGCGGCCGGGCGAGCCTGGCGAAAGGGGGATGTGCTGCAAGGCGATTAAGTTGGGTAA
CGCCAGGGTTTTCCCAGTCACGACGTTGTAAAACGACGGCCAGTGAATTAGCTTGCATGCCGGAACTCACCCGTC
GTCGCGCGCTCGGCGCCGCAGCCGTCGTCGCCGCCGGTGTCCCGCTGGTCGCCCTTCCCGCCGCCCGCGCGGACG
ATCGGGGGCACCACACCCCCGAGGTCCCCGGGAACCCGGCCGCGTCCGGCGCCCCCGCCGCCTTCGACGAGATCT
CCCGTTCCATCCGTTCCCGCTGGGGGCCGTCCCCGACGACGAGGAAGGTCGGGGAGAGGTCGTCCAGGAGTTCGG
CCAGGCGGAGCAGGTCCGGCCAGCCCTTCTCGTGGGCGACCCGGCCGATGTAGCCGATGATCGGCCGGCGGCCGG
TCAGGCCGTGCCGCTGCGCGAACGCCGCCGCCTCGGCGGGTGTGGCCGGGGTAATGTCGACCGCGTCCGGGTTGA
CCACGATGGCGGACCGGTCCAGGCCGGTCGCGGCGCTCACCACGTCGGCGGTGCGGGCGGTCAGGGTGCTGACCC
GGGCGGCGGAGCGCAGCGCCTGTCGTTCCGCCTGGGTGACCAGGCGGTGCGCCATCGCGTCCACCGCCGACATCG
GCTGGTAGACGGAGAGCCGGGAGCAGTGCAGGGTGAGCACGTACGGCACGCCCAGGATGCGGGCGGCGATCCGCC
CGGCGACCAGTGGCCAGATCTGGCCGTCGGCGTGCACGTGGACGAGATCGGGACGCCAGTCACTGCGGCGCAGCC
GGAGGCATTCCCGGATGGTTCCGATCAACCAGGCCTGTCCGAGCCCGACCAGACCGGTGATCTCGGAACGGATCT
GGGGCAGGGACACGCGGGCGATACGAACGGTCAGCCCTGGCTCGATCTGCTTCGTCGGC
